# Identification and tracking of alloreactive T cell clones in Rhesus Macaques through the RM-scTCR-Seq platform

**DOI:** 10.1101/2021.10.29.466482

**Authors:** Ulrike Gerdemann, Ryan A. Fleming, James Kaminski, Connor McGuckin, Xianliang Rui, Jennifer F. Lane, Paula Keskula, Lorenzo Cagnin, Alex K. Shalek, Victor Tkachev, Leslie S. Kean

**Affiliations:** Division of Pediatric Hematology/Oncology, Boston Children’s Hospital; Department of Medical Oncology, Dana Farber Cancer Institute, and Harvard Medical School, Boston, MA; Broad Institute of MIT and Harvard, Cambridge, MA, USA; Institute for Medical Engineering and Science (IMES), Department of Chemistry, and Koch Institute for Integrative Cancer Research, MIT, Cambridge, MA, USA; Ragon Institute of MGH, MIT and Harvard, Cambridge, MA, USA

**Author notes:** UG, RF and JK contributed equally to this work and share first authorship. AKS, VT and LK contributed equally to this work and share last authorship. **Corresponding author:** Leslie S. Kean.

**Keywords:** TCR sequencing, single cell sequencing, alloreactive T cells, Rhesus Macaque, GvHD

## Abstract

T cell receptor clonotype tracking is a powerful tool for interrogating T cell mediated immune processes. New methods to pair a single cell’s transcriptional program with its T cell receptor (TCR) identity allow monitoring of T cell clonotype-specific transcriptional dynamics. While these technologies have been available for human and mouse T cells studies, they have not been developed for Rhesus Macaques, a critical translational organism for autoimmune diseases, vaccine development and transplantation. We describe a new pipeline, ‘RM-scTCR-Seq’, which, for the first time, enables RM specific single cell TCR amplification, reconstitution and pairing of RM TCR’s with their transcriptional profiles. We apply this method to a RM model of GVHD, and identify and track *in vitro* detected alloreactive clonotypes in GVHD target organs and explore their GVHD driven cytotoxic T cell signature. This novel, state-of-the-art platform fundamentally advances the utility of RM to study protective and pathogenic T cell responses.

## Introduction

Expression of the highly polymorphic (up to 10^18^ specificities)(1) T cell receptor (TCR) allows T cells to specifically recognize foreign and self-antigens presented in the context of the Major Histocompatibility Complex (MHC) by antigen presenting cells. This diversity is achieved by complex genetic recombination of the T cell alpha and beta chain during thymic T cell development, and results in a unique molecular barcode for each T cell. These unique TCR sequences enable studies of clonal- and diversity- dynamics in models of transplantation, infection, tumor immunology and vaccine development(2–4). More recently TCR tracking has been combined with high throughput single cell RNA sequencing (scRNA-Seq), enabling transcriptional profiling of distinct T cell clonotypes within and across samples (5, 6).

To date, clonotype tracking technologies have been developed for both mouse and human TCRs, facilitating studies of T cell clonal dynamics at an unprecedented level of molecular sensitivity.(7, 8) However, both murine and human models have drawbacks: The inbred nature of mice and the evolutionary distance between the murine and human immune systems can restrict the clinical applicability of murine results. Moreover, the study of human single cell TCR data continues to focus predominantly on the peripheral blood, being restricted by inherent challenges in accessing human tissues during health and disease. This can result in an incomplete picture of important immunological processes when using human samples.

To address these limitations, nonhuman primate (NHP) models (particularly using rhesus macaques (RM)) have been essential(9), and have enabled critical advances for some of the most pressing human needs, including the development of vaccines against SARS-CoV-2(10, 11), the study of HIV pathogenesis(12–14), and in advancing our mechanistic understanding of solid organ transplant(15) and hematopoietic stem cell transplant (HCT)(16–23). For HCT, NHP models have enabled the translation of several novel therapeutic strategies to the clinic for the prevention of graft-versus-host disease (GVHD), the major cause of transplant-related mortality after HCT(18, 21, 24).

The identification and tracking of TCR clonotypes are of critical importance for each of the clinical settings described above, in order to understand the molecular mechanisms driving immune protection or disease pathogenesis. For acute GVHD (aGVHD), dissecting the link between T cell clonal architecture and disease is particularly relevant, given that donor-derived alloreactive T cells are the main culprits in the aGVHD-mediated destruction of the skin, intestine, and liver^(25),(26)^. Accurate identification and characterization of alloreactive T cell clonotypes is essential to identifying the molecular drivers of aGVHD, and doing so in aGVHD-target tissues as well as the peripheral blood remains a major challenge. Until the work described in this manuscript, the identification and tracking of individual TCR clonotypes in NHP models (in particular, RM) has been limited by a lack of robust methodology for single cell TCR sequencing adapted to this essential pre-clinical model. The key limitations facing the field were the lack of effective RM-specific primers that could successfully amplify the TCR region in a highly multiplexed fashion, as well as the incomplete annotation of the RM genome throughout the TCR alpha and beta chain region, which created a major barrier to accurate TCR chain reconstruction.

We now describe the design and validation of RM-specific primers compatible with the human 10x Genomics single cell sequencing platform, which accurately amplifies the RM TCR alpha and beta regions. Amplified fragments aligned to our custom-assembled and annotated RM TCR genome permitting, for the first time, the *in vitro* and *in vivo* identification and tracking of RM derived T cell clonotypes. Pairing of *in vivo* identified alloreactive clonotypes with their transcriptional profile revealed a highly activated cytotoxic CD8 T cell signature during NHP aGVHD.

## Methods

### NHP

This study was conducted in strict accordance with (USDA) United States Department of Agriculture regulations and the recommendations in the Guide for the Care and Use of Laboratory Animals of the National Institutes of Health. It was approved by the Massachusetts General Hospital and Biomere Animal Care and Use Committees. T cells were obtained from healthy colony RM or from animals who underwent allogenic HCT in the setting of previously published studies(23).

### Mixed Lymphocyte Reaction (MLR)

RM PBMCs were isolated from whole blood by Ficoll gradient centrifugation, and then used for MLR assays either immediately, or after liquid nitrogen cryopreservation in 10% dimethyl sulfoxide (DMSO)/90% fetal bovine serum (FBS). At the time of the MLR, stimulator PBMCs were irradiated with 3500 cGy of (^137^Cs) radiation. Responder PBMCs were stained with cell trace violet (CTV, Invitrogen) as per manufacturer’s instructions. 2× 10^5^ T cell-enriched responder PBMCs along with an equal number of stimulator PBMCs were added to each well in a 96-well plate (Corning) in X-vivo-15 medium (BioWhitaker) supplemented with 10% FBS (Irvine Scietific) and incubated at 37°C for a total of 5 days. Cell culture media change was performed on day 3 of the culture. At the end of 5 days, cells were stained with an extracellular antibody for CD3 (clone SP34-2), CD20 (clone 2H7) and CD14 (clone M5E2, all antibodies from BD Bioscience) for 20min at 4°C, and high-, medium- and non-proliferating CD3+ T cells (identified based on the dilution of CTV) were sorted and processed for single cell sequencing using the Chromium Next GEM Single Cell 5’ Reagent Kit v1 with optimized RM primers as described below.

### NHP HCT, Necropsy and Tissue Processing

RM HCT, necropsy, and tissue processing were performed as previously described(23, 24). Single cell suspensions resulting from tissue processing were sorted on a live CD3+ and CD14/CD20-population and prepared for single cell sequencing.

### Primer design and PCR amplification

TCR alpha and beta primers were designed to perform in a nested PCR approach to optimize target enrichment of the respective constant regions. TRAC inner and outer primers were 297bp apart, and the beta chain inner and outer primers were 357bp apart. Primers were optimized to match the melting temperature of the 10x forward primers for annealing to a known sequence within the gel beads in emulsion (GEMs) according to the human 10x Genomics Chromium Single Cell V(D)J Reagent Kits User Guide for target enrichment (https://assets.ctfassets.net/an68im79xiti/31W4aZOJ8C2ipxzTrcQIWn/72c5c2bc3dd7784f44f5218ca345ce04/CG000086_ChromiumSingleCellV_D_J_ReagentKits_UG_RevM.pdf). Primer sequences are listed in Table1.

**Table 1:**
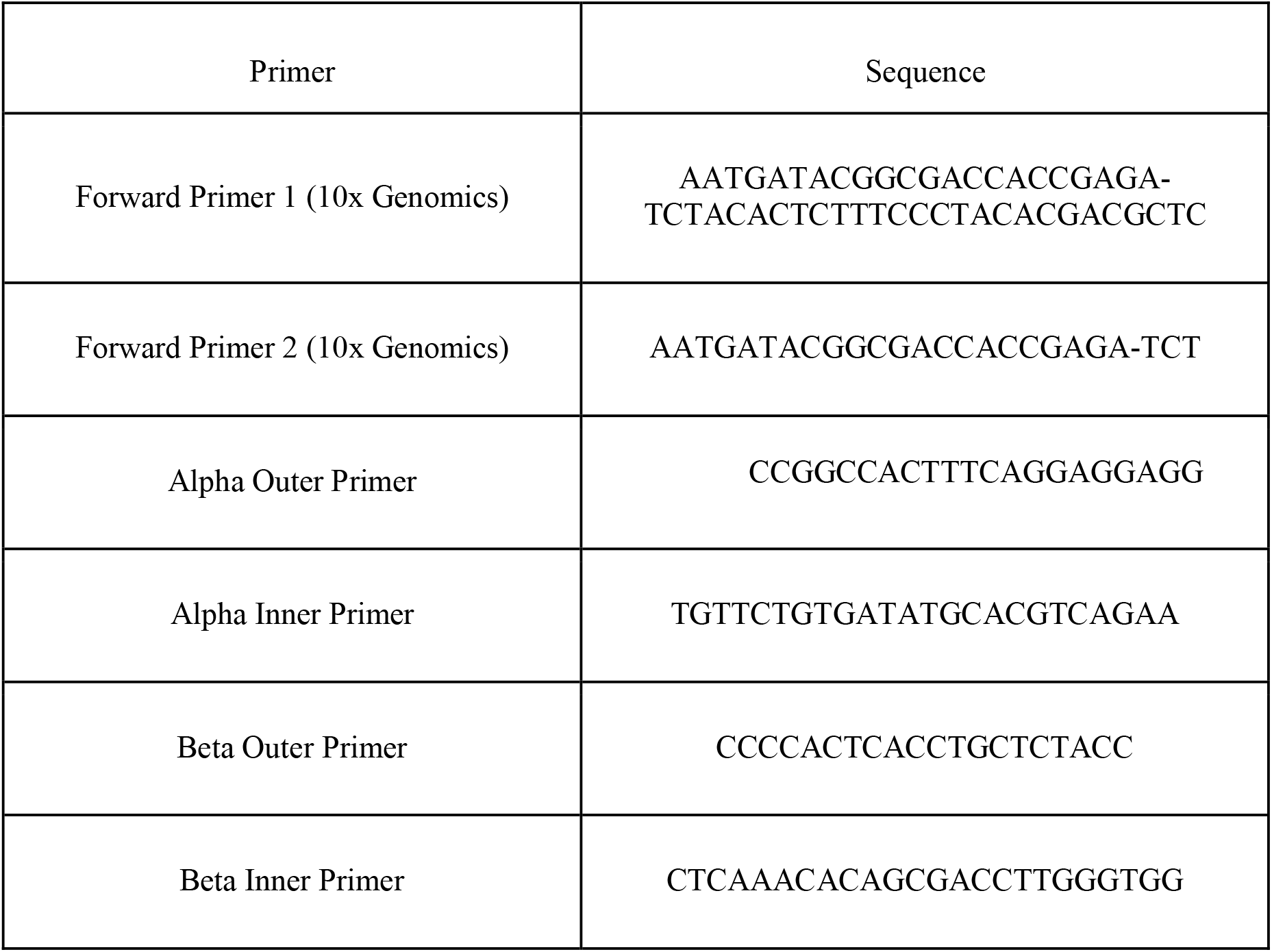
Primer sequences optimized for RM-scTCR-Seq.

PCR was performed with an initial denaturation temperature of 98°C for 45 seconds, followed by denaturation at 98°C for 20 seconds. Annealing was optimal at 67°C for 30 seconds with extension of 72°C for 1 min. A total of 20 PCR cycles were performed with a final extension step at 72°C for 1 min.

### GEX Alignment and TCR Reconstruction

Samples were aligned using the Cellranger multi pipeline (v6.0.2), with VDJ libraries listed as “vdj-t” to indicate T cell libraries. Samples were aggregated with the Cellranger aggr pipeline (v6.0.2) with normalize=none.

### GEX Filtering and QC

We used FASTQC to assess the quality of our sequencing data. After alignment of this data, we loaded the filtered_feature_bc_matrix.h5 file from Cellranger multi into Seurat and removed genes which were present in less than 5 cells. This resulted in a dataset with 38,147 cells by 16,163 genes. Following the recommendations in Lucken and Theis(51), we analyzed the total transcripts detected and total genes detected for each sample to determine appropriate thresholds and retained cells with 500 to 6,000 genes detected and 1,800 to 40,000 transcripts detected, and less than 5% of aligned reads originating from the mtDNA. We then applied SCTransform(52) to normalize the data, reduced the dimensionality of the dataset using PCA (retained 22 PC’s after check of ElbowPlot), clustered the data using Seurat’s FindClusters function with a resolution of 0.2, and subsetted the dataset to T cells by removing two clusters that were not enriched for the T cell genes CD3E, CD4, CD8A, or TRAC. This resulted in a dataset with 28,646 cells by 15,904 genes. We then repeated the above procedure from SCTransform to clustering using 23 PC’s.

We then limited the dataset to samples from the PBMC, the MLR experiments, and the GVHD target organs. We then removed genes present in less than 5 cells, resulting in a dataset with 23,618 cells by 15,195 genes. This dataset was normalized using the same pipeline previously described in this section, retaining 22 PC’s from the PCA.

### Identification of Clonotypes

Clonotypes used in the gene expression analysis were identified using cellranger’s default settings in “cellranger vdj” for the scTCR-Seq libraries.

As discussed in the Results section, the clonotypes used for the alpha chain and beta chain Morisita heatmaps (**Figure 2, panels E and F**) were identified by grouping together T cells on the basis of the CDR3 region in the alpha chain or beta chain. This is to provide a more stringent test of the specificity of clonotype calling, as Cellranger’s default settings do not allow for clonotypes to be shared across different donors. The Morisita index was calculated using the “vegan” package in R.

### Differential Expression (DE) Tests

DE tests were carried out as Wilcoxon tests in Seurat, using the default settings. Results with an adjusted p value <0.05 were considered significant.

### Scoring of Cells with Gene Signatures using VISION

We used VISION to apply gene signatures to the normalized expression data for the analyzed T cells. We used the C7 collection of immunological signatures from MSigDB, and ten signatures from the C2 curated collection that contained the key phrases “allo” or “graft” to identify potential alloreactivity signatures. To test for differences in signature enrichment between cells belonging to MLR+ and MLR-clonotypes within an organ, we used the Wilcoxon test.

## Results

### Efficient amplification of RM TCR alpha and beta chains

Immunological studies in RM have been performed for decades, and have significantly advanced our understanding of adaptive T cell immunity in health and disease. However, in-depth characterization of RM T cell clonality and associated T cell function on a single cell level have been hindered by a paucity of available methods, particularly the ability to efficiently amplify and reconstitute the RM alpha and beta TCR chains. Here we present a robust platform for RM TCR amplification leveraging the well-established human 5’ 10x Genomics single cell sequencing platform. To accomplish this, we optimized primer pairs, adapted to utilize a nested PCR approach analogous to that of the 10x human and mouse workflows, thereby maximizing the specificity of amplification, with primer sets targeting the constant regions of the RM alpha and beta chains (‘TRAC’ and ‘TRBC’, respectively). The primers targeting the beta chain were specifically designed to amplify both TRBC1 and TRBC2. Primer length, CG content and annealing temperatures were adjusted to be used with the forward primers supplied in the human 5’ 10x Genomics kit. Cycling conditions were modified from the original 10x protocol to allow optimal amplification of the targeted region in RM (**Table 1**). We have named this new resource ‘RM-scTCR-Seq’.

RM-scTCR-Seq primer annealed to the alpha and beta RM constant region and amplified transcripts covering the complete alpha and beta loci including the variable V and J regions for the alpha as well as V, D and J regions for the beta locus. (**Figure 1A**). Our assembled contigs extend past the V region, which likely represents unannotated UTR sequences. Such UTR sequences are annotated in the human and mouse VDJ references(27) and can be seen in the assembled contigs for 10x Genomics’ human and mouse sample datasets.

**Figure 1.**
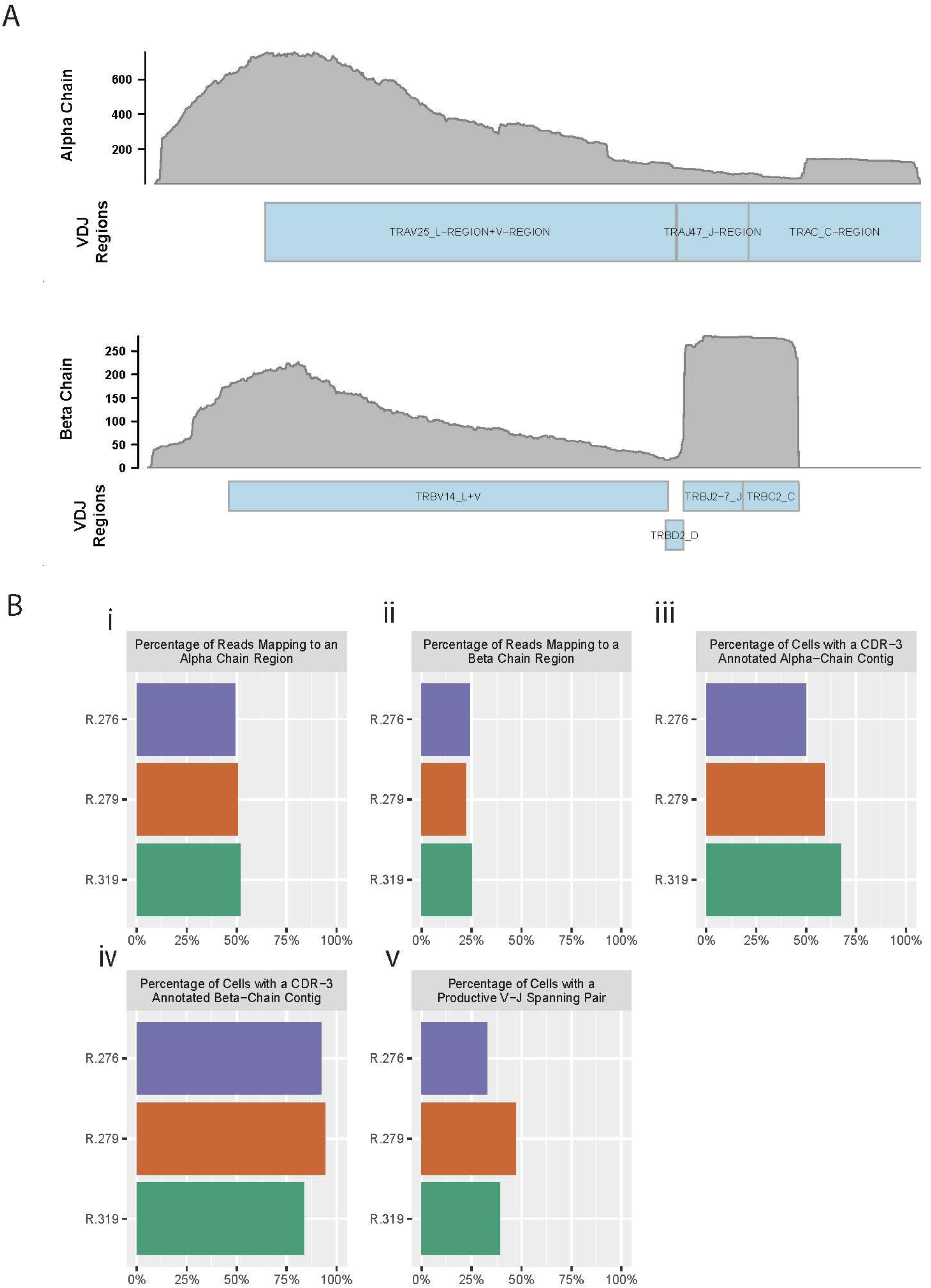
RM primers amplify reads from alpha and beta chains, and enable TCR reconstruction. **(A)** Pile up of reads on the alpha chain (top) and beta chain (bottom) from a reconstructed RM TCR. **(B)** Barplots of “Cellranger vdj” summary statistics for three RM PBMC T cell samples. (i) Percent of reads in each sample that map to the alpha chain of the TCR, (ii) Percent of reads in each sample that map to the beta chain of the TCR, (iii) Percent of cells (defined as droplets identified by Cellranger as containing a valid cell) with a reconstructed CDR3-annotated alpha chain contig, (iv) Percent of cells (defined as droplets identified by Cellranger as containing a valid cell) with a reconstructed CDR3-annotated beta chain contig, (v) Percent of cells with a productive (meaning no premature stop codons) alpha chain and a productive beta chain, both of which span the start of a V sequence to the end of a J sequence.

We assessed the quality of our primers using summary statistics from 10x’s ‘Cellranger’ software (a pipeline for alignment, quantification, and evaluation of 10x Genomics-generated single-cell libraries), and found that our primers consistently amplify high-quality reads from the TCR alpha and beta chains. For this assessment, we prepared peripheral blood mononuclear cells (PBMCs) from three representative RMs (animal IDs: R.276, R279, R.319) and used these PBMCs to perform RM-scTCR-Seq. We observed that 49-52% of the aligned reads mapped to the alpha chain and 22-25% of the reads aligned to the beta chain (**Figure 1B i and ii, Table S1**) in the three RMs, with >=73% of total reads mapping to any V(D)J gene (including to gamma and delta chain sequences).

In addition to demonstrating that the sequencing reads mapped to known TCR loci, we were also able to reconstruct the alpha and beta chains from these reads (**Figure 1B iii and iv, Table S1**). The Complementarity-determining region (CDR) 3 is the most variable of the CDR regions, and spans the V and J regions; thus, identification of this sequence is useful as a benchmark for reconstruction quality(28). As such, an ideal reconstruction of the alpha chain or beta chain would completely span the V-to-J region, and enable inference of the CDR3 sequence(29, 30). Using the preceding criteria as our standard for a high-quality alpha or beta TCR chain reconstruction, we observed that 50 to 67% of the 10X droplets identified as cells by Cellranger produced a full length, CDR3-annotated alpha chain (**Figure 1B iii, Table S1**). Inspecting the beta chain, we found that a high-quality beta chain could be constructed for 83% to 94% for these same cells (**Figure 1B iv, Table S1**). Full reconstruction of the TCR requires both an alpha chain and a beta chain that span the V and J regions, and ideally are productive (no premature stop codons). We identified both alpha and beta chains in 32 to 47% of cells examined (**Figure 1B v, Table S1**).

These data demonstrate, for the first-time, efficient amplification of the RM TCR alpha and beta region by utilizing optimized RM primer pairs compatible with the human 5’ 10x Genomics single cell sequencing platform. Amplified reads enabled high quality beta chain reconstruction in almost all, and reconstruction of full TCRs (alpha and beta chains) in a significant number of sequenced cells.

### RM-scTCR Seq enables tracking of T cell clonotypes in allogenic mixed lymphocyte reactions

Identification of T cell clonotypes using *in vitro* assays or *in vivo* disease models enables the evaluation of the clonal repertoire, clonal dynamics, and the tracking of specific T cells over time. To validate our single cell TCR sequencing and analysis pipeline, we first utilized an *in vitro* allo-proliferation assay: the mixed lymphocyte reaction (MLR)(31, 32). To perform these studies, PBMCs from 4 RM were irradiated with 3500 cGy and used to stimulate “responder” PBMCs from MHC-disparate RM. The responder PBMCs were labeled with cell trace violet (CTV), a cell dye which dilutes with proliferation. After five days of allo-stimulation, responder T cell were sorted into three populations: CTV High (non-proliferating T cells), CTV Mid (intermediate proliferation, 3-5 cell divisions) and CTV Low (highly proliferating T cells, demonstrating > 7 cell divisions by flow cytometry), (**Figure 2A-B**). We applied the RM-scTCR-Seq pipeline to analyze TCR diversity in each of these fractions, and assessed this diversity using the Shannon index(33, 34). As demonstrated in **Figure 2C**, the highest proliferating population displayed the lowest Shannon diversity (CTV Low Shannon index = 7.52 – 8.40), indicating highest clonality, while diversity increased with the less proliferating cells (CTV Mid Shannon index = 7.61 – 8.93 and CTV High Shannon index = 7.94 – 9.09). Using the RM-scTCR-Seq pipeline, we were also able to track individual clonotypes throughout the three MLR peaks (**Figure 2D**).

**Figure 2.**
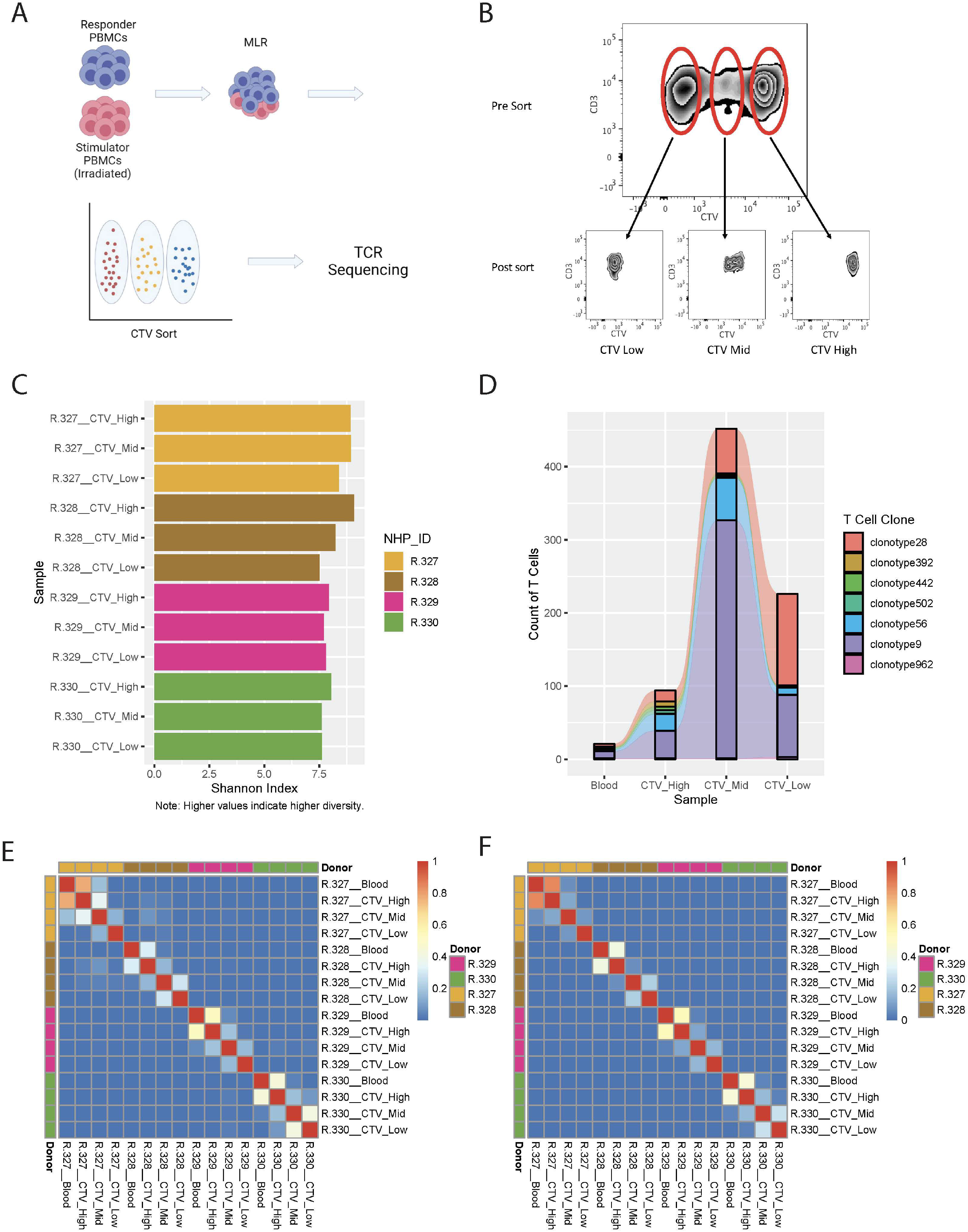
Application of RM primers to an MLR experiment demonstrate the ability to track T cell clonotypes over across samples with high accuracy. **(A)** Experimental schema for MLR experiments. **(B)** CTV staining and sorting strategy for one representative RM. **(C)** Shannon Index for each of the sorted samples from four individual MLRs. **(D)** Representative graph of T cell clonotypes detected in the peripheral blood (Pre-MLR) and from all three sorted samples (CTV-High, CTV-Mid, CTV-Low) for R.330. **(E)** Heatmap of Morisita Index for all MLR and peripheral blood samples, using the CDR3 region of the alpha chain to group cells into clonotypes. **(F)** Heatmap of Morisita Index for all MLR and peripheral blood samples, using the CDR3 region of the beta chain to group cells into clonotypes.

To further assess our ability to accurately track T cell clonotypes *in vitro,* and to quantify TCR similarities/differences between T cells isolated from different animals, we calculated the Morisita Index (‘MI’, measuring the similarity of two repertoires, or, the reproducibility of a single repertoire between different samples)(34, 35) between all pairs of samples from the MLR experiment. We analyzed these data using the CDR3 regions of the beta chain (**Figure 2E**) and the alpha chain **(Figure 2F**).

As has been previously demonstrated, the vast majority of TCR clonotypes in our assay were private to the individual animal, with low levels of similarity when comparing TCR clonotypes between different animals (MI < 0.08, indicating minimal overlap in the TCR repertoire, **Figure 2E,F**). As expected, the highest level of similarity was observed between responder PBMCs in the ‘Blood’ sample (T cells isolated from peripheral blood pre MLR) and T cells in the non-proliferating population (‘CTV high’). Importantly we were also able to identify clonotypes that existed in both the Blood sample and the high-proliferating population (**Figure 2E,F**), substantiating the ability of RM-scTCR-Seq to track clonotypes as they proliferated *in vitro*. The low Morisita indices for comparison of samples between different donors, using either alpha chain MI (MI < 0.08) or beta chain MI (MI <0.001), demonstrate the specificity of the RM-scTCR-Seq approach (**Figure 2E,F)** and validate that clonotype tracking can be performed by utilizing the TCR alpha or beta chains.

### *In vitro* predicted alloreactive T cell clonotypes exhibit an *in vivo* aGVHD expression program in the spleen and liver

We next performed *in vivo* clonotype tracking in HCT recipients who developed aGVHD, in order to trace specific TCRs previously identified as allo-proliferating in an *in vitro* MLR. To accomplish this, we first performed MLR assays using donor and recipient PBMCs from transplants in which recipients subsequently developed severe aGVHD.(23) For these assays, responder PBMCs were from donor animals R.276, R.279, R.319, and stimulator PBMCs were from recipient animals R.227, R.276 and R.312, respectively(23). Terminal analysis of spleen and liver infiltrating donor T cells was performed 8 days after HCT, during active aGvHD(23). Sorted T cells were analyzed with 5’scRNA-Seq using the 10X platform, and clonotype-tracking was performed using the RM-scTCR-Seq pipeline.

We first assessed our ability to track clonotypes across samples by applying the MI to all of the scTCR-Seq libraries (**Figure 3A**). As expected, in all three animals, measurable TCR clonal overlap could be detected in T cells sorted from recipient liver and spleen, consistent with infiltration of activated T cell clonotypes into both of these organs. Overlap of alloreactive (highly proliferating) clonotypes from the MLR with clonotypes derived from both the spleen and the liver was also evident **(Figure 3A)**. Next, we confirmed that the majority of cells with trackable TCRs by RM-scTCR-Seq cells also had a corresponding barcode in the scRNA-Seq GEX library after quality control and filtering. We found that at least 75% of the cells with a valid scTCR-Seq library (assessed by Cellranger) in each sample also had a valid corresponding scRNA-Seq barcode **(Figure 3B)** confirming the efficiency of the assay. After alignment, filtering based on quality metrics, and dimensionality reduction, we plotted a UMAP of the analyzed cells (n= 3 RM, 23,618 cells) separated according to each transplant recipient **(Figure 3C)**.

**Figure 3.**
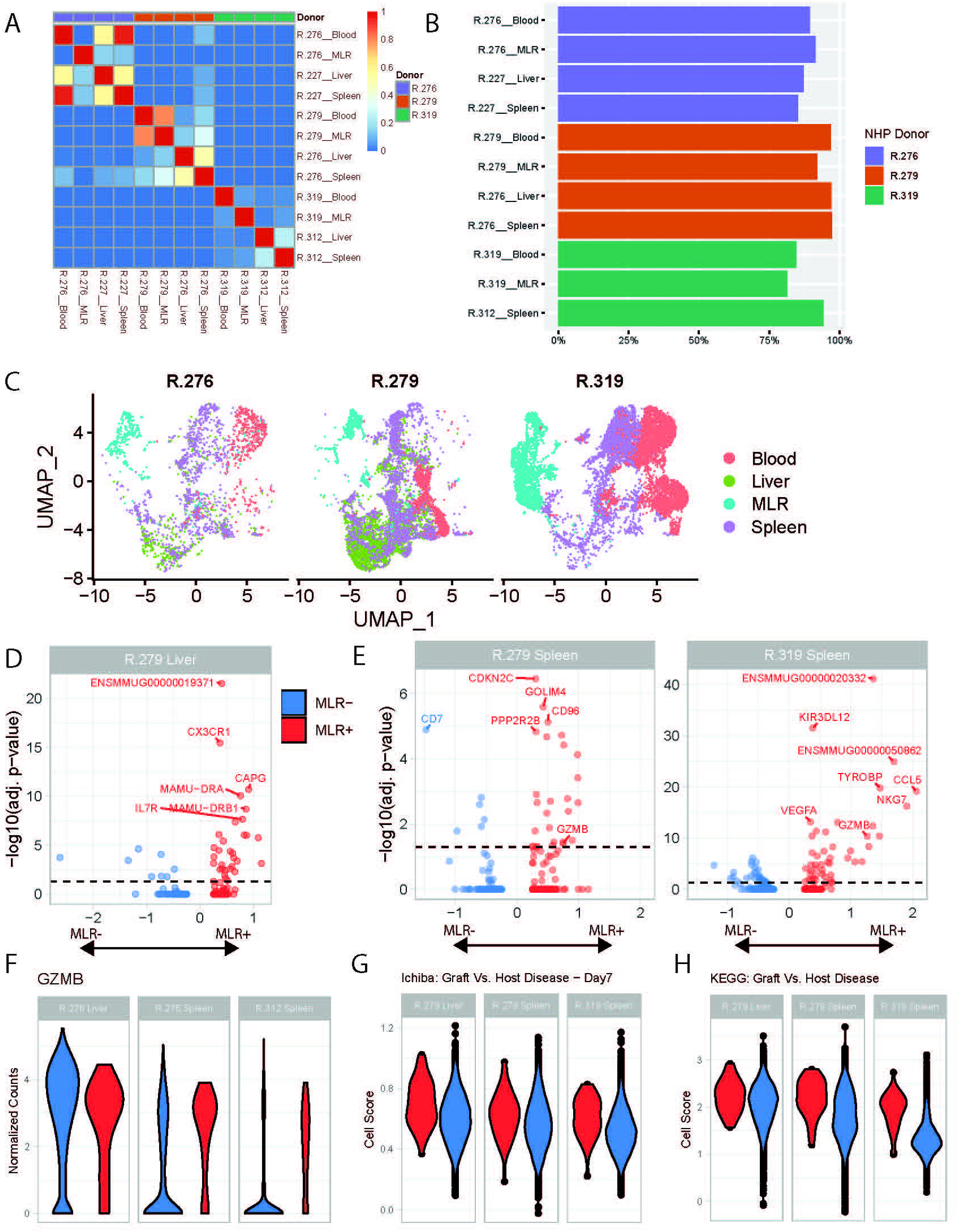
Allo-proliferative T cell clonotypes identified in an MLR exhibit aGVHD-specific gene expression programs in the spleen and the liver. **(A)** Heatmap of the Morisita Index for all analyzed samples. **(B)** Percentage of cells assigned to a specific T cell clonotype using RM-scTCR-Seq that also had high-quality gene expression data using the 10X gene expression platform. **(C)** UMAP of cells from each donor NHP, colored by sample type. Blood = T cells sorted from the blood. MLR= T cells subjected to a MLR. Spleen = T cells purified from the spleen during severe aGVHD. Liver = T cells purified from the liver during severe aGVHD. **(D)** Volcano plots of differential expression between MLR+ and MLR− detected clonotypes within one liver sample. **(E)** Volcano plots of differential expression between MLR+ and MLR− detected clonotypes within two spleen samples. **(F)** Violin plot of Gzmb expression between MLR+ and MLR− cells within two spleen samples (from R.279 and R.319) and one liver sample (from R.279). **(G)** Violin plot of T cells scored with the Ichiba Graft Versus Host Disease signature(38), colored by the detection of a clonotype in MLR+ (red) or MLR− (blue) samples. P <0.05 for the comparisons in the liver, and both spleen samples. **(H)** Violin plot of T cells scored with the KEGG Graft Versus Host Disease signature (39) colored by the detection of a clonotype in MLR+ (red) or MLR− (blue) samples. P <0.001 for the comparisons in the liver, and both spleen samples.

We then characterized cells isolated from the spleen and liver by whether they could be matched with an alloproliferating MLR clonotype (termed ‘MLR+’) or with a non-allo-proliferating clonotype (termed ‘MLR−’). Using these definitions of MLR+ and MLR−, we conducted a gene expression test of MLR+ versus MLR− T cells within two spleen samples and one liver sample with large numbers of MLR clonotypes, to identify differentially expressed (DE) genes. The volcano plots in **Figures 3D-E** demonstrate these DE genes, and highlight the top 5 annotated genes in each comparison (top genes identified only with an Ensembl ID are also identified).This analysis identified several known markers of alloreactivity in each sample. *Gzmb*, a well-recognized aGVHD-associated gene(23, 36) was identified as a DE gene in both spleen samples. **(Table S2)**. We further interrogated the expression of *Gzmb* at the level of each individual cell, to understand its variability within the MLR+ and MLR− populations. As expected, a violin plot of *Gzmb* **(Figure 3F)** demonstrated that the spleen MLR+ samples had higher, less variable expression of *Gzmb* compared to the spleen MLR− samples, which demonstrated a wider distribution of expression, with a larger proportion of low-expressing cells. To further evaluate gene expression in the MLR+ and MLR− cells, we used VISION(37) to score each cell using a set of previously curated immunological and alloreactivity signatures. As shown in **Figure 3G-H**, MLR+ cells from the liver and spleen enriched for the Ichiba: Graft versus Host Disease gene signature compared to MLR− cells (Figure 3G, p < 0.05)(38) as well as for the KEGG Graft versus host disease signature (Figure 3H, p <0.01)(39).

These results demonstrate the feasibility of combining RM-scTCR-Seq and scRNA-Seq to identify and interrogate alloreactive T cell clonotypes in aGVHD target organs in RM, providing proof-of-concept of the *in vivo* utility of the RM-scTCR-Seq pipeline.

## Discussion

TCR repertoire analysis has become a fundamental tool to understand the immune responses to infections and vaccines, as well as the immunopathogenesis of rejection after solid organ transplantation, and GVHD after HCT(2, 40–43). In murine and human studies, the introduction of VDJ target enrichment combined with single cell sequencing platforms has enabled the interrogation of transcriptional profiling of single T cells at an unprecedented level of accuracy(29, 30). However, paired single cell T cell receptor and transcriptional studies in RM have been impeded by the lack of a validated platform to amplify and reconstitute alpha and beta TCR sequences(44–46). This study demonstrates successful identification and tracking of RM single T cell clonotypes by utilizing customized primers compatible with the human 5’ 10x Genomics single cell sequencing platform, and by significantly improving alignment of RM sequencing results by re-constructing a custom genome of the RM TCR alpha and beta region.

Prior to the work described herein, two major limitations significantly hindered the development of T cell clonotype tracking in RM. They were: (1) The lack of optimized, single cell sequencing-compatible primers covering both the TCR alpha and beta region for RM; (2) The poor genomic annotation of the alpha and beta loci in RM, which resulted in inadequate reconstruction of amplified VDJ PCR reads. To address both of these technical barriers, we have designed a robust set of RM-specific TCR primer pairs which anneal to the constant region of the alpha and beta TCR loci, and which are compatible with the commonly used single cell sequencing platform from 10x Genomics. Optimizing PCR cycling and temperature conditions resulted in efficient amplification of the alpha and beta regions of the RM TCR and generated productive TCR reconstructions. Additionally, we created an updated reference for the TCR alpha and beta chain region, enhancing our ability to annotate the assembled RM TCR’s.

We demonstrated amplification of highly specific TCR regions with >73% of total reads mapping to any V(D)J gene. We note that this mapping efficiency is somewhat less than the ~91% of reads mapping to the human VDJ reference, in a human dataset provided by 10x (www.10xgenomics.com/resourses/dataset). The difference in the alignment rate may be explained by differing levels of annotation for these organisms: 10x provides human references which have been refined over time, but this resource does not exist for RM, which necessitated our creation of a new reference (see Methods) by combining elements of the Ensembl and IMGT references, and then carefully filtering out sequences identified by 10x’s enclone software that contained errors (such as premature stop codons, see Methods). As annotation of the RM TCR improves, we anticipate some gains in read mapping. As mentioned earlier, we have noticed that UTR regions are annotated in the human and mouse VDJ references, and we expect that addition of these sequences to the RM reference will improve alignment rates.

Sufficient alignment was also demonstrated by the ability of the RM-scTCR-Seq pipeline to detect allo-proliferating clonotypes using an *in vivo* RM model of aGVHD. While bulk TCR-Seq is well-suited for providing a comprehensive profile of the diversity of T cell clonotypes in a given sample due to its ability to profile large numbers of cells(44, 46), scTCR-Seq is unique in its ability to be paired with other single-cell technologies, allowing users to tag each T cell with its clonal identity, and measure each cell’s gene expression profile as performed here(6). These clonotypes can also be interrogated for their protein expression or epigenetic profiles, or other features of interest (47, 48). Most importantly, these individual clonotypes can be tracked *in vivo*, to better map the clonal architecture of protective versus pathogenic T cell responses, in target tissues that are far more accessible in NHP than in patient samples. In this study utilizing RM-scTCR-Seq, we were able to detect allo-proliferative clonotypes in *in vitro* MLR experiments and track their *in vivo* alloreactive and cytotoxic potential by transcriptional analysis of tissue-infiltrating T cells. We expect that future RM-scTCR-Seq studies from GVHD target tissues including the GI tract and skin will provide further insights into mechanisms controlling T cell alloreactivity, and enhance our ability to identify GVHD-specific targetable pathways.

Given the central importance of NHP models to our understanding of protective and pathogenic immunity, and to the development of novel immune-based therapeutics, the application of RM-scTCR-Seq is expected to have wide application, and further advance the ability of NHP to model human disease.

## 2 Author Contributions

Conceptualization: UG, RAF, VT, LK

Methodology: UG, RAF, JK, VT

Investigation: RAF, UG, JK, XR, LC, JFL

Visualization: JK, UG

Supervision: LK, AS, PK, LC

Writing—original draft: UG, JK, LK

Writing—review & editing: UG, JK, RAF, XR, JFL, PK, LC, VT, AKS, LK

