## Supplement for "Identification and tracking of alloreactive T cell clones in Rhesus Macaques through the RM-scTCR-Seq platform"

Supplementary Table S2: Differential Expression MLR+ vs MLR-

| Test | Gene | Adjusted P Value | P Value | Average Log2 FC | MLR+ Cells Expressing Gene | Cells Expressing Gene |
|---|---|---|---|---|---|---|
| R.279 Liver MLR+ vs MLR- | ENSMMUG00000019371 | 0.000 | 0.000 | 0.41 | 0.362 | 0.048 |
| R.279 Liver MLR+ vs MLR- | CX3CR1 | 0.000 | 0.000 | 0.37 | 0.276 | 0.037 |
| R.279 Liver MLR+ vs MLR- | CAPG | 0.000 | 0.000 | 0.91 | 0.638 | 0.218 |
| R.279 Liver MLR+ vs MLR- | MAMU-DRA | 0.000 | 0.000 | 0.76 | 0.345 | 0.076 |
| R.279 Liver MLR+ vs MLR- | MAMU-DRB1 | 0.000 | 0.000 | 0.86 | 0.534 | 0.179 |
| R.279 Liver MLR+ vs MLR- | IL7R | 0.000 | 0.000 | 0.79 | 0.586 | 0.205 |
| R.279 Liver MLR+ vs MLR- | DHRS7 | 0.000 | 0.000 | 0.66 | 0.621 | 0.252 |
| R.279 Liver MLR+ vs MLR- | PPP2R2B | 0.000 | 0.000 | 0.35 | 0.259 | 0.056 |
| R.279 Liver MLR+ vs MLR- | S100A10 | 0.000 | 0.000 | 0.83 | 0.983 | 0.866 |
| R.279 Liver MLR+ vs MLR- | BCL2A1 | 0.000 | 0.000 | 0.87 | 0.879 | 0.613 |
| R.279 Liver MLR+ vs MLR- | ANXA1 | 0.000 | 0.000 | 1.09 | 0.776 | 0.475 |
| R.279 Liver MLR+ vs MLR- | ENSMMUG00000060606 | 0.000 | 0.000 | 0.43 | 0.207 | 0.04 |
| R.279 Liver MLR+ vs MLR- | CDKN2B | 0.000 | 0.000 | 0.26 | 0.259 | 0.061 |
| R.279 Liver MLR+ vs MLR- | LAG3 | 0.000 | 0.000 | -1.16 | 0.207 | 0.575 |
| R.279 Liver MLR+ vs MLR- | B2M | 0.000 | 0.000 | 0.35 | 1 | 0.999 |
| R.279 Liver MLR+ vs MLR- | CD52 | 0.000 | 0.000 | 0.57 | 0.983 | 0.945 |
| R.279 Liver MLR+ vs MLR- | ENSMMUG00000061119 | 0.000 | 0.000 | 0.61 | 0.241 | 0.059 |
| R.279 Liver MLR+ vs MLR- | SPOCK2 | 0.000 | 0.000 | -0.73 | 0.086 | 0.527 |
| R.279 Liver MLR+ vs MLR- | ENSMMUG00000056183 | 0.000 | 0.000 | 0.26 | 0.172 | 0.034 |
| R.279 Liver MLR+ vs MLR- | GZMA | 0.000 | 0.000 | -2.62 | 0.569 | 0.752 |
| R.279 Liver MLR+ vs MLR- | KLRD1 | 0.000 | 0.000 | 0.63 | 0.741 | 0.409 |
| R.279 Liver MLR+ vs MLR- | PRF1 | 0.000 | 0.000 | -1.34 | 0.517 | 0.728 |
| R.279 Liver MLR+ vs MLR- | EFHD2 | 0.000 | 0.000 | 0.69 | 0.776 | 0.5 |
| R.279 Liver MLR+ vs MLR- | MATK | 0.001 | 0.000 | 0.25 | 0.293 | 0.086 |
| R.279 Liver MLR+ vs MLR- | CD74 | 0.001 | 0.000 | 1.15 | 0.81 | 0.507 |
| R.279 Liver MLR+ vs MLR- | ENSMMUG00000060287 | 0.001 | 0.000 | 0.41 | 0.983 | 0.989 |
| R.279 Liver MLR+ vs MLR- | SERPINA1 | 0.002 | 0.000 | 0.60 | 0.345 | 0.122 |
| R.279 Liver MLR+ vs MLR- | ENSMMUG00000064139 | 0.003 | 0.000 | 0.42 | 1 | 0.997 |
| R.279 Liver MLR+ vs MLR- | PLIN2 | 0.003 | 0.000 | 0.56 | 0.845 | 0.614 |
| R.279 Liver MLR+ vs MLR- | RPL12 | 0.003 | 0.000 | -0.48 | 0.983 | 0.996 |
| R.279 Liver MLR+ vs MLR- | CD96 | 0.003 | 0.000 | 0.37 | 0.466 | 0.192 |
| R.279 Liver MLR+ vs MLR- | TESC | 0.004 | 0.000 | 0.47 | 0.69 | 0.393 |
| R.279 Liver MLR+ vs MLR- | RPS27L | 0.005 | 0.000 | 0.65 | 0.741 | 0.538 |
| R.279 Liver MLR+ vs MLR- | ENSMMUG00000065017 | 0.012 | 0.000 | 0.46 | 0.138 | 0.028 |
| R.279 Liver MLR+ vs MLR- | FAM3C | 0.015 | 0.000 | -0.73 | 0.017 | 0.323 |
| R.279 Liver MLR+ vs MLR- | CD7 | 0.016 | 0.000 | -0.91 | 0.19 | 0.509 |
| R.279 Liver MLR+ vs MLR- | TAPBPL | 0.016 | 0.000 | -0.62 | 0.448 | 0.727 |
| R.279 Liver MLR+ vs MLR- | SCML4 | 0.024 | 0.000 | 0.48 | 0.534 | 0.255 |
| R.279 Liver MLR+ vs MLR- | CD6 | 0.040 | 0.000 | 0.51 | 0.534 | 0.286 |
| R.279 Liver MLR+ vs MLR- | UBASH3A | 0.048 | 0.000 | 0.25 | 0.362 | 0.142 |
| R.279 Liver MLR+ vs MLR- | WTAP | 0.067 | 0.000 | 0.37 | 0.724 | 0.446 |
| R.279 Liver MLR+ vs MLR- | AQR | 0.175 | 0.000 | 0.27 | 0.328 | 0.133 |
| R.279 Liver MLR+ vs MLR- | UBASH3B | 0.227 | 0.000 | -0.47 | 0.121 | 0.393 |
| R.279 Liver MLR+ vs MLR- | TUBA4A | 0.229 | 0.000 | 0.51 | 0.724 | 0.487 |
| R.279 Liver MLR+ vs MLR- | FYB1 | 0.257 | 0.000 | 0.43 | 0.793 | 0.582 |
| R.279 Liver MLR+ vs MLR- | CTLA4 | 0.294 | 0.000 | -0.69 | 0.138 | 0.412 |
| R.279 Liver MLR+ vs MLR- | EML4 | 0.323 | 0.000 | 0.47 | 0.69 | 0.471 |
| R.279 Liver MLR+ vs MLR- | CXCR4 | 0.370 | 0.000 | 0.65 | 0.793 | 0.613 |
| R.279 Liver MLR+ vs MLR- | RAP1B | 0.428 | 0.000 | 0.43 | 1 | 0.802 |
| R.279 Liver MLR+ vs MLR- | CD99 | 0.459 | 0.000 | 0.44 | 0.879 | 0.651 |
| R.279 Liver MLR+ vs MLR- | ENSMMUG00000004441 | 0.490 | 0.000 | 0.44 | 1 | 0.918 |
| R.279 Liver MLR+ vs MLR- | PCED1B | 0.512 | 0.000 | -0.45 | 0.069 | 0.322 |

| | | | | | | |
|---|---|---|---|---|---|---|
| R.279 Liver MLR+ vs MLR- | SLAMF7 | 0.540 | 0.000 | 0.29 | 0.414 | 0.197 |
| R.279 Liver MLR+ vs MLR- | TMSB10 | 0.546 | 0.000 | 0.37 | 1 | 0.997 |
| R.279 Liver MLR+ vs MLR- | KLF3 | 0.596 | 0.000 | 0.30 | 0.293 | 0.12 |
| R.279 Liver MLR+ vs MLR- | ENTPD1 | 0.642 | 0.000 | -0.42 | 0.017 | 0.252 |
| R.279 Liver MLR+ vs MLR- | CCNL1 | 0.684 | 0.000 | 0.45 | 0.862 | 0.663 |
| R.279 Liver MLR+ vs MLR- | BTG1 | 0.747 | 0.000 | 0.47 | 0.897 | 0.75 |
| R.279 Liver MLR+ vs MLR- | RGS9 | 1.000 | 0.000 | 0.39 | 0.517 | 0.294 |
| R.279 Liver MLR+ vs MLR- | IFI6 | 1.000 | 0.000 | 0.52 | 0.793 | 0.624 |
| R.279 Liver MLR+ vs MLR- | PIK3AP1 | 1.000 | 0.000 | -0.38 | 0.052 | 0.287 |
| R.279 Liver MLR+ vs MLR- | UPP1 | 1.000 | 0.000 | -0.41 | 0.086 | 0.331 |
| R.279 Liver MLR+ vs MLR- | FGL2 | 1.000 | 0.000 | 0.28 | 0.259 | 0.103 |
| R.279 Liver MLR+ vs MLR- | LITAF | 1.000 | 0.000 | -0.58 | 0.19 | 0.431 |
| R.279 Liver MLR+ vs MLR- | CD38 | 1.000 | 0.000 | -0.52 | 0.31 | 0.551 |
| R.279 Liver MLR+ vs MLR- | C15orf48 | 1.000 | 0.000 | 0.26 | 0.345 | 0.157 |
| R.279 Liver MLR+ vs MLR- | STAMBPL1 | 1.000 | 0.000 | 0.41 | 0.569 | 0.352 |
| R.279 Liver MLR+ vs MLR- | PLEK | 1.000 | 0.000 | 0.26 | 0.328 | 0.151 |
| R.279 Liver MLR+ vs MLR- | ETS2 | 1.000 | 0.000 | 0.44 | 0.345 | 0.172 |
| R.279 Liver MLR+ vs MLR- | ATF3 | 1.000 | 0.000 | 0.34 | 0.534 | 0.302 |
| R.279 Liver MLR+ vs MLR- | ANXA5 | 1.000 | 0.000 | 0.37 | 0.534 | 0.31 |
| R.279 Liver MLR+ vs MLR- | TAGAP | 1.000 | 0.000 | 0.31 | 0.69 | 0.438 |
| R.279 Liver MLR+ vs MLR- | RPL28 | 1.000 | 0.000 | -0.32 | 1 | 0.993 |
| R.279 Liver MLR+ vs MLR- | UBE2F | 1.000 | 0.000 | -0.41 | 0.121 | 0.352 |
| R.279 Liver MLR+ vs MLR- | CMA1 | 1.000 | 0.000 | 0.34 | 0.328 | 0.159 |
| R.279 Liver MLR+ vs MLR- | DUSP16 | 1.000 | 0.000 | -0.33 | 0.017 | 0.213 |
| R.279 Liver MLR+ vs MLR- | ENSMMUG00000064120 | 1.000 | 0.000 | 0.26 | 1 | 0.993 |
| R.279 Liver MLR+ vs MLR- | RAB7A | 1.000 | 0.000 | 0.29 | 0.603 | 0.377 |
| R.279 Liver MLR+ vs MLR- | MAMU-F | 1.000 | 0.000 | 0.31 | 0.379 | 0.198 |
| R.279 Liver MLR+ vs MLR- | SQOR | 1.000 | 0.000 | 0.26 | 0.31 | 0.15 |
| R.279 Liver MLR+ vs MLR- | ENSMMUG00000050862 | 1.000 | 0.000 | -0.77 | 0.069 | 0.282 |
| R.279 Liver MLR+ vs MLR- | RPS12 | 1.000 | 0.000 | -0.28 | 1 | 0.995 |
| R.279 Liver MLR+ vs MLR- | AHNAK | 1.000 | 0.000 | 0.63 | 0.655 | 0.512 |
| R.279 Liver MLR+ vs MLR- | IFI27L2 | 1.000 | 0.000 | 0.28 | 0.948 | 0.849 |
| R.279 Liver MLR+ vs MLR- | RPS5 | 1.000 | 0.000 | -0.26 | 1 | 0.993 |
| R.279 Liver MLR+ vs MLR- | RNF19B | 1.000 | 0.000 | 0.26 | 0.414 | 0.22 |
| R.279 Liver MLR+ vs MLR- | EMP3 | 1.000 | 0.000 | 0.38 | 0.914 | 0.825 |
| R.279 Liver MLR+ vs MLR- | PDCD1 | 1.000 | 0.000 | -0.50 | 0.155 | 0.364 |
| R.279 Liver MLR+ vs MLR- | LCP1 | 1.000 | 0.000 | 0.32 | 0.931 | 0.849 |
| R.279 Liver MLR+ vs MLR- | ARL6IP5 | 1.000 | 0.001 | 0.27 | 0.724 | 0.475 |
| R.279 Liver MLR+ vs MLR- | PPP1R15A | 1.000 | 0.001 | 0.43 | 0.862 | 0.696 |
| R.279 Liver MLR+ vs MLR- | RPL37A | 1.000 | 0.001 | -0.26 | 1 | 0.994 |
| R.279 Liver MLR+ vs MLR- | ENSMMUG00000039070 | 1.000 | 0.001 | -0.36 | 0.155 | 0.382 |
| R.279 Liver MLR+ vs MLR- | ANTXR2 | 1.000 | 0.001 | 0.31 | 0.19 | 0.075 |
| R.279 Liver MLR+ vs MLR- | STK39 | 1.000 | 0.001 | 0.25 | 0.5 | 0.299 |
| R.279 Liver MLR+ vs MLR- | NTRK1 | 1.000 | 0.001 | -0.32 | 0 | 0.167 |
| R.279 Liver MLR+ vs MLR- | NR3C1 | 1.000 | 0.001 | 0.29 | 0.379 | 0.208 |
| R.279 Liver MLR+ vs MLR- | ENO1 | 1.000 | 0.001 | -0.58 | 0.759 | 0.852 |
| R.279 Liver MLR+ vs MLR- | ABRACL | 1.000 | 0.001 | 0.31 | 0.845 | 0.668 |
| R.279 Liver MLR+ vs MLR- | TIGAR | 1.000 | 0.001 | -0.46 | 0.345 | 0.54 |
| R.279 Liver MLR+ vs MLR- | NEDD9 | 1.000 | 0.001 | 0.34 | 0.707 | 0.528 |
| R.279 Liver MLR+ vs MLR- | ENSMMUG00000064232 | 1.000 | 0.001 | 0.36 | 0.517 | 0.336 |
| R.279 Liver MLR+ vs MLR- | MYADM | 1.000 | 0.001 | 0.35 | 0.552 | 0.363 |
| R.279 Liver MLR+ vs MLR- | IFI27 | 1.000 | 0.001 | 0.53 | 0.69 | 0.578 |
| R.279 Liver MLR+ vs MLR- | ENSMMUG00000058581 | 1.000 | 0.001 | 0.47 | 0.845 | 0.641 |
| R.279 Liver MLR+ vs MLR- | RACK1 | 1.000 | 0.001 | -0.29 | 1 | 0.991 |
| R.279 Liver MLR+ vs MLR- | COA1 | 1.000 | 0.001 | 0.42 | 0.397 | 0.236 |
| R.279 Liver MLR+ vs MLR- | IGF2R | 1.000 | 0.001 | -0.38 | 0.121 | 0.315 |

| | | | | | | |
|---|---|---|---|---|---|---|
| R.279 Liver MLR+ vs MLR- | SH2D1A | 1.000 | 0.001 | 0.29 | 0.414 | 0.24 |
| R.279 Liver MLR+ vs MLR- | IL2RA | 1.000 | 0.001 | -0.70 | 0.138 | 0.334 |
| R.279 Liver MLR+ vs MLR- | AKR1B1 | 1.000 | 0.001 | -0.26 | 0.069 | 0.259 |
| R.279 Liver MLR+ vs MLR- | HAVCR2 | 1.000 | 0.001 | -0.28 | 0.034 | 0.207 |
| R.279 Liver MLR+ vs MLR- | HSD17B4 | 1.000 | 0.001 | 0.26 | 0.431 | 0.252 |
| R.279 Liver MLR+ vs MLR- | HDLBP | 1.000 | 0.001 | -0.34 | 0.138 | 0.334 |
| R.279 Liver MLR+ vs MLR- | LGALS3 | 1.000 | 0.001 | 0.47 | 0.828 | 0.708 |
| R.279 Liver MLR+ vs MLR- | PPP1R12A | 1.000 | 0.002 | 0.31 | 0.621 | 0.417 |
| R.279 Liver MLR+ vs MLR- | SRSF7 | 1.000 | 0.002 | 0.33 | 0.793 | 0.696 |
| R.279 Liver MLR+ vs MLR- | MYO1E | 1.000 | 0.002 | -0.34 | 0.069 | 0.243 |
| R.279 Liver MLR+ vs MLR- | H2AJ | 1.000 | 0.002 | -0.35 | 0.5 | 0.672 |
| R.279 Liver MLR+ vs MLR- | TNFRSF1B | 1.000 | 0.002 | -0.45 | 0.638 | 0.755 |
| R.279 Liver MLR+ vs MLR- | ZFP36 | 1.000 | 0.002 | 0.38 | 0.914 | 0.772 |
| R.279 Liver MLR+ vs MLR- | CRIP1 | 1.000 | 0.002 | 0.42 | 0.983 | 0.952 |
| R.279 Liver MLR+ vs MLR- | SLBP | 1.000 | 0.002 | 0.30 | 0.586 | 0.408 |
| R.279 Liver MLR+ vs MLR- | CDKN1A | 1.000 | 0.002 | 0.39 | 0.552 | 0.364 |
| R.279 Liver MLR+ vs MLR- | PRDX6 | 1.000 | 0.002 | 0.34 | 0.879 | 0.763 |
| R.279 Liver MLR+ vs MLR- | BATF | 1.000 | 0.002 | 0.25 | 0.517 | 0.319 |
| R.279 Liver MLR+ vs MLR- | MRPL4 | 1.000 | 0.002 | -0.26 | 0.069 | 0.241 |
| R.279 Liver MLR+ vs MLR- | SSR1 | 1.000 | 0.002 | -0.30 | 0.19 | 0.386 |
| R.279 Liver MLR+ vs MLR- | STAT3 | 1.000 | 0.002 | -0.33 | 0.19 | 0.382 |
| R.279 Liver MLR+ vs MLR- | SERPINE2 | 1.000 | 0.002 | -0.45 | 0 | 0.14 |
| R.279 Liver MLR+ vs MLR- | CLTC | 1.000 | 0.002 | 0.26 | 0.466 | 0.3 |
| R.279 Liver MLR+ vs MLR- | USP3 | 1.000 | 0.002 | 0.31 | 0.345 | 0.197 |
| R.279 Liver MLR+ vs MLR- | CCL5 | 1.000 | 0.002 | 0.40 | 0.862 | 0.706 |
| R.279 Liver MLR+ vs MLR- | ENSMMUG00000003532 | 1.000 | 0.002 | -0.47 | 0.448 | 0.643 |
| R.279 Liver MLR+ vs MLR- | ARRB2 | 1.000 | 0.002 | 0.34 | 0.69 | 0.533 |
| R.279 Liver MLR+ vs MLR- | ALOX5AP | 1.000 | 0.002 | 0.27 | 0.241 | 0.114 |
| R.279 Liver MLR+ vs MLR- | PLAC8 | 1.000 | 0.003 | -0.67 | 0.259 | 0.437 |
| R.279 Liver MLR+ vs MLR- | SERTAD1 | 1.000 | 0.003 | 0.36 | 0.724 | 0.568 |
| R.279 Liver MLR+ vs MLR- | LMO4 | 1.000 | 0.003 | -0.39 | 0.155 | 0.329 |
| R.279 Liver MLR+ vs MLR- | IL4R | 1.000 | 0.003 | -0.27 | 0.207 | 0.418 |
| R.279 Liver MLR+ vs MLR- | KIR2DL4 | 1.000 | 0.003 | -0.42 | 0.017 | 0.158 |
| R.279 Liver MLR+ vs MLR- | PTPN7 | 1.000 | 0.003 | -0.27 | 0.121 | 0.303 |
| R.279 Liver MLR+ vs MLR- | S100A4 | 1.000 | 0.003 | 0.40 | 0.983 | 0.93 |
| R.279 Liver MLR+ vs MLR- | ENSMMUG00000053735 | 1.000 | 0.003 | -0.32 | 0.293 | 0.476 |
| R.279 Liver MLR+ vs MLR- | CA6 | 1.000 | 0.003 | -0.35 | 0.086 | 0.251 |
| R.279 Liver MLR+ vs MLR- | IDH2 | 1.000 | 0.003 | 0.43 | 0.69 | 0.628 |
| R.279 Liver MLR+ vs MLR- | TIMM8B | 1.000 | 0.003 | -0.25 | 0.069 | 0.229 |
| R.279 Liver MLR+ vs MLR- | TNFSF11 | 1.000 | 0.004 | -0.29 | 0 | 0.129 |
| R.279 Liver MLR+ vs MLR- | CD48 | 1.000 | 0.004 | 0.35 | 0.983 | 0.891 |
| R.279 Liver MLR+ vs MLR- | EIF5B | 1.000 | 0.004 | -0.27 | 0.103 | 0.269 |
| R.279 Liver MLR+ vs MLR- | RNF181 | 1.000 | 0.004 | -0.28 | 0.241 | 0.419 |
| R.279 Liver MLR+ vs MLR- | SQSTM1 | 1.000 | 0.004 | 0.34 | 0.862 | 0.656 |
| R.279 Liver MLR+ vs MLR- | NKG7 | 1.000 | 0.004 | 0.27 | 0.966 | 0.894 |
| R.279 Liver MLR+ vs MLR- | JUND | 1.000 | 0.004 | 0.26 | 0.741 | 0.561 |
| R.279 Liver MLR+ vs MLR- | OSBPL2 | 1.000 | 0.004 | 0.28 | 0.672 | 0.515 |
| R.279 Liver MLR+ vs MLR- | ITM2A | 1.000 | 0.004 | 0.26 | 0.448 | 0.293 |
| R.279 Liver MLR+ vs MLR- | LYST | 1.000 | 0.005 | -0.30 | 0.172 | 0.344 |
| R.279 Liver MLR+ vs MLR- | TBC1D25 | 1.000 | 0.005 | 0.29 | 0.5 | 0.338 |
| R.279 Liver MLR+ vs MLR- | MGST3 | 1.000 | 0.005 | 0.27 | 0.414 | 0.256 |
| R.279 Liver MLR+ vs MLR- | PLAAT4 | 1.000 | 0.005 | 0.50 | 0.776 | 0.671 |
| R.279 Liver MLR+ vs MLR- | PMAIP1 | 1.000 | 0.005 | 0.35 | 0.621 | 0.458 |
| R.279 Liver MLR+ vs MLR- | ISG20 | 1.000 | 0.005 | 0.33 | 0.862 | 0.67 |
| R.279 Liver MLR+ vs MLR- | PSMB9 | 1.000 | 0.005 | 0.33 | 0.897 | 0.842 |
| R.279 Liver MLR+ vs MLR- | ENSMMUG00000014786 | 1.000 | 0.005 | -0.27 | 0.983 | 0.989 |

| | | | | | | |
|---|---|---|---|---|---|---|
| R.279 Liver MLR+ vs MLR- | SUB1 | 1.000 | 0.005 | 0.38 | 0.776 | 0.675 |
| R.279 Liver MLR+ vs MLR- | RGS2 | 1.000 | 0.005 | -0.26 | 0.155 | 0.345 |
| R.279 Liver MLR+ vs MLR- | SRSF5 | 1.000 | 0.005 | 0.27 | 0.879 | 0.701 |
| R.279 Liver MLR+ vs MLR- | GNA13 | 1.000 | 0.005 | 0.32 | 0.534 | 0.401 |
| R.279 Liver MLR+ vs MLR- | ENSMMUG00000053539 | 1.000 | 0.006 | 0.26 | 0.379 | 0.222 |
| R.279 Liver MLR+ vs MLR- | ENSMMUG00000045208 | 1.000 | 0.006 | 0.25 | 0.862 | 0.78 |
| R.279 Liver MLR+ vs MLR- | PGK1 | 1.000 | 0.006 | -0.36 | 0.569 | 0.682 |
| R.279 Liver MLR+ vs MLR- | CLTB | 1.000 | 0.006 | -0.26 | 0.138 | 0.297 |
| R.279 Liver MLR+ vs MLR- | NUDC | 1.000 | 0.007 | -0.34 | 0.397 | 0.535 |
| R.279 Liver MLR+ vs MLR- | PARK7 | 1.000 | 0.007 | -0.28 | 0.345 | 0.508 |
| R.279 Liver MLR+ vs MLR- | HDAC7 | 1.000 | 0.007 | -0.26 | 0.121 | 0.275 |
| R.279 Liver MLR+ vs MLR- | ARF1 | 1.000 | 0.007 | 0.29 | 0.862 | 0.81 |
| R.279 Liver MLR+ vs MLR- | PELI1 | 1.000 | 0.007 | -0.35 | 0.293 | 0.456 |
| R.279 Liver MLR+ vs MLR- | DUSP4 | 1.000 | 0.007 | -0.26 | 0.138 | 0.307 |
| R.279 Liver MLR+ vs MLR- | ENSMMUG00000013779 | 1.000 | 0.007 | -0.52 | 0 | 0.113 |
| R.279 Liver MLR+ vs MLR- | GLIPR2 | 1.000 | 0.007 | 0.27 | 0.621 | 0.47 |
| R.279 Liver MLR+ vs MLR- | RPL4 | 1.000 | 0.007 | -0.25 | 0.966 | 0.983 |
| R.279 Liver MLR+ vs MLR- | TSC22D3 | 1.000 | 0.007 | 0.35 | 0.828 | 0.738 |
| R.279 Liver MLR+ vs MLR- | FLNA | 1.000 | 0.007 | 0.48 | 0.707 | 0.624 |
| R.279 Liver MLR+ vs MLR- | FNBP1 | 1.000 | 0.007 | 0.29 | 0.759 | 0.601 |
| R.279 Liver MLR+ vs MLR- | PDE4B | 1.000 | 0.007 | 0.28 | 0.517 | 0.349 |
| R.279 Liver MLR+ vs MLR- | RASSF5 | 1.000 | 0.007 | 0.26 | 0.534 | 0.386 |
| R.279 Liver MLR+ vs MLR- | ANXA6 | 1.000 | 0.007 | -0.33 | 0.448 | 0.6 |
| R.279 Liver MLR+ vs MLR- | GSTP1 | 1.000 | 0.007 | -0.27 | 0.931 | 0.925 |
| R.279 Liver MLR+ vs MLR- | EMC3 | 1.000 | 0.007 | 0.26 | 0.397 | 0.258 |
| R.279 Liver MLR+ vs MLR- | PSME1 | 1.000 | 0.007 | 0.26 | 0.931 | 0.919 |
| R.279 Liver MLR+ vs MLR- | AHI1 | 1.000 | 0.008 | -0.34 | 0.241 | 0.392 |
| R.279 Liver MLR+ vs MLR- | TTC39C | 1.000 | 0.008 | 0.25 | 0.397 | 0.249 |
| R.279 Liver MLR+ vs MLR- | ITGB7 | 1.000 | 0.008 | 0.26 | 0.483 | 0.323 |
| R.279 Liver MLR+ vs MLR- | ATP5F1D | 1.000 | 0.008 | -0.26 | 0.828 | 0.844 |
| R.279 Liver MLR+ vs MLR- | PEBP1 | 1.000 | 0.008 | -0.28 | 0.379 | 0.566 |
| R.279 Liver MLR+ vs MLR- | ZC3H10 | 1.000 | 0.008 | -0.30 | 0.845 | 0.894 |
| R.279 Liver MLR+ vs MLR- | CCDC85B | 1.000 | 0.009 | -0.28 | 0.224 | 0.381 |
| R.279 Liver MLR+ vs MLR- | APBB1IP | 1.000 | 0.009 | 0.26 | 0.672 | 0.481 |
| R.279 Liver MLR+ vs MLR- | MIF | 1.000 | 0.009 | -0.38 | 0.759 | 0.803 |
| R.279 Liver MLR+ vs MLR- | ENSMMUG00000061128 | 1.000 | 0.009 | 0.26 | 0.776 | 0.582 |
| R.279 Liver MLR+ vs MLR- | ENSMMUG00000006499 | 1.000 | 0.009 | -0.26 | 0.19 | 0.347 |
| R.279 Liver MLR+ vs MLR- | CD37 | 1.000 | 0.010 | 0.28 | 0.707 | 0.559 |
| R.279 Liver MLR+ vs MLR- | DNAJB6 | 1.000 | 0.010 | 0.26 | 0.621 | 0.48 |
| R.279 Liver MLR+ vs MLR- | ENSMMUG00000043332 | 1.000 | 0.010 | 0.25 | 0.81 | 0.631 |
| R.279 Liver MLR+ vs MLR- | ICOS | 1.000 | 0.010 | -0.32 | 0.207 | 0.369 |
| R.279 Liver MLR+ vs MLR- | IRF8 | 1.000 | 0.010 | -0.33 | 0.259 | 0.422 |
| R.279 Liver MLR+ vs MLR- | TANK | 1.000 | 0.011 | 0.27 | 0.397 | 0.257 |
| R.279 Liver MLR+ vs MLR- | XCL1 | 1.000 | 0.011 | -0.75 | 0.034 | 0.151 |
| R.279 Liver MLR+ vs MLR- | DNAJC3 | 1.000 | 0.012 | -0.26 | 0.276 | 0.433 |
| R.279 Liver MLR+ vs MLR- | SLC25A4 | 1.000 | 0.012 | 0.27 | 0.466 | 0.332 |
| R.279 Liver MLR+ vs MLR- | NDUFB9 | 1.000 | 0.012 | -0.30 | 0.448 | 0.565 |
| R.279 Liver MLR+ vs MLR- | CD8A | 1.000 | 0.013 | -0.36 | 0.672 | 0.766 |
| R.279 Liver MLR+ vs MLR- | TUBB2B | 1.000 | 0.013 | 0.38 | 0.638 | 0.545 |
| R.279 Liver MLR+ vs MLR- | PARP6 | 1.000 | 0.014 | -0.36 | 0.466 | 0.57 |
| R.279 Liver MLR+ vs MLR- | PTGER4 | 1.000 | 0.014 | 0.33 | 0.414 | 0.279 |
| R.279 Liver MLR+ vs MLR- | GBP3 | 1.000 | 0.014 | 0.34 | 0.69 | 0.552 |
| R.279 Liver MLR+ vs MLR- | GATA3 | 1.000 | 0.014 | -0.29 | 0.172 | 0.318 |
| R.279 Liver MLR+ vs MLR- | ENSMMUG00000003854 | 1.000 | 0.014 | 0.29 | 0.655 | 0.517 |
| R.279 Liver MLR+ vs MLR- | CLEC2D | 1.000 | 0.015 | 0.26 | 0.914 | 0.737 |
| R.279 Liver MLR+ vs MLR- | PDIA6 | 1.000 | 0.015 | -0.34 | 0.345 | 0.473 |

| | | | | | | |
|---|---|---|---|---|---|---|
| R.279 Liver MLR+ vs MLR- | MRPL10 | 1.000 | 0.016 | 0.26 | 0.448 | 0.325 |
| R.279 Liver MLR+ vs MLR- | ZYX | 1.000 | 0.016 | 0.26 | 0.638 | 0.52 |
| R.279 Liver MLR+ vs MLR- | RRAD | 1.000 | 0.016 | 0.30 | 0.448 | 0.313 |
| R.279 Liver MLR+ vs MLR- | OSR2 | 1.000 | 0.016 | 0.26 | 0.224 | 0.126 |
| R.279 Liver MLR+ vs MLR- | HK3 | 1.000 | 0.017 | -0.29 | 0.052 | 0.169 |
| R.279 Liver MLR+ vs MLR- | COX3 | 1.000 | 0.019 | -0.50 | 0.776 | 0.865 |
| R.279 Liver MLR+ vs MLR- | NPC2 | 1.000 | 0.019 | -0.26 | 0.069 | 0.187 |
| R.279 Liver MLR+ vs MLR- | CD69 | 1.000 | 0.020 | 0.28 | 0.828 | 0.72 |
| R.279 Liver MLR+ vs MLR- | PLK3 | 1.000 | 0.020 | 0.32 | 0.276 | 0.168 |
| R.279 Liver MLR+ vs MLR- | LCP2 | 1.000 | 0.020 | 0.32 | 0.586 | 0.465 |
| R.279 Liver MLR+ vs MLR- | CIB1 | 1.000 | 0.021 | 0.30 | 0.655 | 0.556 |
| R.279 Liver MLR+ vs MLR- | COX1 | 1.000 | 0.021 | -0.46 | 0.776 | 0.841 |
| R.279 Liver MLR+ vs MLR- | C1QBP | 1.000 | 0.021 | -0.27 | 0.259 | 0.395 |
| R.279 Liver MLR+ vs MLR- | IRF1 | 1.000 | 0.022 | 0.27 | 0.793 | 0.681 |
| R.279 Liver MLR+ vs MLR- | ITGAX | 1.000 | 0.022 | -0.30 | 0.086 | 0.205 |
| R.279 Liver MLR+ vs MLR- | ENSMMUG00000061125 | 1.000 | 0.022 | -0.26 | 0.121 | 0.247 |
| R.279 Liver MLR+ vs MLR- | CANX | 1.000 | 0.023 | -0.34 | 0.362 | 0.463 |
| R.279 Liver MLR+ vs MLR- | PDIA4 | 1.000 | 0.023 | -0.34 | 0.224 | 0.35 |
| R.279 Liver MLR+ vs MLR- | IRF4 | 1.000 | 0.024 | -0.38 | 0.293 | 0.413 |
| R.279 Liver MLR+ vs MLR- | HTATSF1 | 1.000 | 0.024 | 0.26 | 0.362 | 0.249 |
| R.279 Liver MLR+ vs MLR- | PPIB | 1.000 | 0.025 | -0.28 | 0.897 | 0.851 |
| R.279 Liver MLR+ vs MLR- | FOS | 1.000 | 0.025 | 0.28 | 0.828 | 0.653 |
| R.279 Liver MLR+ vs MLR- | RGS1 | 1.000 | 0.025 | 0.33 | 0.724 | 0.61 |
| R.279 Liver MLR+ vs MLR- | CD44 | 1.000 | 0.025 | 0.25 | 0.793 | 0.735 |
| R.279 Liver MLR+ vs MLR- | PFKFB3 | 1.000 | 0.026 | 0.26 | 0.31 | 0.201 |
| R.279 Liver MLR+ vs MLR- | PSMB3 | 1.000 | 0.027 | -0.26 | 0.483 | 0.585 |
| R.279 Liver MLR+ vs MLR- | GZMK | 1.000 | 0.030 | -0.26 | 0.345 | 0.52 |
| R.279 Liver MLR+ vs MLR- | ANXA2 | 1.000 | 0.030 | 0.30 | 0.776 | 0.735 |
| R.279 Liver MLR+ vs MLR- | SH3BGRL | 1.000 | 0.032 | 0.25 | 0.5 | 0.363 |
| R.279 Liver MLR+ vs MLR- | ENSMMUG00000057791 | 1.000 | 0.032 | -0.34 | 0.017 | 0.101 |
| R.279 Liver MLR+ vs MLR- | SERPINB1 | 1.000 | 0.033 | -0.52 | 0.621 | 0.67 |
| R.279 Liver MLR+ vs MLR- | ENSMMUG00000015272 | 1.000 | 0.034 | -0.29 | 0.466 | 0.56 |
| R.279 Liver MLR+ vs MLR- | THY1 | 1.000 | 0.037 | -0.25 | 0.224 | 0.354 |
| R.279 Liver MLR+ vs MLR- | TAF15 | 1.000 | 0.040 | 0.26 | 0.569 | 0.464 |
| R.279 Liver MLR+ vs MLR- | BST2 | 1.000 | 0.040 | -0.28 | 0.69 | 0.741 |
| R.279 Liver MLR+ vs MLR- | H1FX | 1.000 | 0.042 | -0.28 | 0.276 | 0.397 |
| R.279 Liver MLR+ vs MLR- | TBCC | 1.000 | 0.044 | 0.35 | 0.5 | 0.414 |
| R.279 Liver MLR+ vs MLR- | SH3KBP1 | 1.000 | 0.046 | 0.27 | 0.638 | 0.597 |
| R.279 Liver MLR+ vs MLR- | SAMSN1 | 1.000 | 0.046 | -0.29 | 0.31 | 0.404 |
| R.279 Liver MLR+ vs MLR- | DBI | 1.000 | 0.047 | 0.28 | 0.707 | 0.656 |
| R.279 Liver MLR+ vs MLR- | ENSMMUG00000013256 | 1.000 | 0.048 | 0.27 | 0.655 | 0.549 |
| R.279 Liver MLR+ vs MLR- | TXNIP | 1.000 | 0.049 | 0.36 | 0.345 | 0.246 |
| R.279 Liver MLR+ vs MLR- | GZMB | 1.000 | 0.054 | -0.62 | 0.897 | 0.871 |
| R.279 Liver MLR+ vs MLR- | ENSMMUG00000056515 | 1.000 | 0.058 | -0.79 | 0.086 | 0.176 |
| R.279 Liver MLR+ vs MLR- | SPN | 1.000 | 0.065 | 0.35 | 0.448 | 0.383 |
| R.279 Liver MLR+ vs MLR- | KLF2 | 1.000 | 0.065 | 0.29 | 0.724 | 0.601 |
| R.279 Liver MLR+ vs MLR- | SDF2L1 | 1.000 | 0.066 | -0.26 | 0.31 | 0.41 |
| R.279 Liver MLR+ vs MLR- | LSM14A | 1.000 | 0.067 | 0.25 | 0.5 | 0.399 |
| R.279 Liver MLR+ vs MLR- | JUNB | 1.000 | 0.068 | 0.31 | 0.466 | 0.369 |
| R.279 Liver MLR+ vs MLR- | TRAC | 1.000 | 0.087 | -0.27 | 0.655 | 0.663 |
| R.279 Liver MLR+ vs MLR- | GPR183 | 1.000 | 0.091 | -0.37 | 0.224 | 0.323 |
| R.279 Liver MLR+ vs MLR- | ENSMMUG00000063583 | 1.000 | 0.094 | -0.61 | 1 | 0.886 |
| R.279 Liver MLR+ vs MLR- | ATP6 | 1.000 | 0.095 | -0.39 | 0.862 | 0.878 |
| R.279 Liver MLR+ vs MLR- | CGA | 1.000 | 0.095 | -0.61 | 0.086 | 0.163 |
| R.279 Liver MLR+ vs MLR- | FABP5 | 1.000 | 0.099 | 0.30 | 0.345 | 0.456 |
| R.279 Liver MLR+ vs MLR- | SELL | 1.000 | 0.114 | -0.27 | 0.086 | 0.158 |

| | | | | | | |
|---|---|---|---|---|---|---|
| R.279 Liver MLR+ vs MLR- | MT2A | 1.000 | 0.116 | 0.26 | 0.483 | 0.6 |
| R.279 Liver MLR+ vs MLR- | TPI1 | 1.000 | 0.138 | -0.25 | 0.81 | 0.825 |
| R.279 Liver MLR+ vs MLR- | CTSW | 1.000 | 0.139 | -0.36 | 0.293 | 0.358 |
| R.279 Liver MLR+ vs MLR- | HSP90B1 | 1.000 | 0.153 | -0.34 | 0.81 | 0.771 |
| R.279 Liver MLR+ vs MLR- | GADD45B | 1.000 | 0.155 | 0.47 | 0.621 | 0.564 |
| R.279 Liver MLR+ vs MLR- | TIGIT | 1.000 | 0.181 | -0.28 | 0.5 | 0.539 |
| R.279 Liver MLR+ vs MLR- | RGCC | 1.000 | 0.187 | 0.38 | 0.793 | 0.759 |
| R.279 Liver MLR+ vs MLR- | CCL3 | 1.000 | 0.191 | -0.47 | 0.483 | 0.547 |
| R.279 Liver MLR+ vs MLR- | JUN | 1.000 | 0.194 | 0.35 | 0.828 | 0.79 |
| R.279 Liver MLR+ vs MLR- | CTSC | 1.000 | 0.212 | -0.30 | 0.5 | 0.547 |
| R.279 Liver MLR+ vs MLR- | PCLAF | 1.000 | 0.217 | 0.25 | 0.31 | 0.252 |
| R.279 Liver MLR+ vs MLR- | ENSMMUG00000062077 | 1.000 | 0.415 | -1.21 | 1 | 0.999 |
| R.279 Liver MLR+ vs MLR- | ENSMMUG00000051385 | 1.000 | 0.456 | 0.36 | 0.172 | 0.145 |
| R.279 Liver MLR+ vs MLR- | COTL1 | 1.000 | 0.463 | 0.27 | 0.793 | 0.857 |
| R.279 Liver MLR+ vs MLR- | TYROBP | 1.000 | 0.469 | -0.36 | 0.103 | 0.128 |
| R.279 Liver MLR+ vs MLR- | ENSMMUG00000062894 | 1.000 | 0.496 | -0.33 | 0.638 | 0.6 |
| R.279 Liver MLR+ vs MLR- | ZFP36L1 | 1.000 | 0.557 | 0.28 | 0.569 | 0.577 |
| R.279 Liver MLR+ vs MLR- | HIST2H3A | 1.000 | 0.713 | -0.33 | 0.138 | 0.155 |
| R.279 Liver MLR+ vs MLR- | STMN1 | 1.000 | 0.751 | 0.43 | 0.483 | 0.519 |
| R.279 Liver MLR+ vs MLR- | RRM2 | 1.000 | 0.802 | 0.27 | 0.328 | 0.333 |
| R.279 Liver MLR+ vs MLR- | COX2 | 1.000 | 0.838 | -0.28 | 0.741 | 0.754 |
| R.279 Spleen MLR+ vs MLR- | CDKN2C | 0.000 | 0.000 | 0.30 | 0.216 | 0.028 |
| R.279 Spleen MLR+ vs MLR- | GOLIM4 | 0.000 | 0.000 | 0.42 | 0.432 | 0.103 |
| R.279 Spleen MLR+ vs MLR- | CD96 | 0.000 | 0.000 | 0.50 | 0.595 | 0.192 |
| R.279 Spleen MLR+ vs MLR- | CD7 | 0.000 | 0.000 | -1.47 | 0.135 | 0.653 |
| R.279 Spleen MLR+ vs MLR- | PPP2R2B | 0.000 | 0.000 | 0.31 | 0.351 | 0.075 |
| R.279 Spleen MLR+ vs MLR- | MAMU-DRB1 | 0.000 | 0.000 | 0.72 | 0.649 | 0.227 |
| R.279 Spleen MLR+ vs MLR- | CLIC5 | 0.000 | 0.000 | 0.48 | 0.405 | 0.103 |
| R.279 Spleen MLR+ vs MLR- | ENSMMUG00000065017 | 0.000 | 0.000 | 0.76 | 0.189 | 0.027 |
| R.279 Spleen MLR+ vs MLR- | S100A10 | 0.000 | 0.000 | 0.98 | 0.946 | 0.821 |
| R.279 Spleen MLR+ vs MLR- | S100A4 | 0.000 | 0.000 | 0.99 | 0.973 | 0.719 |
| R.279 Spleen MLR+ vs MLR- | ENSMMUG00000019371 | 0.001 | 0.000 | 0.31 | 0.297 | 0.068 |
| R.279 Spleen MLR+ vs MLR- | RPS27A.1 | 0.002 | 0.000 | -0.58 | 1 | 0.996 |
| R.279 Spleen MLR+ vs MLR- | NKG7 | 0.002 | 0.000 | 0.82 | 1 | 0.688 |
| R.279 Spleen MLR+ vs MLR- | SH3BP5 | 0.002 | 0.000 | 0.54 | 0.486 | 0.167 |
| R.279 Spleen MLR+ vs MLR- | TSPAN2 | 0.002 | 0.000 | 0.33 | 0.324 | 0.082 |
| R.279 Spleen MLR+ vs MLR- | ANXA1 | 0.002 | 0.000 | 0.99 | 0.892 | 0.573 |
| R.279 Spleen MLR+ vs MLR- | ENSMMUG00000052609 | 0.003 | 0.000 | -0.61 | 1 | 0.995 |
| R.279 Spleen MLR+ vs MLR- | EFHD2 | 0.004 | 0.000 | 0.83 | 0.811 | 0.49 |
| R.279 Spleen MLR+ vs MLR- | ENSMMUG00000060382 | 0.005 | 0.000 | 0.48 | 0.459 | 0.157 |
| R.279 Spleen MLR+ vs MLR- | RPS3A | 0.007 | 0.000 | -0.53 | 1 | 0.996 |
| R.279 Spleen MLR+ vs MLR- | RPS13 | 0.012 | 0.000 | -0.58 | 1 | 0.994 |
| R.279 Spleen MLR+ vs MLR- | RGS9 | 0.013 | 0.000 | 0.53 | 0.676 | 0.305 |
| R.279 Spleen MLR+ vs MLR- | SPOCK2 | 0.016 | 0.000 | -0.97 | 0 | 0.413 |
| R.279 Spleen MLR+ vs MLR- | HABP4 | 0.016 | 0.000 | 0.29 | 0.243 | 0.056 |
| R.279 Spleen MLR+ vs MLR- | IL7R | 0.032 | 0.000 | 0.90 | 0.811 | 0.439 |
| R.279 Spleen MLR+ vs MLR- | CST7 | 0.035 | 0.000 | 0.60 | 0.973 | 0.616 |
| R.279 Spleen MLR+ vs MLR- | CCL5 | 0.036 | 0.000 | 0.75 | 0.946 | 0.529 |
| R.279 Spleen MLR+ vs MLR- | GZMB | 0.039 | 0.000 | 0.75 | 0.838 | 0.489 |
| R.279 Spleen MLR+ vs MLR- | ITGB2 | 0.045 | 0.000 | 0.58 | 0.946 | 0.688 |
| R.279 Spleen MLR+ vs MLR- | SLCO4C1 | 0.048 | 0.000 | 0.26 | 0.189 | 0.039 |
| R.279 Spleen MLR+ vs MLR- | B2M | 0.049 | 0.000 | 0.33 | 1 | 0.997 |
| R.279 Spleen MLR+ vs MLR- | CENPU | 0.050 | 0.000 | 0.25 | 0.189 | 0.038 |
| R.279 Spleen MLR+ vs MLR- | CDT1 | 0.050 | 0.000 | 0.32 | 0.297 | 0.083 |
| R.279 Spleen MLR+ vs MLR- | ENSMMUG00000063583 | 0.065 | 0.000 | 0.67 | 0.946 | 0.651 |
| R.279 Spleen MLR+ vs MLR- | ENSMMUG00000053403 | 0.065 | 0.000 | 0.28 | 0.162 | 0.03 |

| | | | | | | |
|---|---|---|---|---|---|---|
| R.279 Spleen MLR+ vs MLR- | NCAPG | 0.069 | 0.000 | 0.38 | 0.243 | 0.059 |
| R.279 Spleen MLR+ vs MLR- | RPL28 | 0.078 | 0.000 | -0.53 | 1 | 0.996 |
| R.279 Spleen MLR+ vs MLR- | ENSMMUG00000004441 | 0.084 | 0.000 | 0.63 | 1 | 0.882 |
| R.279 Spleen MLR+ vs MLR- | RPL24 | 0.093 | 0.000 | -0.48 | 1 | 0.991 |
| R.279 Spleen MLR+ vs MLR- | CX3CR1 | 0.095 | 0.000 | 0.51 | 0.243 | 0.062 |
| R.279 Spleen MLR+ vs MLR- | GNG5 | 0.097 | 0.000 | 0.55 | 0.865 | 0.597 |
| R.279 Spleen MLR+ vs MLR- | ACD | 0.099 | 0.000 | 0.27 | 0.27 | 0.074 |
| R.279 Spleen MLR+ vs MLR- | GPR183 | 0.137 | 0.000 | -1.10 | 0.081 | 0.467 |
| R.279 Spleen MLR+ vs MLR- | DHRS7 | 0.144 | 0.000 | 0.48 | 0.541 | 0.239 |
| R.279 Spleen MLR+ vs MLR- | SMC6 | 0.154 | 0.000 | 0.25 | 0.324 | 0.1 |
| R.279 Spleen MLR+ vs MLR- | RPS4X | 0.157 | 0.000 | -0.42 | 1 | 0.996 |
| R.279 Spleen MLR+ vs MLR- | AUH | 0.158 | 0.000 | 0.26 | 0.324 | 0.1 |
| R.279 Spleen MLR+ vs MLR- | CKS1B | 0.168 | 0.000 | 0.43 | 0.324 | 0.104 |
| R.279 Spleen MLR+ vs MLR- | RPS24 | 0.231 | 0.000 | -0.44 | 1 | 0.991 |
| R.279 Spleen MLR+ vs MLR- | KLRD1 | 0.255 | 0.000 | 0.58 | 0.811 | 0.449 |
| R.279 Spleen MLR+ vs MLR- | RPL10A | 0.262 | 0.000 | -0.56 | 1 | 0.992 |
| R.279 Spleen MLR+ vs MLR- | RPLP1 | 0.291 | 0.000 | -0.46 | 1 | 0.994 |
| R.279 Spleen MLR+ vs MLR- | OSR2 | 0.332 | 0.000 | 0.40 | 0.27 | 0.08 |
| R.279 Spleen MLR+ vs MLR- | ENSMMUG00000063637 | 0.492 | 0.000 | -0.46 | 1 | 0.994 |
| R.279 Spleen MLR+ vs MLR- | RPL13 | 0.494 | 0.000 | -0.39 | 1 | 0.997 |
| R.279 Spleen MLR+ vs MLR- | EMP3 | 0.496 | 0.000 | 0.62 | 0.919 | 0.758 |
| R.279 Spleen MLR+ vs MLR- | RPS16 | 0.517 | 0.000 | -0.43 | 1 | 0.993 |
| R.279 Spleen MLR+ vs MLR- | CMA1 | 0.544 | 0.000 | 0.40 | 0.324 | 0.111 |
| R.279 Spleen MLR+ vs MLR- | CDCA8 | 0.578 | 0.000 | 0.28 | 0.243 | 0.069 |
| R.279 Spleen MLR+ vs MLR- | RPS8 | 0.590 | 0.000 | -0.45 | 1 | 0.997 |
| R.279 Spleen MLR+ vs MLR- | EEF1A1 | 0.659 | 0.000 | -0.41 | 1 | 0.996 |
| R.279 Spleen MLR+ vs MLR- | NUSAP1 | 0.854 | 0.000 | 0.28 | 0.243 | 0.069 |
| R.279 Spleen MLR+ vs MLR- | RPS12 | 0.993 | 0.000 | -0.38 | 1 | 0.998 |
| R.279 Spleen MLR+ vs MLR- | RPS14 | 1.000 | 0.000 | -0.40 | 1 | 0.992 |
| R.279 Spleen MLR+ vs MLR- | CDK1 | 1.000 | 0.000 | 0.50 | 0.243 | 0.077 |
| R.279 Spleen MLR+ vs MLR- | ZBTB38 | 1.000 | 0.000 | 0.27 | 0.459 | 0.192 |
| R.279 Spleen MLR+ vs MLR- | PRKAA1 | 1.000 | 0.000 | 0.29 | 0.324 | 0.116 |
| R.279 Spleen MLR+ vs MLR- | IDH2 | 1.000 | 0.000 | 0.45 | 0.811 | 0.498 |
| R.279 Spleen MLR+ vs MLR- | MID1IP1 | 1.000 | 0.000 | 0.29 | 0.27 | 0.088 |
| R.279 Spleen MLR+ vs MLR- | SMC2 | 1.000 | 0.000 | 0.38 | 0.297 | 0.104 |
| R.279 Spleen MLR+ vs MLR- | RPS17 | 1.000 | 0.000 | -0.48 | 0.838 | 0.94 |
| R.279 Spleen MLR+ vs MLR- | RACK1 | 1.000 | 0.000 | -0.42 | 1 | 0.992 |
| R.279 Spleen MLR+ vs MLR- | SPC25 | 1.000 | 0.000 | 0.34 | 0.189 | 0.05 |
| R.279 Spleen MLR+ vs MLR- | RHOH | 1.000 | 0.000 | -0.57 | 0.135 | 0.452 |
| R.279 Spleen MLR+ vs MLR- | ARRB2 | 1.000 | 0.000 | 0.44 | 0.676 | 0.418 |
| R.279 Spleen MLR+ vs MLR- | RPS3 | 1.000 | 0.000 | -0.42 | 1 | 0.992 |
| R.279 Spleen MLR+ vs MLR- | PLAC8 | 1.000 | 0.000 | -0.88 | 0.324 | 0.627 |
| R.279 Spleen MLR+ vs MLR- | ENSMMUG00000061750 | 1.000 | 0.000 | 0.31 | 0.243 | 0.076 |
| R.279 Spleen MLR+ vs MLR- | RPL4 | 1.000 | 0.000 | -0.46 | 0.973 | 0.981 |
| R.279 Spleen MLR+ vs MLR- | SLC11A1 | 1.000 | 0.000 | 0.33 | 0.378 | 0.157 |
| R.279 Spleen MLR+ vs MLR- | RPL37A | 1.000 | 0.000 | -0.37 | 1 | 0.997 |
| R.279 Spleen MLR+ vs MLR- | RPL22 | 1.000 | 0.000 | -0.40 | 1 | 0.989 |
| R.279 Spleen MLR+ vs MLR- | HNRNPA2B1 | 1.000 | 0.000 | 0.50 | 0.919 | 0.896 |
| R.279 Spleen MLR+ vs MLR- | CLIC1 | 1.000 | 0.000 | 0.58 | 0.892 | 0.702 |
| R.279 Spleen MLR+ vs MLR- | ENSMMUG00000014786 | 1.000 | 0.000 | -0.38 | 1 | 0.992 |
| R.279 Spleen MLR+ vs MLR- | RPS25 | 1.000 | 0.000 | -0.40 | 1 | 0.99 |
| R.279 Spleen MLR+ vs MLR- | SMC4 | 1.000 | 0.000 | 0.48 | 0.405 | 0.181 |
| R.279 Spleen MLR+ vs MLR- | RPA2 | 1.000 | 0.000 | 0.31 | 0.351 | 0.142 |
| R.279 Spleen MLR+ vs MLR- | CPD | 1.000 | 0.000 | 0.47 | 0.622 | 0.341 |
| R.279 Spleen MLR+ vs MLR- | GCN1 | 1.000 | 0.000 | -0.43 | 1 | 0.995 |
| R.279 Spleen MLR+ vs MLR- | RPL6 | 1.000 | 0.000 | -0.37 | 1 | 0.997 |

| | | | | | | |
|---|---|---|---|---|---|---|
| R.279 Spleen MLR+ vs MLR- | ENSMMUG00000063609 | 1.000 | 0.000 | 0.43 | 0.865 | 0.719 |
| R.279 Spleen MLR+ vs MLR- | HOPX | 1.000 | 0.000 | 0.44 | 0.757 | 0.444 |
| R.279 Spleen MLR+ vs MLR- | GSTK1 | 1.000 | 0.000 | 0.37 | 0.649 | 0.347 |
| R.279 Spleen MLR+ vs MLR- | TYMS | 1.000 | 0.000 | 0.46 | 0.243 | 0.08 |
| R.279 Spleen MLR+ vs MLR- | EED | 1.000 | 0.000 | 0.37 | 0.541 | 0.271 |
| R.279 Spleen MLR+ vs MLR- | RPL30 | 1.000 | 0.000 | -0.39 | 1 | 0.994 |
| R.279 Spleen MLR+ vs MLR- | SERPINA1 | 1.000 | 0.000 | 0.57 | 0.541 | 0.283 |
| R.279 Spleen MLR+ vs MLR- | CCNA2 | 1.000 | 0.000 | 0.31 | 0.243 | 0.079 |
| R.279 Spleen MLR+ vs MLR- | RPS29 | 1.000 | 0.000 | -0.32 | 1 | 0.993 |
| R.279 Spleen MLR+ vs MLR- | TRIM25 | 1.000 | 0.000 | 0.35 | 0.432 | 0.197 |
| R.279 Spleen MLR+ vs MLR- | ENSMMUG00000003867 | 1.000 | 0.000 | -0.37 | 1 | 0.997 |
| R.279 Spleen MLR+ vs MLR- | CRIP1 | 1.000 | 0.000 | 0.54 | 0.973 | 0.911 |
| R.279 Spleen MLR+ vs MLR- | SLC25A6 | 1.000 | 0.000 | -0.46 | 0.838 | 0.942 |
| R.279 Spleen MLR+ vs MLR- | RPL8 | 1.000 | 0.000 | -0.37 | 0.973 | 0.998 |
| R.279 Spleen MLR+ vs MLR- | AHNAK | 1.000 | 0.000 | 0.68 | 0.73 | 0.489 |
| R.279 Spleen MLR+ vs MLR- | FGFBP2 | 1.000 | 0.000 | 0.27 | 0.27 | 0.096 |
| R.279 Spleen MLR+ vs MLR- | GPS2 | 1.000 | 0.000 | 0.29 | 0.459 | 0.218 |
| R.279 Spleen MLR+ vs MLR- | RPS7 | 1.000 | 0.000 | -0.34 | 1 | 0.994 |
| R.279 Spleen MLR+ vs MLR- | ENSMMUG00000056183 | 1.000 | 0.000 | 0.26 | 0.189 | 0.056 |
| R.279 Spleen MLR+ vs MLR- | LARP7 | 1.000 | 0.000 | 0.30 | 0.378 | 0.166 |
| R.279 Spleen MLR+ vs MLR- | RPL27 | 1.000 | 0.000 | -0.36 | 1 | 0.992 |
| R.279 Spleen MLR+ vs MLR- | CBFB | 1.000 | 0.000 | 0.27 | 0.351 | 0.147 |
| R.279 Spleen MLR+ vs MLR- | RPL23A | 1.000 | 0.000 | -0.44 | 0.892 | 0.923 |
| R.279 Spleen MLR+ vs MLR- | ENSMMUG00000003130 | 1.000 | 0.000 | 0.26 | 0.514 | 0.251 |
| R.279 Spleen MLR+ vs MLR- | PRF1 | 1.000 | 0.001 | 0.37 | 0.649 | 0.359 |
| R.279 Spleen MLR+ vs MLR- | LRRC8C | 1.000 | 0.001 | 0.36 | 0.486 | 0.248 |
| R.279 Spleen MLR+ vs MLR- | RPL32 | 1.000 | 0.001 | -0.34 | 1 | 0.996 |
| R.279 Spleen MLR+ vs MLR- | LTB | 1.000 | 0.001 | -0.75 | 0.378 | 0.643 |
| R.279 Spleen MLR+ vs MLR- | RPL10 | 1.000 | 0.001 | -0.30 | 1 | 0.997 |
| R.279 Spleen MLR+ vs MLR- | RPL7A | 1.000 | 0.001 | -0.36 | 1 | 0.994 |
| R.279 Spleen MLR+ vs MLR- | RPS26 | 1.000 | 0.001 | -0.31 | 1 | 0.995 |
| R.279 Spleen MLR+ vs MLR- | RRM2 | 1.000 | 0.001 | 1.06 | 0.27 | 0.107 |
| R.279 Spleen MLR+ vs MLR- | EIF3K | 1.000 | 0.001 | -0.41 | 0.811 | 0.944 |
| R.279 Spleen MLR+ vs MLR- | SSBP4 | 1.000 | 0.001 | 0.35 | 0.622 | 0.362 |
| R.279 Spleen MLR+ vs MLR- | ETS2 | 1.000 | 0.001 | 0.40 | 0.351 | 0.153 |
| R.279 Spleen MLR+ vs MLR- | CCL4L1 | 1.000 | 0.001 | 0.63 | 0.703 | 0.417 |
| R.279 Spleen MLR+ vs MLR- | PPP1CC | 1.000 | 0.001 | -0.61 | 0.459 | 0.65 |
| R.279 Spleen MLR+ vs MLR- | RPS9 | 1.000 | 0.001 | -0.31 | 1 | 0.997 |
| R.279 Spleen MLR+ vs MLR- | ISG20 | 1.000 | 0.001 | 0.35 | 0.892 | 0.641 |
| R.279 Spleen MLR+ vs MLR- | LASP1 | 1.000 | 0.001 | 0.38 | 0.514 | 0.317 |
| R.279 Spleen MLR+ vs MLR- | PLP2 | 1.000 | 0.001 | 0.47 | 0.703 | 0.479 |
| R.279 Spleen MLR+ vs MLR- | RAP1GDS1 | 1.000 | 0.001 | 0.29 | 0.378 | 0.173 |
| R.279 Spleen MLR+ vs MLR- | ENSMMUG00000062350 | 1.000 | 0.001 | -0.28 | 1 | 0.997 |
| R.279 Spleen MLR+ vs MLR- | ID3 | 1.000 | 0.001 | -0.78 | 0.054 | 0.297 |
| R.279 Spleen MLR+ vs MLR- | HMGB2 | 1.000 | 0.001 | 0.99 | 0.676 | 0.486 |
| R.279 Spleen MLR+ vs MLR- | RPL9 | 1.000 | 0.001 | -0.34 | 1 | 0.993 |
| R.279 Spleen MLR+ vs MLR- | RPS5 | 1.000 | 0.001 | -0.33 | 1 | 0.995 |
| R.279 Spleen MLR+ vs MLR- | ENSMMUG00000005593 | 1.000 | 0.001 | -0.29 | 1 | 0.996 |
| R.279 Spleen MLR+ vs MLR- | RPL12 | 1.000 | 0.001 | -0.39 | 1 | 0.995 |
| R.279 Spleen MLR+ vs MLR- | ENSMMUG00000021023 | 1.000 | 0.001 | 0.33 | 0.189 | 0.06 |
| R.279 Spleen MLR+ vs MLR- | PRR13 | 1.000 | 0.001 | 0.36 | 0.946 | 0.698 |
| R.279 Spleen MLR+ vs MLR- | MKI67 | 1.000 | 0.001 | 0.64 | 0.405 | 0.195 |
| R.279 Spleen MLR+ vs MLR- | TRAPPC4 | 1.000 | 0.001 | 0.31 | 0.514 | 0.274 |
| R.279 Spleen MLR+ vs MLR- | NACA | 1.000 | 0.001 | -0.37 | 1 | 0.99 |
| R.279 Spleen MLR+ vs MLR- | KIF20B | 1.000 | 0.001 | 0.31 | 0.297 | 0.123 |
| R.279 Spleen MLR+ vs MLR- | GLIPR2 | 1.000 | 0.001 | 0.42 | 0.73 | 0.494 |

| | | | | | | |
|---|---|---|---|---|---|---|
| R.279 Spleen MLR+ vs MLR- | H1-5 | 1.000 | 0.001 | 0.35 | 0.162 | 0.049 |
| R.279 Spleen MLR+ vs MLR- | ND6 | 1.000 | 0.001 | -0.45 | 0.054 | 0.29 |
| R.279 Spleen MLR+ vs MLR- | GPX4 | 1.000 | 0.001 | 0.47 | 0.811 | 0.658 |
| R.279 Spleen MLR+ vs MLR- | CDKN1A | 1.000 | 0.002 | 0.48 | 0.514 | 0.298 |
| R.279 Spleen MLR+ vs MLR- | RAB27A | 1.000 | 0.002 | 0.26 | 0.432 | 0.217 |
| R.279 Spleen MLR+ vs MLR- | FLNA | 1.000 | 0.002 | 0.58 | 0.73 | 0.546 |
| R.279 Spleen MLR+ vs MLR- | RPL18 | 1.000 | 0.002 | -0.29 | 1 | 0.994 |
| R.279 Spleen MLR+ vs MLR- | RPL13A | 1.000 | 0.002 | -0.30 | 1 | 0.995 |
| R.279 Spleen MLR+ vs MLR- | IFI27L2 | 1.000 | 0.002 | 0.37 | 0.919 | 0.845 |
| R.279 Spleen MLR+ vs MLR- | ST3GAL1 | 1.000 | 0.002 | 0.38 | 0.405 | 0.212 |
| R.279 Spleen MLR+ vs MLR- | PTTG1 | 1.000 | 0.002 | 0.48 | 0.405 | 0.2 |
| R.279 Spleen MLR+ vs MLR- | UBTF | 1.000 | 0.002 | 0.27 | 0.351 | 0.166 |
| R.279 Spleen MLR+ vs MLR- | RBBP8 | 1.000 | 0.002 | 0.26 | 0.243 | 0.094 |
| R.279 Spleen MLR+ vs MLR- | RPL15 | 1.000 | 0.002 | -0.33 | 1 | 0.982 |
| R.279 Spleen MLR+ vs MLR- | AP3B1 | 1.000 | 0.002 | 0.36 | 0.378 | 0.202 |
| R.279 Spleen MLR+ vs MLR- | H1-4 | 1.000 | 0.002 | 0.57 | 0.459 | 0.262 |
| R.279 Spleen MLR+ vs MLR- | MAMU-E | 1.000 | 0.002 | 0.33 | 1 | 0.992 |
| R.279 Spleen MLR+ vs MLR- | CD99 | 1.000 | 0.002 | 0.47 | 0.703 | 0.506 |
| R.279 Spleen MLR+ vs MLR- | ENSMMUG00000016898 | 1.000 | 0.002 | 0.62 | 0.649 | 0.413 |
| R.279 Spleen MLR+ vs MLR- | CLTA | 1.000 | 0.003 | 0.42 | 0.622 | 0.436 |
| R.279 Spleen MLR+ vs MLR- | RPS28 | 1.000 | 0.003 | -0.30 | 1 | 0.993 |
| R.279 Spleen MLR+ vs MLR- | ENSMMUG00000058581 | 1.000 | 0.003 | 0.62 | 0.892 | 0.742 |
| R.279 Spleen MLR+ vs MLR- | TUBA1A | 1.000 | 0.003 | 0.76 | 0.811 | 0.805 |
| R.279 Spleen MLR+ vs MLR- | MAMU-DRA | 1.000 | 0.003 | 0.47 | 0.243 | 0.101 |
| R.279 Spleen MLR+ vs MLR- | CENPW | 1.000 | 0.003 | 0.29 | 0.216 | 0.082 |
| R.279 Spleen MLR+ vs MLR- | RPL17 | 1.000 | 0.003 | -0.35 | 1 | 0.988 |
| R.279 Spleen MLR+ vs MLR- | RPL3 | 1.000 | 0.003 | -0.28 | 1 | 0.996 |
| R.279 Spleen MLR+ vs MLR- | TRAF3IP3 | 1.000 | 0.003 | 0.30 | 0.676 | 0.423 |
| R.279 Spleen MLR+ vs MLR- | HELZ | 1.000 | 0.003 | 0.29 | 0.351 | 0.172 |
| R.279 Spleen MLR+ vs MLR- | RGS2 | 1.000 | 0.003 | -0.59 | 0.027 | 0.234 |
| R.279 Spleen MLR+ vs MLR- | NUDT3 | 1.000 | 0.003 | -0.27 | 1 | 0.994 |
| R.279 Spleen MLR+ vs MLR- | KIFC1 | 1.000 | 0.003 | 0.26 | 0.297 | 0.131 |
| R.279 Spleen MLR+ vs MLR- | LEF1 | 1.000 | 0.003 | -0.43 | 0.027 | 0.232 |
| R.279 Spleen MLR+ vs MLR- | CLEC2D | 1.000 | 0.003 | 0.32 | 0.838 | 0.673 |
| R.279 Spleen MLR+ vs MLR- | RPL27A | 1.000 | 0.003 | -0.38 | 0.946 | 0.952 |
| R.279 Spleen MLR+ vs MLR- | ENSMMUG00000022075 | 1.000 | 0.003 | -0.37 | 0.297 | 0.555 |
| R.279 Spleen MLR+ vs MLR- | CEP57 | 1.000 | 0.003 | 0.28 | 0.324 | 0.154 |
| R.279 Spleen MLR+ vs MLR- | FOS | 1.000 | 0.004 | -0.77 | 0.703 | 0.84 |
| R.279 Spleen MLR+ vs MLR- | NME1 | 1.000 | 0.004 | -0.39 | 0.838 | 0.956 |
| R.279 Spleen MLR+ vs MLR- | RPS21 | 1.000 | 0.004 | -0.26 | 1 | 0.995 |
| R.279 Spleen MLR+ vs MLR- | CSTB | 1.000 | 0.004 | 0.28 | 0.541 | 0.321 |
| R.279 Spleen MLR+ vs MLR- | RPL5 | 1.000 | 0.004 | -0.30 | 1 | 0.992 |
| R.279 Spleen MLR+ vs MLR- | TPST2 | 1.000 | 0.004 | 0.32 | 0.541 | 0.335 |
| R.279 Spleen MLR+ vs MLR- | CAPN2 | 1.000 | 0.004 | 0.26 | 0.432 | 0.235 |
| R.279 Spleen MLR+ vs MLR- | JPT1 | 1.000 | 0.004 | 0.37 | 0.946 | 0.87 |
| R.279 Spleen MLR+ vs MLR- | CCR7 | 1.000 | 0.004 | -0.59 | 0.081 | 0.299 |
| R.279 Spleen MLR+ vs MLR- | USP36 | 1.000 | 0.004 | -0.34 | 0.054 | 0.261 |
| R.279 Spleen MLR+ vs MLR- | APMAP | 1.000 | 0.004 | 0.26 | 0.351 | 0.177 |
| R.279 Spleen MLR+ vs MLR- | IPO7 | 1.000 | 0.004 | 0.26 | 0.378 | 0.198 |
| R.279 Spleen MLR+ vs MLR- | SRI | 1.000 | 0.004 | 0.32 | 0.73 | 0.501 |
| R.279 Spleen MLR+ vs MLR- | CTLA4 | 1.000 | 0.005 | -0.42 | 0 | 0.181 |
| R.279 Spleen MLR+ vs MLR- | G3BP2 | 1.000 | 0.005 | -0.50 | 0.378 | 0.582 |
| R.279 Spleen MLR+ vs MLR- | HPCAL1 | 1.000 | 0.005 | -0.32 | 0.081 | 0.291 |
| R.279 Spleen MLR+ vs MLR- | TMED10 | 1.000 | 0.005 | -0.37 | 0.189 | 0.405 |
| R.279 Spleen MLR+ vs MLR- | KLF6 | 1.000 | 0.005 | -0.44 | 0.622 | 0.851 |
| R.279 Spleen MLR+ vs MLR- | NFKB1 | 1.000 | 0.005 | -0.40 | 0.135 | 0.346 |

| | | | | | | |
|---|---|---|---|---|---|---|
| R.279 Spleen MLR+ vs MLR- | EPC1 | 1.000 | 0.005 | -0.29 | 0.054 | 0.264 |
| R.279 Spleen MLR+ vs MLR- | PCLAF | 1.000 | 0.005 | 0.36 | 0.216 | 0.086 |
| R.279 Spleen MLR+ vs MLR- | MAPK1 | 1.000 | 0.006 | 0.26 | 0.459 | 0.26 |
| R.279 Spleen MLR+ vs MLR- | SELL | 1.000 | 0.006 | -0.41 | 0.135 | 0.373 |
| R.279 Spleen MLR+ vs MLR- | PPIA | 1.000 | 0.006 | 0.33 | 1 | 0.99 |
| R.279 Spleen MLR+ vs MLR- | ENSMMUG00000017890 | 1.000 | 0.006 | -0.35 | 0.162 | 0.371 |
| R.279 Spleen MLR+ vs MLR- | MTPN | 1.000 | 0.006 | 0.31 | 0.595 | 0.395 |
| R.279 Spleen MLR+ vs MLR- | RAP1B | 1.000 | 0.007 | 0.39 | 0.811 | 0.768 |
| R.279 Spleen MLR+ vs MLR- | RNASEH2A | 1.000 | 0.007 | 0.26 | 0.216 | 0.093 |
| R.279 Spleen MLR+ vs MLR- | TGFBR3 | 1.000 | 0.007 | 0.32 | 0.486 | 0.288 |
| R.279 Spleen MLR+ vs MLR- | ENSMMUG00000013429 | 1.000 | 0.007 | -0.40 | 1 | 0.996 |
| R.279 Spleen MLR+ vs MLR- | RPA3 | 1.000 | 0.007 | 0.39 | 0.703 | 0.523 |
| R.279 Spleen MLR+ vs MLR- | PRPF6 | 1.000 | 0.007 | 0.27 | 0.459 | 0.269 |
| R.279 Spleen MLR+ vs MLR- | MT2A | 1.000 | 0.008 | 0.44 | 0.595 | 0.411 |
| R.279 Spleen MLR+ vs MLR- | SOCS3 | 1.000 | 0.008 | -0.30 | 0.081 | 0.284 |
| R.279 Spleen MLR+ vs MLR- | TRAPPC3 | 1.000 | 0.008 | 0.26 | 0.432 | 0.246 |
| R.279 Spleen MLR+ vs MLR- | MLF2 | 1.000 | 0.008 | -0.32 | 0.162 | 0.365 |
| R.279 Spleen MLR+ vs MLR- | RPS11 | 1.000 | 0.008 | -0.30 | 1 | 0.995 |
| R.279 Spleen MLR+ vs MLR- | COX7A1 | 1.000 | 0.008 | -0.25 | 0 | 0.16 |
| R.279 Spleen MLR+ vs MLR- | ANXA2 | 1.000 | 0.008 | 0.49 | 0.676 | 0.516 |
| R.279 Spleen MLR+ vs MLR- | DNAJC1 | 1.000 | 0.008 | 0.28 | 0.486 | 0.295 |
| R.279 Spleen MLR+ vs MLR- | ENSMMUG00000004633 | 1.000 | 0.008 | 0.26 | 0.243 | 0.112 |
| R.279 Spleen MLR+ vs MLR- | IFI6 | 1.000 | 0.008 | 0.26 | 0.784 | 0.619 |
| R.279 Spleen MLR+ vs MLR- | SH3KBP1 | 1.000 | 0.009 | 0.39 | 0.703 | 0.57 |
| R.279 Spleen MLR+ vs MLR- | CMTR1 | 1.000 | 0.009 | 0.27 | 0.324 | 0.166 |
| R.279 Spleen MLR+ vs MLR- | FLOT1 | 1.000 | 0.009 | 0.44 | 0.541 | 0.36 |
| R.279 Spleen MLR+ vs MLR- | RNF166 | 1.000 | 0.009 | 0.27 | 0.297 | 0.154 |
| R.279 Spleen MLR+ vs MLR- | DNMT1 | 1.000 | 0.009 | 0.43 | 0.405 | 0.247 |
| R.279 Spleen MLR+ vs MLR- | CSRNP1 | 1.000 | 0.010 | -0.44 | 0.405 | 0.581 |
| R.279 Spleen MLR+ vs MLR- | HSPA8 | 1.000 | 0.010 | 0.50 | 0.973 | 0.98 |
| R.279 Spleen MLR+ vs MLR- | GTF2I | 1.000 | 0.010 | -0.29 | 0.081 | 0.267 |
| R.279 Spleen MLR+ vs MLR- | PLCG1 | 1.000 | 0.010 | -0.26 | 0.027 | 0.195 |
| R.279 Spleen MLR+ vs MLR- | CD38 | 1.000 | 0.010 | -0.45 | 0.27 | 0.45 |
| R.279 Spleen MLR+ vs MLR- | SSU72 | 1.000 | 0.010 | -0.30 | 0.108 | 0.299 |
| R.279 Spleen MLR+ vs MLR- | EIF4EBP2 | 1.000 | 0.010 | 0.29 | 0.351 | 0.191 |
| R.279 Spleen MLR+ vs MLR- | FBL | 1.000 | 0.010 | -0.33 | 0.189 | 0.39 |
| R.279 Spleen MLR+ vs MLR- | ARPC5L | 1.000 | 0.010 | 0.31 | 0.757 | 0.561 |
| R.279 Spleen MLR+ vs MLR- | AIMP1 | 1.000 | 0.011 | -0.33 | 0.162 | 0.354 |
| R.279 Spleen MLR+ vs MLR- | INPP4B | 1.000 | 0.011 | -0.34 | 0.135 | 0.33 |
| R.279 Spleen MLR+ vs MLR- | ENSMMUG00000057791 | 1.000 | 0.011 | -0.50 | 0 | 0.15 |
| R.279 Spleen MLR+ vs MLR- | MAMU-F | 1.000 | 0.011 | 0.28 | 0.432 | 0.265 |
| R.279 Spleen MLR+ vs MLR- | MCM7 | 1.000 | 0.011 | 0.31 | 0.243 | 0.116 |
| R.279 Spleen MLR+ vs MLR- | RABAC1 | 1.000 | 0.011 | 0.38 | 0.595 | 0.425 |
| R.279 Spleen MLR+ vs MLR- | SUB1 | 1.000 | 0.011 | 0.30 | 0.865 | 0.702 |
| R.279 Spleen MLR+ vs MLR- | CA6 | 1.000 | 0.011 | -0.41 | 0.027 | 0.189 |
| R.279 Spleen MLR+ vs MLR- | CALR | 1.000 | 0.012 | -0.41 | 0.622 | 0.804 |
| R.279 Spleen MLR+ vs MLR- | TCF7 | 1.000 | 0.012 | -0.37 | 0.162 | 0.354 |
| R.279 Spleen MLR+ vs MLR- | SYTL2 | 1.000 | 0.013 | 0.37 | 0.378 | 0.228 |
| R.279 Spleen MLR+ vs MLR- | ADSL | 1.000 | 0.013 | -0.30 | 0.081 | 0.258 |
| R.279 Spleen MLR+ vs MLR- | SDHAF2 | 1.000 | 0.013 | -0.27 | 0.081 | 0.258 |
| R.279 Spleen MLR+ vs MLR- | DGKA | 1.000 | 0.013 | -0.30 | 0.108 | 0.296 |
| R.279 Spleen MLR+ vs MLR- | WBP11 | 1.000 | 0.013 | -0.29 | 0.108 | 0.289 |
| R.279 Spleen MLR+ vs MLR- | SLC2A3 | 1.000 | 0.014 | -0.57 | 0.405 | 0.559 |
| R.279 Spleen MLR+ vs MLR- | LCP1 | 1.000 | 0.014 | 0.28 | 0.838 | 0.767 |
| R.279 Spleen MLR+ vs MLR- | LAG3 | 1.000 | 0.014 | -0.50 | 0.081 | 0.255 |
| R.279 Spleen MLR+ vs MLR- | SH2D2A | 1.000 | 0.014 | 0.32 | 0.514 | 0.35 |

| | | | | | | |
|---|---|---|---|---|---|---|
| R.279 Spleen MLR+ vs MLR- | CKS2 | 1.000 | 0.014 | 0.29 | 0.541 | 0.354 |
| R.279 Spleen MLR+ vs MLR- | RBM39 | 1.000 | 0.014 | 0.30 | 0.919 | 0.775 |
| R.279 Spleen MLR+ vs MLR- | ALDOA | 1.000 | 0.015 | -0.38 | 0.784 | 0.836 |
| R.279 Spleen MLR+ vs MLR- | CASP3 | 1.000 | 0.015 | -0.30 | 0.135 | 0.322 |
| R.279 Spleen MLR+ vs MLR- | SKAP1 | 1.000 | 0.015 | 0.26 | 0.865 | 0.651 |
| R.279 Spleen MLR+ vs MLR- | PSMC5 | 1.000 | 0.015 | -0.28 | 0.189 | 0.383 |
| R.279 Spleen MLR+ vs MLR- | APOBEC3G | 1.000 | 0.016 | 0.30 | 0.541 | 0.362 |
| R.279 Spleen MLR+ vs MLR- | RRM1 | 1.000 | 0.016 | 0.29 | 0.351 | 0.201 |
| R.279 Spleen MLR+ vs MLR- | DUSP2 | 1.000 | 0.016 | -0.63 | 0.378 | 0.53 |
| R.279 Spleen MLR+ vs MLR- | MRPL23 | 1.000 | 0.016 | -0.33 | 0.351 | 0.551 |
| R.279 Spleen MLR+ vs MLR- | ICOS | 1.000 | 0.016 | -0.43 | 0.135 | 0.31 |
| R.279 Spleen MLR+ vs MLR- | MBNL1 | 1.000 | 0.016 | -0.32 | 0.595 | 0.762 |
| R.279 Spleen MLR+ vs MLR- | ATP1A1 | 1.000 | 0.017 | 0.30 | 0.486 | 0.326 |
| R.279 Spleen MLR+ vs MLR- | SFRP5 | 1.000 | 0.017 | -0.30 | 0 | 0.134 |
| R.279 Spleen MLR+ vs MLR- | CCNI | 1.000 | 0.017 | -0.35 | 0.378 | 0.552 |
| R.279 Spleen MLR+ vs MLR- | SERPINB1 | 1.000 | 0.017 | 0.38 | 0.486 | 0.343 |
| R.279 Spleen MLR+ vs MLR- | ENSMMUG00000002320 | 1.000 | 0.017 | -0.36 | 0.946 | 0.986 |
| R.279 Spleen MLR+ vs MLR- | ENSMMUG00000008604 | 1.000 | 0.018 | -0.26 | 0.054 | 0.213 |
| R.279 Spleen MLR+ vs MLR- | ENSMMUG00000006283 | 1.000 | 0.018 | -0.34 | 0.838 | 0.81 |
| R.279 Spleen MLR+ vs MLR- | CD164 | 1.000 | 0.018 | -0.32 | 0.432 | 0.608 |
| R.279 Spleen MLR+ vs MLR- | FOSB | 1.000 | 0.018 | -0.54 | 0.622 | 0.772 |
| R.279 Spleen MLR+ vs MLR- | CYLD | 1.000 | 0.018 | -0.27 | 0.189 | 0.386 |
| R.279 Spleen MLR+ vs MLR- | UBE2F | 1.000 | 0.018 | -0.27 | 0.054 | 0.212 |
| R.279 Spleen MLR+ vs MLR- | YWHAB | 1.000 | 0.018 | 0.28 | 0.892 | 0.749 |
| R.279 Spleen MLR+ vs MLR- | PIM2 | 1.000 | 0.018 | -0.27 | 0.054 | 0.211 |
| R.279 Spleen MLR+ vs MLR- | LENG8 | 1.000 | 0.018 | -0.26 | 0.081 | 0.247 |
| R.279 Spleen MLR+ vs MLR- | CENPM | 1.000 | 0.019 | 0.31 | 0.216 | 0.099 |
| R.279 Spleen MLR+ vs MLR- | RCSD1 | 1.000 | 0.019 | -0.29 | 0.162 | 0.345 |
| R.279 Spleen MLR+ vs MLR- | GALNS | 1.000 | 0.019 | -0.35 | 0.378 | 0.55 |
| R.279 Spleen MLR+ vs MLR- | STK10 | 1.000 | 0.019 | 0.25 | 0.514 | 0.339 |
| R.279 Spleen MLR+ vs MLR- | ENSMMUG00000012140 | 1.000 | 0.020 | 0.30 | 1 | 0.997 |
| R.279 Spleen MLR+ vs MLR- | COMMD6 | 1.000 | 0.020 | -0.33 | 0.568 | 0.733 |
| R.279 Spleen MLR+ vs MLR- | CASP4 | 1.000 | 0.021 | -0.25 | 0.081 | 0.242 |
| R.279 Spleen MLR+ vs MLR- | STING1 | 1.000 | 0.021 | -0.27 | 0.054 | 0.206 |
| R.279 Spleen MLR+ vs MLR- | ENSMMUG00000003412 | 1.000 | 0.021 | -0.26 | 1 | 0.995 |
| R.279 Spleen MLR+ vs MLR- | PRRC2C | 1.000 | 0.021 | -0.32 | 0.486 | 0.64 |
| R.279 Spleen MLR+ vs MLR- | CEMIP2 | 1.000 | 0.021 | -0.25 | 0.162 | 0.35 |
| R.279 Spleen MLR+ vs MLR- | ORAI1 | 1.000 | 0.022 | 0.29 | 0.622 | 0.465 |
| R.279 Spleen MLR+ vs MLR- | CD74 | 1.000 | 0.022 | 0.52 | 0.703 | 0.602 |
| R.279 Spleen MLR+ vs MLR- | USP1 | 1.000 | 0.022 | 0.28 | 0.378 | 0.229 |
| R.279 Spleen MLR+ vs MLR- | PELI1 | 1.000 | 0.022 | -0.38 | 0.216 | 0.375 |
| R.279 Spleen MLR+ vs MLR- | CD28 | 1.000 | 0.022 | -0.28 | 0.054 | 0.204 |
| R.279 Spleen MLR+ vs MLR- | PM20D2 | 1.000 | 0.022 | -0.29 | 0.676 | 0.847 |
| R.279 Spleen MLR+ vs MLR- | BIRC5 | 1.000 | 0.022 | 0.26 | 0.162 | 0.07 |
| R.279 Spleen MLR+ vs MLR- | DGKZ | 1.000 | 0.022 | 0.26 | 0.568 | 0.399 |
| R.279 Spleen MLR+ vs MLR- | IL10RA | 1.000 | 0.022 | 0.30 | 0.568 | 0.398 |
| R.279 Spleen MLR+ vs MLR- | ENSMMUG00000056515 | 1.000 | 0.023 | -0.83 | 0 | 0.124 |
| R.279 Spleen MLR+ vs MLR- | TOP2A | 1.000 | 0.023 | 0.28 | 0.189 | 0.084 |
| R.279 Spleen MLR+ vs MLR- | TUBB | 1.000 | 0.023 | 0.78 | 0.622 | 0.521 |
| R.279 Spleen MLR+ vs MLR- | EEF1G | 1.000 | 0.023 | -0.31 | 0.865 | 0.894 |
| R.279 Spleen MLR+ vs MLR- | RAB1B | 1.000 | 0.023 | 0.34 | 0.595 | 0.441 |
| R.279 Spleen MLR+ vs MLR- | ENSMMUG00000051392 | 1.000 | 0.023 | 0.58 | 0.595 | 0.519 |
| R.279 Spleen MLR+ vs MLR- | PHIP | 1.000 | 0.023 | 0.29 | 0.405 | 0.267 |
| R.279 Spleen MLR+ vs MLR- | HBP1 | 1.000 | 0.023 | -0.26 | 0.135 | 0.31 |
| R.279 Spleen MLR+ vs MLR- | JUNB | 1.000 | 0.024 | -0.42 | 0.405 | 0.595 |
| R.279 Spleen MLR+ vs MLR- | ENSMMUG00000065238 | 1.000 | 0.024 | 0.55 | 0.216 | 0.107 |

| | | | | | | |
|---|---|---|---|---|---|---|
| R.279 Spleen MLR+ vs MLR- | TBC1D10B | 1.000 | 0.024 | -0.35 | 0.541 | 0.646 |
| R.279 Spleen MLR+ vs MLR- | BHLHE40 | 1.000 | 0.024 | 0.33 | 0.784 | 0.598 |
| R.279 Spleen MLR+ vs MLR- | NDFIP1 | 1.000 | 0.025 | -0.32 | 0.405 | 0.574 |
| R.279 Spleen MLR+ vs MLR- | CUTA | 1.000 | 0.025 | -0.32 | 0.351 | 0.572 |
| R.279 Spleen MLR+ vs MLR- | FOSL2 | 1.000 | 0.025 | -0.31 | 0.135 | 0.295 |
| R.279 Spleen MLR+ vs MLR- | BIRC2 | 1.000 | 0.025 | -0.30 | 0.189 | 0.354 |
| R.279 Spleen MLR+ vs MLR- | CD83 | 1.000 | 0.026 | -0.61 | 0.324 | 0.479 |
| R.279 Spleen MLR+ vs MLR- | FTH1 | 1.000 | 0.027 | -0.37 | 0.838 | 0.918 |
| R.279 Spleen MLR+ vs MLR- | APOL2 | 1.000 | 0.027 | 0.26 | 0.622 | 0.428 |
| R.279 Spleen MLR+ vs MLR- | UBE2B | 1.000 | 0.028 | 0.30 | 0.622 | 0.475 |
| R.279 Spleen MLR+ vs MLR- | KLF3 | 1.000 | 0.028 | 0.28 | 0.378 | 0.241 |
| R.279 Spleen MLR+ vs MLR- | SPCS2 | 1.000 | 0.028 | 0.30 | 0.541 | 0.376 |
| R.279 Spleen MLR+ vs MLR- | KIFBP | 1.000 | 0.029 | -0.25 | 0.162 | 0.347 |
| R.279 Spleen MLR+ vs MLR- | EGLN3 | 1.000 | 0.029 | -0.27 | 0.054 | 0.195 |
| R.279 Spleen MLR+ vs MLR- | ALYREF | 1.000 | 0.029 | 0.36 | 0.541 | 0.386 |
| R.279 Spleen MLR+ vs MLR- | DEK | 1.000 | 0.030 | 0.37 | 0.622 | 0.47 |
| R.279 Spleen MLR+ vs MLR- | NR4A2 | 1.000 | 0.030 | -0.51 | 0.703 | 0.774 |
| R.279 Spleen MLR+ vs MLR- | H2AFX | 1.000 | 0.030 | 0.53 | 0.486 | 0.336 |
| R.279 Spleen MLR+ vs MLR- | SPN | 1.000 | 0.030 | 0.28 | 0.514 | 0.351 |
| R.279 Spleen MLR+ vs MLR- | ENSMMUG00000002324 | 1.000 | 0.031 | -0.28 | 0.189 | 0.347 |
| R.279 Spleen MLR+ vs MLR- | ETS1 | 1.000 | 0.031 | -0.30 | 0.649 | 0.77 |
| R.279 Spleen MLR+ vs MLR- | ENSMMUG00000064232 | 1.000 | 0.032 | -0.27 | 0.162 | 0.325 |
| R.279 Spleen MLR+ vs MLR- | STMN1 | 1.000 | 0.033 | 1.16 | 0.459 | 0.348 |
| R.279 Spleen MLR+ vs MLR- | LGALS3 | 1.000 | 0.033 | 0.26 | 0.703 | 0.499 |
| R.279 Spleen MLR+ vs MLR- | ATF2 | 1.000 | 0.034 | -0.27 | 0.135 | 0.287 |
| R.279 Spleen MLR+ vs MLR- | ENSMMUG00000060689 | 1.000 | 0.034 | 0.29 | 0.108 | 0.039 |
| R.279 Spleen MLR+ vs MLR- | REL | 1.000 | 0.034 | -0.35 | 0.378 | 0.529 |
| R.279 Spleen MLR+ vs MLR- | CD3G | 1.000 | 0.036 | 0.29 | 0.811 | 0.769 |
| R.279 Spleen MLR+ vs MLR- | UCP2 | 1.000 | 0.037 | 0.46 | 0.757 | 0.694 |
| R.279 Spleen MLR+ vs MLR- | HSD17B11 | 1.000 | 0.037 | 0.31 | 0.405 | 0.269 |
| R.279 Spleen MLR+ vs MLR- | ATP5F1C | 1.000 | 0.038 | -0.29 | 0.351 | 0.496 |
| R.279 Spleen MLR+ vs MLR- | GPRIN3 | 1.000 | 0.038 | -0.25 | 0.108 | 0.252 |
| R.279 Spleen MLR+ vs MLR- | SRSF5 | 1.000 | 0.038 | -0.27 | 0.595 | 0.722 |
| R.279 Spleen MLR+ vs MLR- | ID2 | 1.000 | 0.039 | -0.48 | 0.514 | 0.65 |
| R.279 Spleen MLR+ vs MLR- | B3GNT2 | 1.000 | 0.039 | -0.30 | 0.216 | 0.362 |
| R.279 Spleen MLR+ vs MLR- | MVP | 1.000 | 0.040 | 0.27 | 0.541 | 0.385 |
| R.279 Spleen MLR+ vs MLR- | ARF4 | 1.000 | 0.041 | -0.27 | 0.459 | 0.607 |
| R.279 Spleen MLR+ vs MLR- | VIM | 1.000 | 0.041 | 0.37 | 0.946 | 0.91 |
| R.279 Spleen MLR+ vs MLR- | RPL31 | 1.000 | 0.041 | -0.32 | 0.514 | 0.604 |
| R.279 Spleen MLR+ vs MLR- | TALDO1 | 1.000 | 0.042 | -0.27 | 0.324 | 0.478 |
| R.279 Spleen MLR+ vs MLR- | ATP5MJ | 1.000 | 0.042 | 0.25 | 0.973 | 0.843 |
| R.279 Spleen MLR+ vs MLR- | MAGEH1 | 1.000 | 0.043 | -0.33 | 0.108 | 0.245 |
| R.279 Spleen MLR+ vs MLR- | RBM38 | 1.000 | 0.043 | -0.38 | 0.459 | 0.571 |
| R.279 Spleen MLR+ vs MLR- | ZFP36 | 1.000 | 0.043 | -0.40 | 0.865 | 0.913 |
| R.279 Spleen MLR+ vs MLR- | PECAM1 | 1.000 | 0.044 | -0.25 | 0.135 | 0.283 |
| R.279 Spleen MLR+ vs MLR- | RNF167 | 1.000 | 0.044 | 0.29 | 0.973 | 0.811 |
| R.279 Spleen MLR+ vs MLR- | RGS16 | 1.000 | 0.045 | 0.47 | 0.243 | 0.143 |
| R.279 Spleen MLR+ vs MLR- | HIST1H2AE | 1.000 | 0.046 | 0.36 | 0.216 | 0.112 |
| R.279 Spleen MLR+ vs MLR- | CCDC82 | 1.000 | 0.046 | 0.32 | 0.405 | 0.278 |
| R.279 Spleen MLR+ vs MLR- | MIF | 1.000 | 0.046 | -0.33 | 0.595 | 0.735 |
| R.279 Spleen MLR+ vs MLR- | EIF3E | 1.000 | 0.046 | -0.29 | 0.649 | 0.715 |
| R.279 Spleen MLR+ vs MLR- | PIM3 | 1.000 | 0.046 | -0.30 | 0.108 | 0.242 |
| R.279 Spleen MLR+ vs MLR- | MGST3 | 1.000 | 0.047 | 0.26 | 0.459 | 0.334 |
| R.279 Spleen MLR+ vs MLR- | TAF15 | 1.000 | 0.049 | 0.35 | 0.568 | 0.418 |
| R.279 Spleen MLR+ vs MLR- | TKT | 1.000 | 0.052 | -0.25 | 0.243 | 0.39 |
| R.279 Spleen MLR+ vs MLR- | ENSMMUG00000006499 | 1.000 | 0.053 | -0.27 | 0.216 | 0.354 |

| | | | | | | |
|---|---|---|---|---|---|---|
| R.279 Spleen MLR+ vs MLR- | ENSMMUG00000017466 | 1.000 | 0.053 | 0.35 | 0.784 | 0.684 |
| R.279 Spleen MLR+ vs MLR- | GNB2 | 1.000 | 0.054 | 0.27 | 0.676 | 0.582 |
| R.279 Spleen MLR+ vs MLR- | HNRNPAB | 1.000 | 0.054 | 0.38 | 0.541 | 0.444 |
| R.279 Spleen MLR+ vs MLR- | RPS27L | 1.000 | 0.054 | 0.47 | 0.595 | 0.557 |
| R.279 Spleen MLR+ vs MLR- | CNN2 | 1.000 | 0.054 | -0.30 | 0.514 | 0.613 |
| R.279 Spleen MLR+ vs MLR- | LSM4 | 1.000 | 0.054 | -0.26 | 0.27 | 0.42 |
| R.279 Spleen MLR+ vs MLR- | ACAP1 | 1.000 | 0.054 | -0.27 | 0.324 | 0.458 |
| R.279 Spleen MLR+ vs MLR- | ATF3 | 1.000 | 0.055 | 0.30 | 0.405 | 0.264 |
| R.279 Spleen MLR+ vs MLR- | TOMM20 | 1.000 | 0.056 | -0.32 | 0.541 | 0.634 |
| R.279 Spleen MLR+ vs MLR- | ENSMMUG00000050862 | 1.000 | 0.058 | -0.64 | 0.081 | 0.199 |
| R.279 Spleen MLR+ vs MLR- | TAF7 | 1.000 | 0.059 | 0.26 | 0.568 | 0.437 |
| R.279 Spleen MLR+ vs MLR- | LMNB1 | 1.000 | 0.059 | 0.26 | 0.297 | 0.186 |
| R.279 Spleen MLR+ vs MLR- | AIP | 1.000 | 0.061 | -0.25 | 0.351 | 0.492 |
| R.279 Spleen MLR+ vs MLR- | TMEM123 | 1.000 | 0.065 | -0.28 | 0.324 | 0.461 |
| R.279 Spleen MLR+ vs MLR- | HNRNPDL | 1.000 | 0.065 | -0.26 | 0.838 | 0.855 |
| R.279 Spleen MLR+ vs MLR- | TMEM258 | 1.000 | 0.068 | 0.30 | 0.784 | 0.681 |
| R.279 Spleen MLR+ vs MLR- | NR4A3 | 1.000 | 0.068 | -0.30 | 0.162 | 0.284 |
| R.279 Spleen MLR+ vs MLR- | STX11 | 1.000 | 0.072 | 0.27 | 0.459 | 0.336 |
| R.279 Spleen MLR+ vs MLR- | ARHGAP9 | 1.000 | 0.073 | -0.28 | 0.297 | 0.426 |
| R.279 Spleen MLR+ vs MLR- | ENSMMUG00000061128 | 1.000 | 0.073 | 0.33 | 0.622 | 0.504 |
| R.279 Spleen MLR+ vs MLR- | TIGAR | 1.000 | 0.075 | -0.26 | 0.189 | 0.327 |
| R.279 Spleen MLR+ vs MLR- | CYTIP | 1.000 | 0.077 | 0.27 | 0.622 | 0.534 |
| R.279 Spleen MLR+ vs MLR- | SLC9A3R1 | 1.000 | 0.081 | 0.26 | 0.622 | 0.481 |
| R.279 Spleen MLR+ vs MLR- | PPDPF | 1.000 | 0.086 | 0.31 | 0.622 | 0.5 |
| R.279 Spleen MLR+ vs MLR- | LPXN | 1.000 | 0.087 | -0.25 | 0.27 | 0.383 |
| R.279 Spleen MLR+ vs MLR- | SAP18 | 1.000 | 0.087 | 0.26 | 0.703 | 0.618 |
| R.279 Spleen MLR+ vs MLR- | HOOK3 | 1.000 | 0.090 | 0.26 | 0.459 | 0.364 |
| R.279 Spleen MLR+ vs MLR- | PSMB10 | 1.000 | 0.092 | -0.26 | 0.351 | 0.462 |
| R.279 Spleen MLR+ vs MLR- | CXCR4 | 1.000 | 0.092 | 0.34 | 0.838 | 0.726 |
| R.279 Spleen MLR+ vs MLR- | NMT1 | 1.000 | 0.094 | 0.33 | 0.27 | 0.184 |
| R.279 Spleen MLR+ vs MLR- | RNASEH2B | 1.000 | 0.096 | 0.29 | 0.324 | 0.234 |
| R.279 Spleen MLR+ vs MLR- | PCNA | 1.000 | 0.103 | 0.30 | 0.459 | 0.335 |
| R.279 Spleen MLR+ vs MLR- | TYROBP | 1.000 | 0.106 | -0.31 | 0.054 | 0.15 |
| R.279 Spleen MLR+ vs MLR- | C11H12orf57 | 1.000 | 0.106 | -0.26 | 0.757 | 0.785 |
| R.279 Spleen MLR+ vs MLR- | LIMD2 | 1.000 | 0.106 | -0.27 | 0.595 | 0.648 |
| R.279 Spleen MLR+ vs MLR- | CRTAM | 1.000 | 0.106 | -0.41 | 0.216 | 0.317 |
| R.279 Spleen MLR+ vs MLR- | DNAJB1 | 1.000 | 0.108 | 0.32 | 0.459 | 0.339 |
| R.279 Spleen MLR+ vs MLR- | H2AFZ | 1.000 | 0.109 | 0.43 | 0.703 | 0.633 |
| R.279 Spleen MLR+ vs MLR- | HMGB1 | 1.000 | 0.110 | 0.51 | 0.865 | 0.78 |
| R.279 Spleen MLR+ vs MLR- | ND4L | 1.000 | 0.113 | -0.49 | 0.027 | 0.11 |
| R.279 Spleen MLR+ vs MLR- | ENSMMUG00000063316 | 1.000 | 0.114 | -0.29 | 0.054 | 0.142 |
| R.279 Spleen MLR+ vs MLR- | COX2 | 1.000 | 0.116 | -0.46 | 0.892 | 0.95 |
| R.279 Spleen MLR+ vs MLR- | EIF1AX | 1.000 | 0.120 | -0.25 | 0.351 | 0.454 |
| R.279 Spleen MLR+ vs MLR- | GPCPD1 | 1.000 | 0.121 | -0.29 | 0.378 | 0.463 |
| R.279 Spleen MLR+ vs MLR- | JUN | 1.000 | 0.126 | -0.30 | 0.73 | 0.825 |
| R.279 Spleen MLR+ vs MLR- | HNRNPA3 | 1.000 | 0.141 | 0.39 | 0.649 | 0.576 |
| R.279 Spleen MLR+ vs MLR- | XCL1 | 1.000 | 0.153 | -0.55 | 0.054 | 0.133 |
| R.279 Spleen MLR+ vs MLR- | TMPO | 1.000 | 0.153 | 0.31 | 0.351 | 0.283 |
| R.279 Spleen MLR+ vs MLR- | NR4A1 | 1.000 | 0.161 | -0.25 | 0.243 | 0.351 |
| R.279 Spleen MLR+ vs MLR- | ENSMMUG00000060751 | 1.000 | 0.163 | 0.29 | 0.162 | 0.098 |
| R.279 Spleen MLR+ vs MLR- | GSTP1 | 1.000 | 0.163 | -0.27 | 0.784 | 0.878 |
| R.279 Spleen MLR+ vs MLR- | SELPLG | 1.000 | 0.169 | 0.31 | 0.459 | 0.38 |
| R.279 Spleen MLR+ vs MLR- | HSP90AA1 | 1.000 | 0.171 | 0.28 | 0.892 | 0.901 |
| R.279 Spleen MLR+ vs MLR- | YPEL5 | 1.000 | 0.171 | -0.27 | 0.622 | 0.651 |
| R.279 Spleen MLR+ vs MLR- | H1-3 | 1.000 | 0.182 | 0.44 | 0.189 | 0.123 |
| R.279 Spleen MLR+ vs MLR- | LUC7L3 | 1.000 | 0.198 | 0.27 | 0.27 | 0.194 |

| | | | | | | |
|---|---|---|---|---|---|---|
| R.279 Spleen MLR+ vs MLR- | ZFP36L1 | 1.000 | 0.199 | -0.30 | 0.541 | 0.611 |
| R.279 Spleen MLR+ vs MLR- | S100A6 | 1.000 | 0.203 | -0.29 | 0.595 | 0.691 |
| R.279 Spleen MLR+ vs MLR- | CCL3 | 1.000 | 0.203 | -0.27 | 0.081 | 0.152 |
| R.279 Spleen MLR+ vs MLR- | IL2RB | 1.000 | 0.227 | -0.27 | 0.405 | 0.48 |
| R.279 Spleen MLR+ vs MLR- | ENSMMUG00000028672 | 1.000 | 0.240 | -0.28 | 0.568 | 0.616 |
| R.279 Spleen MLR+ vs MLR- | BCL2A1 | 1.000 | 0.321 | 0.26 | 0.649 | 0.649 |
| R.279 Spleen MLR+ vs MLR- | RGS1 | 1.000 | 0.330 | -0.27 | 0.405 | 0.457 |
| R.279 Spleen MLR+ vs MLR- | TSC22D3 | 1.000 | 0.333 | 0.26 | 0.865 | 0.818 |
| R.279 Spleen MLR+ vs MLR- | IFI27 | 1.000 | 0.354 | 0.25 | 0.622 | 0.563 |
| R.279 Spleen MLR+ vs MLR- | CHORDC1 | 1.000 | 0.359 | 0.30 | 0.324 | 0.271 |
| R.279 Spleen MLR+ vs MLR- | ENSMMUG00000055690 | 1.000 | 0.363 | -0.38 | 1 | 0.998 |
| R.279 Spleen MLR+ vs MLR- | MYH9 | 1.000 | 0.369 | 0.37 | 0.73 | 0.705 |
| R.279 Spleen MLR+ vs MLR- | RGCC | 1.000 | 0.390 | -0.29 | 0.73 | 0.789 |
| R.279 Spleen MLR+ vs MLR- | TXNIP | 1.000 | 0.398 | 0.31 | 0.432 | 0.426 |
| R.279 Spleen MLR+ vs MLR- | ANP32B | 1.000 | 0.416 | 0.31 | 0.568 | 0.583 |
| R.279 Spleen MLR+ vs MLR- | LMNA | 1.000 | 0.447 | -0.33 | 0.351 | 0.385 |
| R.279 Spleen MLR+ vs MLR- | GZMA | 1.000 | 0.502 | -0.67 | 0.405 | 0.314 |
| R.279 Spleen MLR+ vs MLR- | TUBA4A | 1.000 | 0.505 | -0.34 | 0.703 | 0.624 |
| R.279 Spleen MLR+ vs MLR- | CD69 | 1.000 | 0.789 | 0.30 | 0.73 | 0.764 |
| R.279 Spleen MLR+ vs MLR- | ENSMMUG00000062894 | 1.000 | 0.844 | -0.28 | 0.73 | 0.701 |
| R.279 Spleen MLR+ vs MLR- | ENSMMUG00000062077 | 1.000 | 0.868 | -1.00 | 1 | 0.998 |
| R.319 Spleen MLR+ vs MLR- | ENSMMUG00000020332 | 0.000 | 0.000 | 1.37 | 0.231 | 0.008 |
| R.319 Spleen MLR+ vs MLR- | KIR3DL12 | 0.000 | 0.000 | 0.38 | 0.282 | 0.015 |
| R.319 Spleen MLR+ vs MLR- | ENSMMUG00000050862 | 0.000 | 0.000 | 1.70 | 0.641 | 0.099 |
| R.319 Spleen MLR+ vs MLR- | TYROBP | 0.000 | 0.000 | 1.48 | 0.59 | 0.104 |
| R.319 Spleen MLR+ vs MLR- | CCL5 | 0.000 | 0.000 | 2.06 | 0.769 | 0.187 |
| R.319 Spleen MLR+ vs MLR- | NKG7 | 0.000 | 0.000 | 1.91 | 0.897 | 0.314 |
| R.319 Spleen MLR+ vs MLR- | VEGFA | 0.000 | 0.000 | 0.35 | 0.231 | 0.021 |
| R.319 Spleen MLR+ vs MLR- | ITGAX | 0.000 | 0.000 | 0.78 | 0.538 | 0.106 |
| R.319 Spleen MLR+ vs MLR- | CST7 | 0.000 | 0.000 | 1.36 | 0.872 | 0.331 |
| R.319 Spleen MLR+ vs MLR- | ENSMMUG00000056183 | 0.000 | 0.000 | 0.44 | 0.333 | 0.047 |
| R.319 Spleen MLR+ vs MLR- | ZEB2 | 0.000 | 0.000 | 0.59 | 0.436 | 0.08 |
| R.319 Spleen MLR+ vs MLR- | ENSMMUG00000063583 | 0.000 | 0.000 | 1.46 | 0.846 | 0.31 |
| R.319 Spleen MLR+ vs MLR- | NCR1 | 0.000 | 0.000 | 0.37 | 0.231 | 0.025 |
| R.319 Spleen MLR+ vs MLR- | GZMB | 0.000 | 0.000 | 1.27 | 0.795 | 0.244 |
| R.319 Spleen MLR+ vs MLR- | CTSW | 0.000 | 0.000 | 1.29 | 0.667 | 0.222 |
| R.319 Spleen MLR+ vs MLR- | MAMU-DRB1 | 0.000 | 0.000 | 0.71 | 0.41 | 0.085 |
| R.319 Spleen MLR+ vs MLR- | CCL4L1 | 0.000 | 0.000 | 1.00 | 0.615 | 0.178 |
| R.319 Spleen MLR+ vs MLR- | KLRD1 | 0.000 | 0.000 | 0.56 | 0.513 | 0.123 |
| R.319 Spleen MLR+ vs MLR- | EOMES | 0.000 | 0.000 | 0.37 | 0.308 | 0.053 |
| R.319 Spleen MLR+ vs MLR- | B2M | 0.000 | 0.000 | 0.46 | 1 | 1 |
| R.319 Spleen MLR+ vs MLR- | RPL8 | 0.000 | 0.000 | -0.59 | 1 | 1 |
| R.319 Spleen MLR+ vs MLR- | HCST | 0.000 | 0.000 | 0.95 | 0.974 | 0.662 |
| R.319 Spleen MLR+ vs MLR- | ENSMMUG00000064139 | 0.000 | 0.000 | 0.67 | 1 | 0.998 |
| R.319 Spleen MLR+ vs MLR- | RPS8 | 0.000 | 0.000 | -0.61 | 1 | 1 |
| R.319 Spleen MLR+ vs MLR- | GZMM | 0.000 | 0.000 | 1.18 | 0.564 | 0.198 |
| R.319 Spleen MLR+ vs MLR- | ENSMMUG00000058581 | 0.000 | 0.000 | 1.07 | 0.667 | 0.269 |
| R.319 Spleen MLR+ vs MLR- | EEF1A1 | 0.000 | 0.000 | -0.54 | 1 | 1 |
| R.319 Spleen MLR+ vs MLR- | ENSMMUG00000003532 | 0.000 | 0.000 | 0.62 | 0.641 | 0.211 |
| R.319 Spleen MLR+ vs MLR- | HOPX | 0.000 | 0.000 | 0.65 | 0.667 | 0.23 |
| R.319 Spleen MLR+ vs MLR- | ALOX5AP | 0.000 | 0.000 | 0.52 | 0.487 | 0.144 |
| R.319 Spleen MLR+ vs MLR- | MAMU-A | 0.000 | 0.000 | 0.43 | 1 | 1 |
| R.319 Spleen MLR+ vs MLR- | MYRF | 0.000 | 0.000 | 0.35 | 0.231 | 0.039 |
| R.319 Spleen MLR+ vs MLR- | RPL28 | 0.000 | 0.000 | -0.61 | 1 | 1 |
| R.319 Spleen MLR+ vs MLR- | LTB | 0.000 | 0.000 | -1.22 | 0.513 | 0.842 |
| R.319 Spleen MLR+ vs MLR- | RPS4X | 0.000 | 0.000 | -0.55 | 1 | 1 |

| | | | | | | |
|---|---|---|---|---|---|---|
| R.319 Spleen MLR+ vs MLR- | ADAMTS7 | 0.000 | 0.000 | 0.27 | 0.231 | 0.041 |
| R.319 Spleen MLR+ vs MLR- | CD160 | 0.000 | 0.000 | 0.44 | 0.205 | 0.034 |
| R.319 Spleen MLR+ vs MLR- | RPL10 | 0.000 | 0.000 | -0.50 | 1 | 1 |
| R.319 Spleen MLR+ vs MLR- | GZMK | 0.000 | 0.000 | 0.54 | 0.436 | 0.125 |
| R.319 Spleen MLR+ vs MLR- | PLEK | 0.000 | 0.000 | 0.35 | 0.256 | 0.052 |
| R.319 Spleen MLR+ vs MLR- | ENSMMUG00000005593 | 0.000 | 0.000 | -0.54 | 1 | 1 |
| R.319 Spleen MLR+ vs MLR- | SPOCK2 | 0.000 | 0.000 | -0.90 | 0.256 | 0.734 |
| R.319 Spleen MLR+ vs MLR- | RACK1 | 0.001 | 0.000 | -0.57 | 1 | 1 |
| R.319 Spleen MLR+ vs MLR- | FGR | 0.001 | 0.000 | 0.34 | 0.205 | 0.037 |
| R.319 Spleen MLR+ vs MLR- | ENSMMUG00000064120 | 0.001 | 0.000 | 0.53 | 1 | 0.946 |
| R.319 Spleen MLR+ vs MLR- | SCML4 | 0.002 | 0.000 | 0.67 | 0.59 | 0.257 |
| R.319 Spleen MLR+ vs MLR- | ETS2 | 0.002 | 0.000 | 0.43 | 0.41 | 0.128 |
| R.319 Spleen MLR+ vs MLR- | ZC3H10 | 0.003 | 0.000 | -0.62 | 0.897 | 0.979 |
| R.319 Spleen MLR+ vs MLR- | GCN1 | 0.003 | 0.000 | -0.48 | 1 | 1 |
| R.319 Spleen MLR+ vs MLR- | RPS3A | 0.003 | 0.000 | -0.50 | 1 | 1 |
| R.319 Spleen MLR+ vs MLR- | RGS9 | 0.003 | 0.000 | 0.58 | 0.359 | 0.111 |
| R.319 Spleen MLR+ vs MLR- | RPL22L1 | 0.003 | 0.000 | -0.91 | 0.436 | 0.789 |
| R.319 Spleen MLR+ vs MLR- | RPL7A | 0.005 | 0.000 | -0.49 | 1 | 1 |
| R.319 Spleen MLR+ vs MLR- | RPL12 | 0.006 | 0.000 | -0.56 | 1 | 1 |
| R.319 Spleen MLR+ vs MLR- | ENSMMUG00000052609 | 0.006 | 0.000 | -0.48 | 1 | 1 |
| R.319 Spleen MLR+ vs MLR- | EEF1G | 0.007 | 0.000 | -0.69 | 0.846 | 0.937 |
| R.319 Spleen MLR+ vs MLR- | CCR7 | 0.008 | 0.000 | -0.95 | 0.103 | 0.544 |
| R.319 Spleen MLR+ vs MLR- | ENSMMUG00000014786 | 0.008 | 0.000 | -0.47 | 1 | 1 |
| R.319 Spleen MLR+ vs MLR- | SKAP2 | 0.008 | 0.000 | 0.39 | 0.308 | 0.084 |
| R.319 Spleen MLR+ vs MLR- | KLRG1 | 0.010 | 0.000 | 0.32 | 0.179 | 0.033 |
| R.319 Spleen MLR+ vs MLR- | HBA | 0.011 | 0.000 | 0.36 | 0.308 | 0.086 |
| R.319 Spleen MLR+ vs MLR- | RPL30 | 0.012 | 0.000 | -0.44 | 1 | 1 |
| R.319 Spleen MLR+ vs MLR- | RPL5 | 0.015 | 0.000 | -0.47 | 1 | 1 |
| R.319 Spleen MLR+ vs MLR- | S100A10 | 0.015 | 0.000 | 0.68 | 0.949 | 0.909 |
| R.319 Spleen MLR+ vs MLR- | RPS23 | 0.015 | 0.000 | -0.43 | 1 | 1 |
| R.319 Spleen MLR+ vs MLR- | MAMU-E | 0.016 | 0.000 | 0.43 | 1 | 0.999 |
| R.319 Spleen MLR+ vs MLR- | RPS7 | 0.018 | 0.000 | -0.39 | 1 | 1 |
| R.319 Spleen MLR+ vs MLR- | CD7 | 0.023 | 0.000 | -0.85 | 0.308 | 0.679 |
| R.319 Spleen MLR+ vs MLR- | ENSMMUG00000013429 | 0.028 | 0.000 | -0.48 | 1 | 1 |
| R.319 Spleen MLR+ vs MLR- | HARS2 | 0.031 | 0.000 | 0.26 | 0.256 | 0.066 |
| R.319 Spleen MLR+ vs MLR- | ELOA | 0.035 | 0.000 | -0.40 | 1 | 1 |
| R.319 Spleen MLR+ vs MLR- | RPL18 | 0.037 | 0.000 | -0.41 | 1 | 1 |
| R.319 Spleen MLR+ vs MLR- | RPS16 | 0.039 | 0.000 | -0.41 | 1 | 1 |
| R.319 Spleen MLR+ vs MLR- | CD63 | 0.047 | 0.000 | 0.70 | 0.615 | 0.33 |
| R.319 Spleen MLR+ vs MLR- | EFHD2 | 0.047 | 0.000 | 0.59 | 0.718 | 0.403 |
| R.319 Spleen MLR+ vs MLR- | RPL37A | 0.053 | 0.000 | -0.45 | 1 | 0.998 |
| R.319 Spleen MLR+ vs MLR- | RPS26 | 0.056 | 0.000 | -0.35 | 1 | 1 |
| R.319 Spleen MLR+ vs MLR- | CD74 | 0.056 | 0.000 | 0.67 | 0.641 | 0.322 |
| R.319 Spleen MLR+ vs MLR- | ENSMMUG00000003867 | 0.057 | 0.000 | -0.44 | 1 | 1 |
| R.319 Spleen MLR+ vs MLR- | ENSMMUG00000062350 | 0.063 | 0.000 | -0.38 | 1 | 1 |
| R.319 Spleen MLR+ vs MLR- | CPD | 0.066 | 0.000 | 0.40 | 0.436 | 0.168 |
| R.319 Spleen MLR+ vs MLR- | RPS24 | 0.072 | 0.000 | -0.45 | 1 | 0.999 |
| R.319 Spleen MLR+ vs MLR- | RPS13 | 0.073 | 0.000 | -0.38 | 1 | 1 |
| R.319 Spleen MLR+ vs MLR- | PPP1CC | 0.078 | 0.000 | -0.79 | 0.538 | 0.756 |
| R.319 Spleen MLR+ vs MLR- | RPSA | 0.138 | 0.000 | -0.39 | 1 | 1 |
| R.319 Spleen MLR+ vs MLR- | GPR183 | 0.153 | 0.000 | -0.88 | 0.282 | 0.646 |
| R.319 Spleen MLR+ vs MLR- | RPS5 | 0.160 | 0.000 | -0.37 | 1 | 1 |
| R.319 Spleen MLR+ vs MLR- | NACA | 0.161 | 0.000 | -0.44 | 1 | 0.999 |
| R.319 Spleen MLR+ vs MLR- | TPT1 | 0.176 | 0.000 | -0.51 | 1 | 1 |
| R.319 Spleen MLR+ vs MLR- | ENSMMUG00000063637 | 0.203 | 0.000 | -0.41 | 1 | 1 |
| R.319 Spleen MLR+ vs MLR- | RPLP1 | 0.241 | 0.000 | -0.32 | 1 | 1 |

| | | | | | | |
|---|---|---|---|---|---|---|
| R.319 Spleen MLR+ vs MLR- | RPS12 | 0.286 | 0.000 | -0.37 | 1 | 1 |
| R.319 Spleen MLR+ vs MLR- | RPS21 | 0.317 | 0.000 | -0.39 | 1 | 1 |
| R.319 Spleen MLR+ vs MLR- | S100A4 | 0.345 | 0.000 | 0.57 | 0.974 | 0.695 |
| R.319 Spleen MLR+ vs MLR- | RPS28 | 0.374 | 0.000 | -0.32 | 1 | 1 |
| R.319 Spleen MLR+ vs MLR- | LEF1 | 0.432 | 0.000 | -0.64 | 0.077 | 0.406 |
| R.319 Spleen MLR+ vs MLR- | DDAH2 | 0.513 | 0.000 | 0.27 | 0.179 | 0.044 |
| R.319 Spleen MLR+ vs MLR- | ENSMMUG00000054038 | 0.518 | 0.000 | 0.41 | 1 | 0.988 |
| R.319 Spleen MLR+ vs MLR- | CD84 | 0.639 | 0.000 | 0.31 | 0.359 | 0.136 |
| R.319 Spleen MLR+ vs MLR- | NKG2D | 0.722 | 0.000 | 0.35 | 0.333 | 0.119 |
| R.319 Spleen MLR+ vs MLR- | CASP12 | 0.838 | 0.000 | -0.61 | 0.462 | 0.738 |
| R.319 Spleen MLR+ vs MLR- | SERPINA1 | 0.857 | 0.000 | 0.42 | 0.59 | 0.287 |
| R.319 Spleen MLR+ vs MLR- | RPL32 | 1.000 | 0.000 | -0.35 | 1 | 1 |
| R.319 Spleen MLR+ vs MLR- | COX2 | 1.000 | 0.000 | -0.45 | 1 | 1 |
| R.319 Spleen MLR+ vs MLR- | THY1 | 1.000 | 0.000 | 0.51 | 0.462 | 0.209 |
| R.319 Spleen MLR+ vs MLR- | RNF103 | 1.000 | 0.000 | 0.29 | 0.333 | 0.126 |
| R.319 Spleen MLR+ vs MLR- | COX1 | 1.000 | 0.000 | -0.52 | 0.949 | 0.979 |
| R.319 Spleen MLR+ vs MLR- | RPL10A | 1.000 | 0.000 | -0.43 | 1 | 0.999 |
| R.319 Spleen MLR+ vs MLR- | CMC1 | 1.000 | 0.000 | 0.49 | 0.436 | 0.204 |
| R.319 Spleen MLR+ vs MLR- | RPL23A | 1.000 | 0.000 | -0.51 | 0.667 | 0.88 |
| R.319 Spleen MLR+ vs MLR- | CXCR4 | 1.000 | 0.000 | 0.68 | 0.846 | 0.643 |
| R.319 Spleen MLR+ vs MLR- | RPAP3 | 1.000 | 0.000 | 0.27 | 0.205 | 0.06 |
| R.319 Spleen MLR+ vs MLR- | MAP4K1 | 1.000 | 0.000 | 0.40 | 0.59 | 0.322 |
| R.319 Spleen MLR+ vs MLR- | DDX5 | 1.000 | 0.000 | 0.37 | 0.949 | 0.958 |
| R.319 Spleen MLR+ vs MLR- | RPS27A.1 | 1.000 | 0.000 | -0.35 | 1 | 1 |
| R.319 Spleen MLR+ vs MLR- | CRTAM | 1.000 | 0.000 | 0.36 | 0.282 | 0.103 |
| R.319 Spleen MLR+ vs MLR- | RPL6 | 1.000 | 0.000 | -0.31 | 1 | 1 |
| R.319 Spleen MLR+ vs MLR- | PPP1R12A | 1.000 | 0.000 | 0.31 | 0.59 | 0.312 |
| R.319 Spleen MLR+ vs MLR- | RPL24 | 1.000 | 0.000 | -0.33 | 1 | 1 |
| R.319 Spleen MLR+ vs MLR- | RPL22 | 1.000 | 0.000 | -0.35 | 1 | 0.999 |
| R.319 Spleen MLR+ vs MLR- | DHRS7 | 1.000 | 0.000 | 0.37 | 0.487 | 0.24 |
| R.319 Spleen MLR+ vs MLR- | CD28 | 1.000 | 0.000 | -0.47 | 0.077 | 0.349 |
| R.319 Spleen MLR+ vs MLR- | BST2 | 1.000 | 0.000 | 0.52 | 0.821 | 0.614 |
| R.319 Spleen MLR+ vs MLR- | UBB | 1.000 | 0.000 | 0.42 | 0.949 | 0.932 |
| R.319 Spleen MLR+ vs MLR- | ITGB2 | 1.000 | 0.000 | 0.49 | 0.795 | 0.637 |
| R.319 Spleen MLR+ vs MLR- | RSRP1 | 1.000 | 0.000 | 0.49 | 0.641 | 0.382 |
| R.319 Spleen MLR+ vs MLR- | CAPG | 1.000 | 0.000 | -0.67 | 0.513 | 0.708 |
| R.319 Spleen MLR+ vs MLR- | ENSMMUG00000004441 | 1.000 | 0.000 | 0.42 | 0.974 | 0.93 |
| R.319 Spleen MLR+ vs MLR- | DGKA | 1.000 | 0.001 | -0.46 | 0.179 | 0.463 |
| R.319 Spleen MLR+ vs MLR- | RPS3 | 1.000 | 0.001 | -0.32 | 1 | 0.998 |
| R.319 Spleen MLR+ vs MLR- | NUDT3 | 1.000 | 0.001 | -0.29 | 1 | 1 |
| R.319 Spleen MLR+ vs MLR- | NMI | 1.000 | 0.001 | 0.34 | 0.41 | 0.195 |
| R.319 Spleen MLR+ vs MLR- | SLC2A3 | 1.000 | 0.001 | -0.65 | 0.359 | 0.621 |
| R.319 Spleen MLR+ vs MLR- | RPS15 | 1.000 | 0.001 | -0.28 | 1 | 1 |
| R.319 Spleen MLR+ vs MLR- | SSBP4 | 1.000 | 0.001 | 0.45 | 0.487 | 0.274 |
| R.319 Spleen MLR+ vs MLR- | PRF1 | 1.000 | 0.001 | 0.38 | 0.41 | 0.19 |
| R.319 Spleen MLR+ vs MLR- | PXN | 1.000 | 0.001 | 0.33 | 0.385 | 0.182 |
| R.319 Spleen MLR+ vs MLR- | FCMR | 1.000 | 0.001 | -0.44 | 0.103 | 0.373 |
| R.319 Spleen MLR+ vs MLR- | RPL13A | 1.000 | 0.001 | -0.29 | 1 | 1 |
| R.319 Spleen MLR+ vs MLR- | RPL38 | 1.000 | 0.001 | -0.30 | 1 | 1 |
| R.319 Spleen MLR+ vs MLR- | NBEAL2 | 1.000 | 0.001 | 0.30 | 0.308 | 0.128 |
| R.319 Spleen MLR+ vs MLR- | ICOS | 1.000 | 0.001 | -0.52 | 0.154 | 0.414 |
| R.319 Spleen MLR+ vs MLR- | RPS14 | 1.000 | 0.001 | -0.30 | 1 | 1 |
| R.319 Spleen MLR+ vs MLR- | LAMP1 | 1.000 | 0.001 | 0.40 | 0.641 | 0.421 |
| R.319 Spleen MLR+ vs MLR- | RPL35A | 1.000 | 0.001 | -0.27 | 1 | 1 |
| R.319 Spleen MLR+ vs MLR- | SERTAD2 | 1.000 | 0.001 | -0.34 | 0 | 0.224 |
| R.319 Spleen MLR+ vs MLR- | ST8SIA4 | 1.000 | 0.001 | 0.29 | 0.436 | 0.215 |

| | | | | | | |
|---|---|---|---|---|---|---|
| R.319 Spleen MLR+ vs MLR- | ANP32B | 1.000 | 0.001 | -0.45 | 0.333 | 0.608 |
| R.319 Spleen MLR+ vs MLR- | TIGIT | 1.000 | 0.001 | 0.28 | 0.513 | 0.268 |
| R.319 Spleen MLR+ vs MLR- | PECAM1 | 1.000 | 0.001 | -0.41 | 0.051 | 0.292 |
| R.319 Spleen MLR+ vs MLR- | DGKZ | 1.000 | 0.001 | 0.43 | 0.59 | 0.385 |
| R.319 Spleen MLR+ vs MLR- | HNRNPA1 | 1.000 | 0.001 | -0.37 | 0.974 | 0.963 |
| R.319 Spleen MLR+ vs MLR- | RGCC | 1.000 | 0.001 | -0.61 | 0.718 | 0.873 |
| R.319 Spleen MLR+ vs MLR- | EEF2 | 1.000 | 0.001 | -0.35 | 0.974 | 0.997 |
| R.319 Spleen MLR+ vs MLR- | ATP6AP2 | 1.000 | 0.001 | 0.36 | 0.462 | 0.253 |
| R.319 Spleen MLR+ vs MLR- | JUNB | 1.000 | 0.001 | -0.57 | 0.487 | 0.695 |
| R.319 Spleen MLR+ vs MLR- | ELF4 | 1.000 | 0.001 | 0.25 | 0.359 | 0.166 |
| R.319 Spleen MLR+ vs MLR- | CFP | 1.000 | 0.001 | -0.32 | 0.051 | 0.305 |
| R.319 Spleen MLR+ vs MLR- | CLEC2D | 1.000 | 0.001 | 0.36 | 0.795 | 0.592 |
| R.319 Spleen MLR+ vs MLR- | STK17A | 1.000 | 0.001 | -0.44 | 0.718 | 0.832 |
| R.319 Spleen MLR+ vs MLR- | ENSMMUG00000013256 | 1.000 | 0.001 | -0.38 | 0.41 | 0.69 |
| R.319 Spleen MLR+ vs MLR- | ENSMMUG00000002320 | 1.000 | 0.001 | 0.43 | 1 | 0.997 |
| R.319 Spleen MLR+ vs MLR- | COTL1 | 1.000 | 0.001 | -0.57 | 0.564 | 0.808 |
| R.319 Spleen MLR+ vs MLR- | ZBTB38 | 1.000 | 0.001 | 0.25 | 0.282 | 0.119 |
| R.319 Spleen MLR+ vs MLR- | LAPTM5 | 1.000 | 0.001 | -0.47 | 0.923 | 0.949 |
| R.319 Spleen MLR+ vs MLR- | RPL13 | 1.000 | 0.002 | -0.27 | 1 | 1 |
| R.319 Spleen MLR+ vs MLR- | DCTN3 | 1.000 | 0.002 | -0.38 | 0.154 | 0.404 |
| R.319 Spleen MLR+ vs MLR- | RPL3 | 1.000 | 0.002 | -0.29 | 0.974 | 1 |
| R.319 Spleen MLR+ vs MLR- | UBA52 | 1.000 | 0.002 | -0.26 | 1 | 1 |
| R.319 Spleen MLR+ vs MLR- | INPP5D | 1.000 | 0.002 | 0.26 | 0.359 | 0.17 |
| R.319 Spleen MLR+ vs MLR- | CTSL | 1.000 | 0.002 | -0.41 | 0.051 | 0.28 |
| R.319 Spleen MLR+ vs MLR- | ENSMMUG00000022489 | 1.000 | 0.002 | -0.35 | 0.974 | 0.942 |
| R.319 Spleen MLR+ vs MLR- | RPL4 | 1.000 | 0.002 | -0.32 | 1 | 0.988 |
| R.319 Spleen MLR+ vs MLR- | TRIB2 | 1.000 | 0.002 | -0.34 | 0.026 | 0.236 |
| R.319 Spleen MLR+ vs MLR- | PLAC8 | 1.000 | 0.002 | -0.78 | 0.282 | 0.524 |
| R.319 Spleen MLR+ vs MLR- | H2AFZ | 1.000 | 0.002 | -0.59 | 0.462 | 0.69 |
| R.319 Spleen MLR+ vs MLR- | FYB1 | 1.000 | 0.002 | -0.41 | 0.462 | 0.741 |
| R.319 Spleen MLR+ vs MLR- | ITGA1 | 1.000 | 0.002 | -0.33 | 0 | 0.197 |
| R.319 Spleen MLR+ vs MLR- | CSNK2B | 1.000 | 0.002 | -0.36 | 0.282 | 0.545 |
| R.319 Spleen MLR+ vs MLR- | CTSC | 1.000 | 0.002 | 0.27 | 0.436 | 0.228 |
| R.319 Spleen MLR+ vs MLR- | EIF2S2 | 1.000 | 0.002 | -0.31 | 0.103 | 0.36 |
| R.319 Spleen MLR+ vs MLR- | PTPRC | 1.000 | 0.002 | 0.35 | 0.974 | 0.94 |
| R.319 Spleen MLR+ vs MLR- | TSC22D3 | 1.000 | 0.002 | 0.43 | 0.974 | 0.875 |
| R.319 Spleen MLR+ vs MLR- | IRF1 | 1.000 | 0.002 | 0.31 | 0.974 | 0.858 |
| R.319 Spleen MLR+ vs MLR- | REL | 1.000 | 0.003 | -0.40 | 0.205 | 0.446 |
| R.319 Spleen MLR+ vs MLR- | PDE4B | 1.000 | 0.003 | -0.42 | 0.154 | 0.384 |
| R.319 Spleen MLR+ vs MLR- | ATP5MC2 | 1.000 | 0.003 | -0.40 | 0.846 | 0.885 |
| R.319 Spleen MLR+ vs MLR- | CHD3 | 1.000 | 0.003 | -0.36 | 0.154 | 0.398 |
| R.319 Spleen MLR+ vs MLR- | HNRNPA2B1 | 1.000 | 0.003 | 0.27 | 0.974 | 0.894 |
| R.319 Spleen MLR+ vs MLR- | CNBP | 1.000 | 0.003 | 0.36 | 0.821 | 0.745 |
| R.319 Spleen MLR+ vs MLR- | ATP5F1D | 1.000 | 0.003 | -0.39 | 0.615 | 0.808 |
| R.319 Spleen MLR+ vs MLR- | FRG1 | 1.000 | 0.003 | -0.31 | 0.051 | 0.261 |
| R.319 Spleen MLR+ vs MLR- | DNAJA1 | 1.000 | 0.003 | 0.52 | 0.744 | 0.549 |
| R.319 Spleen MLR+ vs MLR- | CLK1 | 1.000 | 0.003 | 0.32 | 0.538 | 0.332 |
| R.319 Spleen MLR+ vs MLR- | RPL9 | 1.000 | 0.003 | -0.29 | 0.974 | 0.999 |
| R.319 Spleen MLR+ vs MLR- | ABRACL | 1.000 | 0.003 | -0.40 | 0.385 | 0.66 |
| R.319 Spleen MLR+ vs MLR- | SOCS3 | 1.000 | 0.003 | -0.40 | 0.179 | 0.406 |
| R.319 Spleen MLR+ vs MLR- | SMCHD1 | 1.000 | 0.003 | -0.34 | 0.128 | 0.365 |
| R.319 Spleen MLR+ vs MLR- | TCF7 | 1.000 | 0.003 | -0.49 | 0.308 | 0.526 |
| R.319 Spleen MLR+ vs MLR- | ENSMMUG00000064692 | 1.000 | 0.003 | -0.38 | 0.513 | 0.744 |
| R.319 Spleen MLR+ vs MLR- | EIF3B | 1.000 | 0.004 | -0.30 | 0.051 | 0.252 |
| R.319 Spleen MLR+ vs MLR- | RPL39 | 1.000 | 0.004 | -0.25 | 1 | 1 |
| R.319 Spleen MLR+ vs MLR- | ARHGAP9 | 1.000 | 0.004 | 0.41 | 0.538 | 0.35 |

| | | | | | | |
|---|---|---|---|---|---|---|
| R.319 Spleen MLR+ vs MLR- | SELL | 1.000 | 0.004 | -0.58 | 0.256 | 0.484 |
| R.319 Spleen MLR+ vs MLR- | RPS6KA1 | 1.000 | 0.004 | 0.33 | 0.41 | 0.227 |
| R.319 Spleen MLR+ vs MLR- | EEF1B2 | 1.000 | 0.005 | -0.35 | 0.949 | 0.983 |
| R.319 Spleen MLR+ vs MLR- | EIF3E | 1.000 | 0.005 | -0.39 | 0.513 | 0.712 |
| R.319 Spleen MLR+ vs MLR- | CD38 | 1.000 | 0.005 | -0.48 | 0.154 | 0.353 |
| R.319 Spleen MLR+ vs MLR- | VAMP5 | 1.000 | 0.005 | -0.32 | 0.103 | 0.307 |
| R.319 Spleen MLR+ vs MLR- | FKBP8 | 1.000 | 0.005 | 0.28 | 0.667 | 0.459 |
| R.319 Spleen MLR+ vs MLR- | DECR2 | 1.000 | 0.005 | -0.26 | 0.128 | 0.369 |
| R.319 Spleen MLR+ vs MLR- | PGD | 1.000 | 0.005 | -0.27 | 0.026 | 0.209 |
| R.319 Spleen MLR+ vs MLR- | CD5 | 1.000 | 0.005 | -0.37 | 0.179 | 0.403 |
| R.319 Spleen MLR+ vs MLR- | IL2RB | 1.000 | 0.005 | 0.50 | 0.59 | 0.393 |
| R.319 Spleen MLR+ vs MLR- | ZFP36 | 1.000 | 0.005 | -0.45 | 0.872 | 0.938 |
| R.319 Spleen MLR+ vs MLR- | HSD17B4 | 1.000 | 0.005 | -0.32 | 0.103 | 0.314 |
| R.319 Spleen MLR+ vs MLR- | RPL23 | 1.000 | 0.005 | -0.33 | 0.872 | 0.918 |
| R.319 Spleen MLR+ vs MLR- | SLC25A6 | 1.000 | 0.005 | -0.32 | 0.897 | 0.977 |
| R.319 Spleen MLR+ vs MLR- | ENSMMUG00000017097 | 1.000 | 0.005 | 0.28 | 0.41 | 0.231 |
| R.319 Spleen MLR+ vs MLR- | CDKN1A | 1.000 | 0.006 | -0.55 | 0.128 | 0.327 |
| R.319 Spleen MLR+ vs MLR- | SFRP5 | 1.000 | 0.006 | -0.34 | 0 | 0.166 |
| R.319 Spleen MLR+ vs MLR- | ENSMMUG00000020050 | 1.000 | 0.006 | -0.29 | 0.179 | 0.395 |
| R.319 Spleen MLR+ vs MLR- | PPM1J | 1.000 | 0.006 | 0.34 | 0.308 | 0.155 |
| R.319 Spleen MLR+ vs MLR- | FNTB | 1.000 | 0.006 | -0.32 | 0.179 | 0.403 |
| R.319 Spleen MLR+ vs MLR- | CYB5A | 1.000 | 0.006 | -0.33 | 0.154 | 0.374 |
| R.319 Spleen MLR+ vs MLR- | COX5A | 1.000 | 0.007 | -0.37 | 0.487 | 0.661 |
| R.319 Spleen MLR+ vs MLR- | RABAC1 | 1.000 | 0.007 | 0.33 | 0.615 | 0.424 |
| R.319 Spleen MLR+ vs MLR- | ITGB7 | 1.000 | 0.007 | -0.30 | 0.256 | 0.503 |
| R.319 Spleen MLR+ vs MLR- | RPL7 | 1.000 | 0.007 | -0.29 | 0.897 | 0.953 |
| R.319 Spleen MLR+ vs MLR- | HSPA8 | 1.000 | 0.007 | 0.28 | 1 | 0.955 |
| R.319 Spleen MLR+ vs MLR- | MRPS34 | 1.000 | 0.007 | -0.30 | 0.154 | 0.363 |
| R.319 Spleen MLR+ vs MLR- | ENSMMUG00000064873 | 1.000 | 0.007 | -0.29 | 0.179 | 0.39 |
| R.319 Spleen MLR+ vs MLR- | CD3G | 1.000 | 0.007 | 0.33 | 0.897 | 0.798 |
| R.319 Spleen MLR+ vs MLR- | CLIC1 | 1.000 | 0.007 | 0.31 | 0.872 | 0.745 |
| R.319 Spleen MLR+ vs MLR- | PLK2 | 1.000 | 0.007 | -0.31 | 0 | 0.157 |
| R.319 Spleen MLR+ vs MLR- | FAM102A | 1.000 | 0.007 | -0.26 | 0.026 | 0.196 |
| R.319 Spleen MLR+ vs MLR- | SLC25A43 | 1.000 | 0.008 | -0.34 | 0.385 | 0.67 |
| R.319 Spleen MLR+ vs MLR- | FLOT1 | 1.000 | 0.008 | -0.34 | 0.205 | 0.42 |
| R.319 Spleen MLR+ vs MLR- | PAG1 | 1.000 | 0.008 | -0.30 | 0.077 | 0.263 |
| R.319 Spleen MLR+ vs MLR- | UBE2L6 | 1.000 | 0.008 | -0.27 | 0.385 | 0.606 |
| R.319 Spleen MLR+ vs MLR- | HNRNPH2 | 1.000 | 0.008 | -0.29 | 1 | 0.985 |
| R.319 Spleen MLR+ vs MLR- | ENSMMUG00000059937 | 1.000 | 0.008 | -0.38 | 0.077 | 0.267 |
| R.319 Spleen MLR+ vs MLR- | STMN1 | 1.000 | 0.008 | -0.72 | 0.103 | 0.297 |
| R.319 Spleen MLR+ vs MLR- | TMEM50A | 1.000 | 0.009 | 0.32 | 0.59 | 0.424 |
| R.319 Spleen MLR+ vs MLR- | EIF3F | 1.000 | 0.009 | -0.32 | 0.615 | 0.865 |
| R.319 Spleen MLR+ vs MLR- | NDUFAB1 | 1.000 | 0.009 | -0.29 | 0.103 | 0.289 |
| R.319 Spleen MLR+ vs MLR- | LGALS3 | 1.000 | 0.009 | 0.39 | 0.615 | 0.437 |
| R.319 Spleen MLR+ vs MLR- | IL4R | 1.000 | 0.009 | -0.31 | 0.103 | 0.294 |
| R.319 Spleen MLR+ vs MLR- | RNF138 | 1.000 | 0.010 | -0.26 | 0.051 | 0.223 |
| R.319 Spleen MLR+ vs MLR- | LIMD2 | 1.000 | 0.010 | -0.34 | 0.487 | 0.729 |
| R.319 Spleen MLR+ vs MLR- | ATP5PO | 1.000 | 0.011 | -0.26 | 0.744 | 0.863 |
| R.319 Spleen MLR+ vs MLR- | VIM | 1.000 | 0.011 | -0.41 | 0.974 | 0.975 |
| R.319 Spleen MLR+ vs MLR- | SPRYD3 | 1.000 | 0.012 | 0.28 | 0.282 | 0.147 |
| R.319 Spleen MLR+ vs MLR- | PPP3CC | 1.000 | 0.012 | 0.30 | 0.436 | 0.276 |
| R.319 Spleen MLR+ vs MLR- | R3HDM4 | 1.000 | 0.012 | -0.27 | 0.154 | 0.355 |
| R.319 Spleen MLR+ vs MLR- | RANBP1 | 1.000 | 0.012 | -0.32 | 0.128 | 0.307 |
| R.319 Spleen MLR+ vs MLR- | COPRS | 1.000 | 0.012 | -0.32 | 0.128 | 0.306 |
| R.319 Spleen MLR+ vs MLR- | PARP8 | 1.000 | 0.012 | 0.31 | 0.333 | 0.184 |
| R.319 Spleen MLR+ vs MLR- | NPM1 | 1.000 | 0.012 | -0.28 | 0.923 | 0.975 |

| | | | | | | |
|---|---|---|---|---|---|---|
| R.319 Spleen MLR+ vs MLR- | CUTA | 1.000 | 0.013 | -0.31 | 0.179 | 0.367 |
| R.319 Spleen MLR+ vs MLR- | NFKB2 | 1.000 | 0.014 | -0.25 | 0.051 | 0.212 |
| R.319 Spleen MLR+ vs MLR- | SYAP1 | 1.000 | 0.014 | -0.28 | 0.103 | 0.276 |
| R.319 Spleen MLR+ vs MLR- | NDUFS8 | 1.000 | 0.014 | -0.30 | 0.205 | 0.393 |
| R.319 Spleen MLR+ vs MLR- | EIF3M | 1.000 | 0.014 | -0.32 | 0.359 | 0.576 |
| R.319 Spleen MLR+ vs MLR- | ORAI1 | 1.000 | 0.014 | -0.28 | 0.231 | 0.428 |
| R.319 Spleen MLR+ vs MLR- | S1PR1 | 1.000 | 0.014 | -0.28 | 0.179 | 0.391 |
| R.319 Spleen MLR+ vs MLR- | HERPUD1 | 1.000 | 0.014 | 0.42 | 0.641 | 0.506 |
| R.319 Spleen MLR+ vs MLR- | NUTF2 | 1.000 | 0.014 | -0.30 | 0.205 | 0.383 |
| R.319 Spleen MLR+ vs MLR- | SERTAD1 | 1.000 | 0.014 | 0.40 | 0.692 | 0.585 |
| R.319 Spleen MLR+ vs MLR- | DUSP2 | 1.000 | 0.014 | -0.59 | 0.513 | 0.656 |
| R.319 Spleen MLR+ vs MLR- | PNISR | 1.000 | 0.015 | 0.35 | 0.513 | 0.386 |
| R.319 Spleen MLR+ vs MLR- | PBXIP1 | 1.000 | 0.015 | -0.29 | 0.256 | 0.459 |
| R.319 Spleen MLR+ vs MLR- | NUDC | 1.000 | 0.015 | 0.36 | 0.564 | 0.405 |
| R.319 Spleen MLR+ vs MLR- | DRAP1 | 1.000 | 0.015 | 0.35 | 0.564 | 0.44 |
| R.319 Spleen MLR+ vs MLR- | PTP4A1 | 1.000 | 0.015 | -0.31 | 0.154 | 0.336 |
| R.319 Spleen MLR+ vs MLR- | LIMS1 | 1.000 | 0.015 | -0.36 | 0.231 | 0.403 |
| R.319 Spleen MLR+ vs MLR- | CYBA | 1.000 | 0.015 | 0.32 | 0.821 | 0.765 |
| R.319 Spleen MLR+ vs MLR- | SLCO3A1 | 1.000 | 0.016 | 0.27 | 0.179 | 0.078 |
| R.319 Spleen MLR+ vs MLR- | SERPINB9 | 1.000 | 0.016 | 0.35 | 0.513 | 0.343 |
| R.319 Spleen MLR+ vs MLR- | GPSM3 | 1.000 | 0.016 | -0.34 | 0.538 | 0.726 |
| R.319 Spleen MLR+ vs MLR- | HM13 | 1.000 | 0.016 | -0.30 | 0.179 | 0.354 |
| R.319 Spleen MLR+ vs MLR- | TMEM243 | 1.000 | 0.016 | -0.27 | 0.103 | 0.271 |
| R.319 Spleen MLR+ vs MLR- | RPL35 | 1.000 | 0.016 | -0.26 | 1 | 0.987 |
| R.319 Spleen MLR+ vs MLR- | LMNA | 1.000 | 0.016 | -0.43 | 0.282 | 0.495 |
| R.319 Spleen MLR+ vs MLR- | NUFIP2 | 1.000 | 0.017 | -0.26 | 0.103 | 0.277 |
| R.319 Spleen MLR+ vs MLR- | NDUFV2 | 1.000 | 0.017 | -0.31 | 0.256 | 0.455 |
| R.319 Spleen MLR+ vs MLR- | RHOG | 1.000 | 0.017 | -0.33 | 0.41 | 0.557 |
| R.319 Spleen MLR+ vs MLR- | RGS10 | 1.000 | 0.018 | -0.27 | 0.103 | 0.267 |
| R.319 Spleen MLR+ vs MLR- | ND4L | 1.000 | 0.018 | -0.33 | 0.846 | 0.978 |
| R.319 Spleen MLR+ vs MLR- | SELENOK | 1.000 | 0.018 | 0.25 | 0.538 | 0.368 |
| R.319 Spleen MLR+ vs MLR- | GADD45B | 1.000 | 0.018 | 0.31 | 0.718 | 0.583 |
| R.319 Spleen MLR+ vs MLR- | DNAJC19 | 1.000 | 0.019 | 0.26 | 0.385 | 0.239 |
| R.319 Spleen MLR+ vs MLR- | CD53 | 1.000 | 0.020 | 0.30 | 0.769 | 0.631 |
| R.319 Spleen MLR+ vs MLR- | AHI1 | 1.000 | 0.020 | -0.29 | 0.179 | 0.368 |
| R.319 Spleen MLR+ vs MLR- | INPP4B | 1.000 | 0.021 | -0.27 | 0.154 | 0.341 |
| R.319 Spleen MLR+ vs MLR- | PABPC1 | 1.000 | 0.022 | -0.29 | 0.897 | 0.955 |
| R.319 Spleen MLR+ vs MLR- | APOBEC3G | 1.000 | 0.022 | 0.32 | 0.41 | 0.266 |
| R.319 Spleen MLR+ vs MLR- | PTPN22 | 1.000 | 0.023 | 0.27 | 0.333 | 0.193 |
| R.319 Spleen MLR+ vs MLR- | HINT1 | 1.000 | 0.023 | -0.26 | 0.974 | 0.941 |
| R.319 Spleen MLR+ vs MLR- | CHMP2A | 1.000 | 0.024 | -0.25 | 0.128 | 0.288 |
| R.319 Spleen MLR+ vs MLR- | GSTP1 | 1.000 | 0.024 | 0.26 | 0.923 | 0.885 |
| R.319 Spleen MLR+ vs MLR- | ENSMMUG00000018740 | 1.000 | 0.025 | 0.28 | 0.641 | 0.589 |
| R.319 Spleen MLR+ vs MLR- | CD6 | 1.000 | 0.025 | -0.32 | 0.282 | 0.445 |
| R.319 Spleen MLR+ vs MLR- | CD164 | 1.000 | 0.025 | -0.32 | 0.462 | 0.629 |
| R.319 Spleen MLR+ vs MLR- | PRR13 | 1.000 | 0.026 | 0.28 | 0.744 | 0.681 |
| R.319 Spleen MLR+ vs MLR- | CD99 | 1.000 | 0.027 | 0.40 | 0.641 | 0.537 |
| R.319 Spleen MLR+ vs MLR- | WDR83OS | 1.000 | 0.027 | -0.27 | 0.538 | 0.763 |
| R.319 Spleen MLR+ vs MLR- | CCT5 | 1.000 | 0.027 | -0.30 | 0.333 | 0.49 |
| R.319 Spleen MLR+ vs MLR- | MT1E | 1.000 | 0.028 | -0.42 | 0.179 | 0.343 |
| R.319 Spleen MLR+ vs MLR- | IK | 1.000 | 0.028 | 0.28 | 0.564 | 0.44 |
| R.319 Spleen MLR+ vs MLR- | ITGA6 | 1.000 | 0.028 | -0.25 | 0.103 | 0.251 |
| R.319 Spleen MLR+ vs MLR- | CA6 | 1.000 | 0.029 | -0.36 | 0.077 | 0.215 |
| R.319 Spleen MLR+ vs MLR- | ENO1 | 1.000 | 0.029 | -0.39 | 0.564 | 0.758 |
| R.319 Spleen MLR+ vs MLR- | KLF10 | 1.000 | 0.029 | -0.32 | 0.154 | 0.304 |
| R.319 Spleen MLR+ vs MLR- | CTLA4 | 1.000 | 0.029 | -0.30 | 0.128 | 0.284 |

| | | | | | | |
|---|---|---|---|---|---|---|
| R.319 Spleen MLR+ vs MLR- | TNFSF12 | 1.000 | 0.030 | 0.26 | 0.359 | 0.23 |
| R.319 Spleen MLR+ vs MLR- | ERGIC3 | 1.000 | 0.030 | -0.27 | 0.231 | 0.384 |
| R.319 Spleen MLR+ vs MLR- | NUCB2 | 1.000 | 0.030 | -0.28 | 0.231 | 0.393 |
| R.319 Spleen MLR+ vs MLR- | CAST | 1.000 | 0.031 | 0.27 | 0.564 | 0.391 |
| R.319 Spleen MLR+ vs MLR- | AHNAK | 1.000 | 0.031 | 0.33 | 0.667 | 0.554 |
| R.319 Spleen MLR+ vs MLR- | ENSMMUG00000063316 | 1.000 | 0.031 | -0.48 | 0.487 | 0.637 |
| R.319 Spleen MLR+ vs MLR- | ITGB1 | 1.000 | 0.031 | -0.49 | 0.385 | 0.547 |
| R.319 Spleen MLR+ vs MLR- | EIF3H | 1.000 | 0.032 | -0.25 | 0.667 | 0.788 |
| R.319 Spleen MLR+ vs MLR- | ALDOA | 1.000 | 0.032 | -0.30 | 0.821 | 0.925 |
| R.319 Spleen MLR+ vs MLR- | ENSMMUG00000053146 | 1.000 | 0.033 | -0.26 | 0.256 | 0.41 |
| R.319 Spleen MLR+ vs MLR- | ENSMMUG00000055584 | 1.000 | 0.034 | -0.27 | 0.462 | 0.625 |
| R.319 Spleen MLR+ vs MLR- | STX11 | 1.000 | 0.034 | 0.35 | 0.333 | 0.207 |
| R.319 Spleen MLR+ vs MLR- | CCL3 | 1.000 | 0.034 | 0.38 | 0.154 | 0.069 |
| R.319 Spleen MLR+ vs MLR- | CYLD | 1.000 | 0.034 | -0.27 | 0.231 | 0.393 |
| R.319 Spleen MLR+ vs MLR- | PLK3 | 1.000 | 0.035 | -0.30 | 0.256 | 0.404 |
| R.319 Spleen MLR+ vs MLR- | ANXA1 | 1.000 | 0.036 | 0.36 | 0.795 | 0.681 |
| R.319 Spleen MLR+ vs MLR- | PRDX1 | 1.000 | 0.037 | -0.34 | 0.436 | 0.602 |
| R.319 Spleen MLR+ vs MLR- | ENSMMUG00000043332 | 1.000 | 0.037 | -0.26 | 0.692 | 0.813 |
| R.319 Spleen MLR+ vs MLR- | PSMA6 | 1.000 | 0.037 | 0.30 | 0.564 | 0.433 |
| R.319 Spleen MLR+ vs MLR- | C3H7orf50 | 1.000 | 0.038 | 0.36 | 0.641 | 0.571 |
| R.319 Spleen MLR+ vs MLR- | NFATC1 | 1.000 | 0.039 | -0.27 | 0.128 | 0.274 |
| R.319 Spleen MLR+ vs MLR- | ENSMMUG00000003854 | 1.000 | 0.039 | 0.27 | 0.59 | 0.468 |
| R.319 Spleen MLR+ vs MLR- | SSH2 | 1.000 | 0.039 | -0.25 | 0.154 | 0.298 |
| R.319 Spleen MLR+ vs MLR- | CALR | 1.000 | 0.040 | 0.29 | 0.897 | 0.848 |
| R.319 Spleen MLR+ vs MLR- | TMSB10 | 1.000 | 0.040 | -0.30 | 1 | 1 |
| R.319 Spleen MLR+ vs MLR- | IGFLR1 | 1.000 | 0.040 | -0.27 | 0.205 | 0.354 |
| R.319 Spleen MLR+ vs MLR- | SON | 1.000 | 0.040 | 0.25 | 0.821 | 0.736 |
| R.319 Spleen MLR+ vs MLR- | APBB1IP | 1.000 | 0.041 | 0.29 | 0.59 | 0.47 |
| R.319 Spleen MLR+ vs MLR- | SEPTIN6 | 1.000 | 0.042 | -0.31 | 0.436 | 0.559 |
| R.319 Spleen MLR+ vs MLR- | RHEB | 1.000 | 0.043 | -0.27 | 0.231 | 0.384 |
| R.319 Spleen MLR+ vs MLR- | PFKL | 1.000 | 0.044 | -0.27 | 0.256 | 0.4 |
| R.319 Spleen MLR+ vs MLR- | TBC1D10B | 1.000 | 0.045 | -0.28 | 0.564 | 0.745 |
| R.319 Spleen MLR+ vs MLR- | TNFSF10 | 1.000 | 0.046 | -0.29 | 0.103 | 0.231 |
| R.319 Spleen MLR+ vs MLR- | TAPBPL | 1.000 | 0.048 | 0.27 | 0.872 | 0.775 |
| R.319 Spleen MLR+ vs MLR- | MVP | 1.000 | 0.048 | -0.27 | 0.359 | 0.5 |
| R.319 Spleen MLR+ vs MLR- | KRTCAP2 | 1.000 | 0.049 | -0.28 | 0.692 | 0.805 |
| R.319 Spleen MLR+ vs MLR- | ARID5B | 1.000 | 0.050 | -0.26 | 0.231 | 0.378 |
| R.319 Spleen MLR+ vs MLR- | SEPTIN7 | 1.000 | 0.051 | 0.26 | 0.641 | 0.554 |
| R.319 Spleen MLR+ vs MLR- | CYCS | 1.000 | 0.051 | -0.27 | 0.359 | 0.513 |
| R.319 Spleen MLR+ vs MLR- | IKZF3 | 1.000 | 0.052 | 0.29 | 0.333 | 0.214 |
| R.319 Spleen MLR+ vs MLR- | MAPRE2 | 1.000 | 0.052 | 0.29 | 0.538 | 0.408 |
| R.319 Spleen MLR+ vs MLR- | IL7R | 1.000 | 0.052 | -0.36 | 0.359 | 0.536 |
| R.319 Spleen MLR+ vs MLR- | IGFBP4 | 1.000 | 0.053 | -0.28 | 0.026 | 0.132 |
| R.319 Spleen MLR+ vs MLR- | RCSD1 | 1.000 | 0.053 | -0.26 | 0.308 | 0.454 |
| R.319 Spleen MLR+ vs MLR- | HNRNPDL | 1.000 | 0.054 | 0.25 | 0.923 | 0.88 |
| R.319 Spleen MLR+ vs MLR- | KLF2 | 1.000 | 0.056 | -0.32 | 0.923 | 0.917 |
| R.319 Spleen MLR+ vs MLR- | SRI | 1.000 | 0.056 | 0.27 | 0.59 | 0.461 |
| R.319 Spleen MLR+ vs MLR- | TIMP1 | 1.000 | 0.058 | -0.33 | 0.231 | 0.371 |
| R.319 Spleen MLR+ vs MLR- | ND5 | 1.000 | 0.060 | -0.27 | 0.436 | 0.553 |
| R.319 Spleen MLR+ vs MLR- | CKS2 | 1.000 | 0.062 | -0.31 | 0.231 | 0.365 |
| R.319 Spleen MLR+ vs MLR- | GTF2B | 1.000 | 0.063 | 0.29 | 0.692 | 0.553 |
| R.319 Spleen MLR+ vs MLR- | MAP4 | 1.000 | 0.064 | 0.27 | 0.359 | 0.248 |
| R.319 Spleen MLR+ vs MLR- | SKAP1 | 1.000 | 0.067 | 0.26 | 0.744 | 0.68 |
| R.319 Spleen MLR+ vs MLR- | ARL6IP5 | 1.000 | 0.067 | 0.30 | 0.59 | 0.525 |
| R.319 Spleen MLR+ vs MLR- | HSP90AA1 | 1.000 | 0.068 | 0.32 | 0.949 | 0.923 |
| R.319 Spleen MLR+ vs MLR- | PIM3 | 1.000 | 0.071 | -0.32 | 0.256 | 0.375 |

| | | | | | | |
|---|---|---|---|---|---|---|
| R.319 Spleen MLR+ vs MLR- | HSPA5 | 1.000 | 0.074 | 0.30 | 0.872 | 0.842 |
| R.319 Spleen MLR+ vs MLR- | CNN2 | 1.000 | 0.075 | -0.27 | 0.615 | 0.719 |
| R.319 Spleen MLR+ vs MLR- | DNAJB1 | 1.000 | 0.078 | 0.49 | 0.538 | 0.424 |
| R.319 Spleen MLR+ vs MLR- | COX8A | 1.000 | 0.078 | 0.28 | 0.718 | 0.646 |
| R.319 Spleen MLR+ vs MLR- | GHITM | 1.000 | 0.081 | 0.25 | 0.641 | 0.511 |
| R.319 Spleen MLR+ vs MLR- | RILPL2 | 1.000 | 0.082 | -0.26 | 0.231 | 0.351 |
| R.319 Spleen MLR+ vs MLR- | JAK1 | 1.000 | 0.083 | 0.32 | 0.692 | 0.612 |
| R.319 Spleen MLR+ vs MLR- | RBM39 | 1.000 | 0.083 | 0.28 | 0.821 | 0.733 |
| R.319 Spleen MLR+ vs MLR- | LRPAP1 | 1.000 | 0.087 | 0.29 | 0.333 | 0.226 |
| R.319 Spleen MLR+ vs MLR- | ARRB2 | 1.000 | 0.095 | 0.26 | 0.462 | 0.364 |
| R.319 Spleen MLR+ vs MLR- | ARHGAP4 | 1.000 | 0.095 | 0.26 | 0.513 | 0.427 |
| R.319 Spleen MLR+ vs MLR- | KIAA0040 | 1.000 | 0.096 | 0.28 | 0.308 | 0.213 |
| R.319 Spleen MLR+ vs MLR- | CYFIP2 | 1.000 | 0.097 | 0.30 | 0.41 | 0.326 |
| R.319 Spleen MLR+ vs MLR- | WIPF1 | 1.000 | 0.097 | 0.29 | 0.641 | 0.595 |
| R.319 Spleen MLR+ vs MLR- | ODC1 | 1.000 | 0.097 | 0.26 | 0.564 | 0.473 |
| R.319 Spleen MLR+ vs MLR- | TUBB | 1.000 | 0.097 | -0.54 | 0.513 | 0.63 |
| R.319 Spleen MLR+ vs MLR- | HMGB1 | 1.000 | 0.098 | -0.30 | 0.59 | 0.756 |
| R.319 Spleen MLR+ vs MLR- | ITM2A | 1.000 | 0.099 | -0.29 | 0.462 | 0.555 |
| R.319 Spleen MLR+ vs MLR- | COPE | 1.000 | 0.101 | 0.27 | 0.564 | 0.519 |
| R.319 Spleen MLR+ vs MLR- | HSPH1 | 1.000 | 0.114 | 0.41 | 0.462 | 0.411 |
| R.319 Spleen MLR+ vs MLR- | GADD45A | 1.000 | 0.119 | 0.33 | 0.282 | 0.197 |
| R.319 Spleen MLR+ vs MLR- | ND4 | 1.000 | 0.119 | -0.26 | 0.846 | 0.878 |
| R.319 Spleen MLR+ vs MLR- | USP47 | 1.000 | 0.125 | 0.29 | 0.282 | 0.207 |
| R.319 Spleen MLR+ vs MLR- | TOB1 | 1.000 | 0.126 | -0.28 | 0.538 | 0.63 |
| R.319 Spleen MLR+ vs MLR- | G3BP2 | 1.000 | 0.127 | -0.33 | 0.487 | 0.554 |
| R.319 Spleen MLR+ vs MLR- | TAF7 | 1.000 | 0.130 | 0.27 | 0.359 | 0.271 |
| R.319 Spleen MLR+ vs MLR- | RGS1 | 1.000 | 0.132 | 0.26 | 0.538 | 0.432 |
| R.319 Spleen MLR+ vs MLR- | ENSMMUG00000055690 | 1.000 | 0.133 | 0.42 | 1 | 1 |
| R.319 Spleen MLR+ vs MLR- | RSRC2 | 1.000 | 0.142 | 0.29 | 0.462 | 0.387 |
| R.319 Spleen MLR+ vs MLR- | SRSF7 | 1.000 | 0.150 | 0.35 | 0.615 | 0.624 |
| R.319 Spleen MLR+ vs MLR- | ID3 | 1.000 | 0.159 | -0.32 | 0.154 | 0.249 |
| R.319 Spleen MLR+ vs MLR- | MKI67 | 1.000 | 0.181 | -0.27 | 0.051 | 0.123 |
| R.319 Spleen MLR+ vs MLR- | FOSB | 1.000 | 0.211 | 0.38 | 0.718 | 0.794 |
| R.319 Spleen MLR+ vs MLR- | ENSMMUG00000051392 | 1.000 | 0.228 | -0.34 | 0.436 | 0.489 |
| R.319 Spleen MLR+ vs MLR- | JUN | 1.000 | 0.251 | 0.51 | 0.872 | 0.829 |
| R.319 Spleen MLR+ vs MLR- | NONO | 1.000 | 0.277 | 0.26 | 0.436 | 0.396 |
| R.319 Spleen MLR+ vs MLR- | DUSP1 | 1.000 | 0.287 | -0.26 | 0.846 | 0.849 |
| R.319 Spleen MLR+ vs MLR- | ZNF800 | 1.000 | 0.293 | 0.26 | 0.256 | 0.194 |
| R.319 Spleen MLR+ vs MLR- | ACTG1 | 1.000 | 0.322 | -0.28 | 1 | 0.995 |
| R.319 Spleen MLR+ vs MLR- | TNFSF9 | 1.000 | 0.323 | 0.29 | 0.179 | 0.129 |
| R.319 Spleen MLR+ vs MLR- | PSMA5 | 1.000 | 0.344 | 0.25 | 0.359 | 0.332 |
| R.319 Spleen MLR+ vs MLR- | GZMA | 1.000 | 0.371 | -0.40 | 0.179 | 0.127 |
| R.319 Spleen MLR+ vs MLR- | ENSMMUG00000062077 | 1.000 | 0.421 | 0.26 | 1 | 0.999 |
| R.319 Spleen MLR+ vs MLR- | DDIT3 | 1.000 | 0.511 | 0.25 | 0.205 | 0.174 |
| R.319 Spleen MLR+ vs MLR- | HMGB2 | 1.000 | 0.736 | -0.31 | 0.436 | 0.449 |
